# Minding The Gap: Range Size And Economic Use Drive Functional Trait Data Shortfall In The Atlantic Forest

**DOI:** 10.1101/2020.10.30.361311

**Authors:** Ana Carolina Petisco-Souza, Fernanda Thiesen Brum, Vinícius Marcilio-Silva, Victor P. Zwiener, Andressa Zanella, Arildo S. Dias, Andrés González-Melo, Steven Jansen, Guilherme G. Mazzochini, Ülo Niinemets, Valério D. Pillar, Enio Sosinski, Márcia C. M. Marques, Marcos B. Carlucci

## Abstract

Biodiversity shortfalls are knowledge gaps that may result from uneven sampling through time and space and human interest biases. Gaps in data of functional traits of species may add uncertainty in functional diversity and structure measures and hinder inference on ecosystem functioning and ecosystem services, with negative implications for conservation and restoration practices, such as in Atlantic Forest hotspot. Here we investigate which are the potential drivers of trait data gaps and where geographically they are in the Atlantic Forest. We quantified trait gaps for four key plant functional traits of 2335 trees species, and evaluated which factors drive trait gap at the species and at the geographical level. At the species level, we found larger trait gaps for small-ranged and with no economic use. At the geographical level, we found larger gaps at the Atlantic Forest east coast. Trait gaps were higher away from urban areas, and among species with smaller mean range size and smaller mean economic use of wood, and smaller near protected areas. Efforts on reducing trait gaps of small-ranged and of species with economic use of wood can further advance theory-driven studies and improve knowledge coverage

## BACKGROUND

Biodiversity shortfalls are gaps between currently known and complete knowledge of nature [1,2]. These knowledge gaps exist for different biological domains, such as taxonomy, species distribution, species abundances, evolutionary history, biotic interactions, biotic-abiotic interactions, species traits, and ecological functions [1,2], and can result from uneven sampling efforts at the species level, when little or nothing is known about features of a particular species [1,2]. Moreover, knowledge gaps may result from uneven sampling through time, or through geographical space [1]. The absence or poor quality of information at any scale could weaken the accuracy of biodiversity measures [1] which could set conservation standarts and, thereby, the robustness of arguments used to increase public awareness concerning the importance of conservation policies [3,4]. Hence, in order to minimize these gaps and set grounds for biodiversity research and conservation, it is necessary to identify the biodiversity knowledge that is missing, where it is missing more and why major gaps exist [3,4].

The ongoing mass extinction, driven by climate and land use changes urge us to understand biodiversity gaps [5,6], to prioritize ways of filling them, as the speed of such global changes could outpace the time to collect and analyze biodiversity data before it is lost. The lack of information regarding biodiversity lost by human intervention, could hinder the knowledge about ecosystem functioning [5,6]. Biodiversity might directly or indirectly mediate important ecosystem functions, such as nutrient cycling, water purification and primary productivity [7]. Moreover, ecosystem functions are important for human well-being, by providing (*e.g.* food, water and fiber), regulating (*e.g.* climate, pests and floods), supporting ecosystem services (*e.g.* soil formation, nutrient cycling and pollination), as well as delivering cultural value services (*e.g.* aesthetic, enjoyment and spiritual fulfillment) [8,9]. These ecosystem services are mediated by species functional traits, which are phenotypic, morphological or physiological features interact with other species features and with the environment, and affect species survival, growth and reproduction [10–13].

The variability of trait values among species within a community at the geographical scale can be correlated with ecosystem functions, such as biomass production [14,15], soil properties [16] and water consumption by plants [15]. Trait variability and mean can be estimated through functional diversity measures and related approaches [17,18]. At the community level, the contribution of dominant species can drive ecosystem functioning [14,15,19,20], as predicted by the biomass ratio hypothesis [19], which can be assessed by using the community weighted mean (CWM) approach [21]. Moreover, between-species differences in resource use (i.e. niche complementarity) may also drive ecosystem functioning [22,23] and it has been hypothesized that nearly complete resource use will enhance ecosystem functions, such as primary productivity [24], and can be assessed by means of functional dispersion measures. Trait dominance and dispersion can be used to clarify how biodiversity functionality is distributed throughout the geographical space [25,26] to predict ecosystem service *hotspots* [10,27]. Hence, it is possible to link functional diversity and structure measures to environmental changes, thereby guiding conservation priorities [28], and restoration actions [29]. The lack of information about functional traits may hamper BEF studies by producing uncertainties in measures based on functional traits, thereby leading to incomplete knowledge about how strong is the influence of biodiversity in mediating ecosystem functions, which in turn may impair conservation policies [2,3]. Thus, investigating which factors contribute to functional trait data gaps (hereafter trait gaps) is a fundamental step towards strengthening the accuracy and predictability of BEF approaches.

Trait gaps, also referred to as the Raunkiaeran biodiversity shortfall [1], may be caused by human interest, the lack of taxonomic research, and uneven spatial sampling. For instance, some species are better known as a consequence of their particular features of economic value. When the economic value of a species is known, such as the use of tree species for timber, trait information is more likely available [30]. There might also be a bias caused by research concentration in or near protected areas and nearby accessible sites, such as urban areas and highways [6,30–33]. Furthermore, evidence suggests that human interest in recording rare species might set aside widespread common species sampling [34], although there is also evidence that rare species with narrow ranges could be underrepresented [35]. Reducing such biases in biodiversity data collection is necessary to maximize knowledge about species ecology, including functional trait data that can be further integrated with the provisioning of ecosystem services. For this, investigating and understanding trait gaps is of paramount importance. One possible way to investigate trait gaps is to create “maps of ignorance” that incorporate spatially explicit estimates of data gaps, which can help to create agendas of spatial prioritization of sampling effort [1,36].

Megadiverse tropical countries might benefit from supporting investigations on which trait gaps exist and where functional diversity measures are uncertain. In this study, we chose to work with the Brazilian Atlantic Forest, a recognized biodiversity hotspot [37]. The Atlantic Forest has historically suffered from degradation and fragmentation [38], with potential loss of ecosystem services [39]. Therefore, understanding trait gaps is urgent to support conservation and restoration strategies for the Atlantic Forest [40]. In this study, we aimed to answer the following questions at different levels of analysis. At the species level: (1) what is the extent of trait gaps for tree species in the Brazilian Atlantic Forest? And (2) are trait gaps biased by the economic use of wood of species and the species range size? At the geographical level: (3) Are trait gaps explained by the economic use of wood, species range size, and distances from urban areas and protected areas across the Atlantic Forest? And (4) where do trait gaps cause high uncertainty in trait dominance and dispersion estimates across the Atlantic Forest?

## METHODS

### Species

In order to quantify trait gaps among Atlantic Forest trees we compiled data from 2335 species including woody plants, tree-like cacti and palms compiled by Zwiener et al. (2017) [41]. The 2335 species were selected from summarized checklists of 300 localities across the Atlantic forest obtained in field inventories and georeferenced data from the electronic database SpeciesLink [41].

We obtained the species range and range size from a previous study [41], which uses the potential geographical distribution generated by species distribution modeling (SDM) using the “MaxEnt” algorithm, based on climatic and edaphic variables, at the spatial resolution of 0.0833 ° in each grid cell, totalizing 13685 cells. The SDM was performed only for species with more than 15 occurrence points, which resulted in 2239 modeled species [41]. In the present study, we overlaid all the species ranges so each grid cell could represent an operational plant community. We also used the environmental suitability values from the previous study as a proxy for species abundance of each grid cell [42], assuming that both are correlated as previously shown for vertebrates, invertebrates and plants [42], throughout the Atlantic Forest.

### Traits

For the trait gathering, we focused on LHS (Leaf, Height and Seed) scheme traits, which synthesizes the leading dimensions of plant ecological strategies [43,44], and the wood economic spectrum. Here, these ecological strategy dimensions are represented by specific leaf area (SLA), maximum plant vegetative height at maturity (Hmax), seed dry mass (SDM), and stem specific density (SSD). Functional traits indicate trade-offs and life history stages [45–48], in a way that species ecological strategies vary from slow to fast growth [46]. Specific leaf area tends to be positively correlated with maximum photosynthetic rate per dry mass, growth rate and primary productivity, but inversely correlated with leaf longevity [45,46,49]. Plant height is a good predictor of plant light and water demands and hence its habitat [43,46,50]. Seed mass is inversely correlated with seed output per square meter, and positively correlated with seedling survival and size due to larger nutritious reserve [46]. Finally, denser stem is important in resisting drought and mechanical stress but it can reduce hydraulic conductivity and growth rates [46,51–53].

We searched for SLA, HMAX, SDM and SSD data first in the Botanical Information and Ecology Network (BIEN) database [54], the Plant-Trait database (TRY) [55], and Floristic and Forest Inventory of Santa Catarina database [56] (more details in Appendix S1 and S2 in Supplementary Information). HMAX was also compiled using the search tool of Reflora – Virtual Herbarium website [57]. Then, we searched in the literature (Appendix S3 in Supplementary Information) for information of the species that did not present trait information in the above-mentioned databases, and for information about species economic use of wood.

We considered as trait gap, at the species level, the presence or absence of each trait information for each species. To understand where gaps are geographically, we used a trait gap matrix with missing trait information per species and calculated the species proportion with trait gap per operational plant community.

To estimate the community trait dominance and dispersion we used the community weighted-mean [21] and the Rao’s quadratic entropy (*Q*) [58]. Each of these analyses requires two matrices. The first matrix consists of species per trait, while the second of grid cells per species abundance. Rao’s quadratic entropy is a measure of functional dissimilarities between species and higher values mean that a given community presents higher trait variation [58]. To understand where gaps are geographically, we used a trait gap matrix with missing trait information per species and used the calculated species trait gap proportion per operational plant community. We computed CWM using the function “functcomp” in the FD [59] and functional dispersion using the function “rao.diversity” in the package SYNCSA [60,61], in the R statistical software [62].

### Geographical data

We obtained the geographical coordinates of each urban area in Brazil from the Brazilian Institute of Geography and Statistics (IBGE– https://ibge.gov.br/). We obtained protected area polygons from the Chico Mendes Institute for Biodiversity Conservation (ICMBio – https://icmbio.gov.br). The protected areas belong to two groups: for sustainable use and for strictly protected area (Brazilian System of Protected Areas – SNUC, Law #9.985/2000). We filtered the urban areas up to 50,000 habitants and protected areas within the Atlantic Forest. Then we calculated the distance of each grid cell to the nearest urban area and to the nearest protected area. We calculated mean species range size of each grid cell, and the mean economic use of each grid cell by accounting for the sum of species with economic use.

### Data Analyses

#### Species

To assess whether economic use (binary data) and species range size (continuous data) bias species trait gaps (binary data), we fitted a generalized linear model (GLM) with a binomial distribution of the response variable. We fitted models with the two predictors, with only one predictor at a time, and also with only the intercept. We selected the model that best explained the relationship between variables using the Akaike’s information criterion (AIC), i.e. those with the lowest AIC value regarding the ΔAIC_i_ ≤ 2 criterion [63].

#### Trait gaps bias at the geographical level

In order to test which drivers lead to geographical bias in trait gaps we fitted GLMs using the proportion of species missing trait information at each grid cell as response variables and the mean species range and use, the distance from urban areas and distance from protected areas as predictor variables. We used GLMs with beta distribution, which uses maximum likelihood for parameters estimation, because our data ranged between of 0 and 1. Then, we selected the model with the lowest AIC value regarding the ΔAIC_i_ ≤ 2, and accounted for spatial autocorrelation [64] and Type I error [65,66] due to species cooccurences (more details in Appendix S5 and FigureS13 and 14 in Supplementary Information). We performed GLM using the function betareg in the R package [67,68].

To understand where the trait gap causes higher uncertainty in trait dominance and trait dispersion maps, we build bivariate maps considering the trait gap at the geographical level proportion (hereafter uncertainty degree) and the functiona structure measures estimated with the available information by grid cell. The supplementary material provides further information on analyses: bivariate function R script for building a bivariate figure (is provided in Appendix S4), single maps of CWM, QRAO, trait gaps and species richness (in FigureS1-12), and Atlantic Forest vegetation cover (in FigureS14).

### Data Deposition

Most of the data used in this paper is available in the BIEN and TRY databases. The additional data is now available in the TRY Plant Trait Database 6, under the name UFPR Atlantic Forest Tree Traits (UFPR_AFTT) [55]. Species range data is available at Zwiener et al., 2017 [41].

## RESULTS

### Species

We found that trait gap (percentage of species with missing data for each studied trait) among Atlantic Forest tree species was 88% for SDM, 85% for SLA, 65% for SSD and less than 1% for HMAX (see Appendix 6 in Supplementary information for details). Only 0.42% of the species have no information for any of the four traits, more than half (58%) missed information for at least three traits, followed by 12.3% with two traits missing information, and 6.2% with one trait missing information. Wood economic use and range size as predictor variables better-explained gaps of SLA, SDM and SSD. Species with wide range size and wood economic use had better trait data coverage. None of the predictor variables explained maximum height trait gap, given that all the models with different predictos combinations, including the model with only the intercept, presented ΔAIC_i_ ≤ 2 (See Appendix 7 in Supplementary information for AIC and GLM output tables and figure).

### Geographical

Mean wood economic use, mean range size, distance from protected areas and distance from urban areas explained trait gaps of SLA (pseudo R^2^=0.83), HMAX (pseudo R^2^=0.2215), SDM (pseudo R^2^=0.7987) and SSD (pseudo R^2^=0.8093 – Figure 1b). See AIC and GLM outputs tables in Appendix 8.1, Supplementary Information. That is to say, we know more about the species traits in places closest to protected areas, distant from urban areas, and with higher proportion of wide-ranged and economic used species. Spatial autocorrelation did not interfere in models results (Figure S15 in Supplementary Information). Only HMAX gap models presented a Type I error (*p*=0.49), possibly due to low gap spatial variation (Figure S16 in Supplementary information). Hence, we did not consider outputs from GLM regarding HMAX gap as significant.

**Figure 1:**
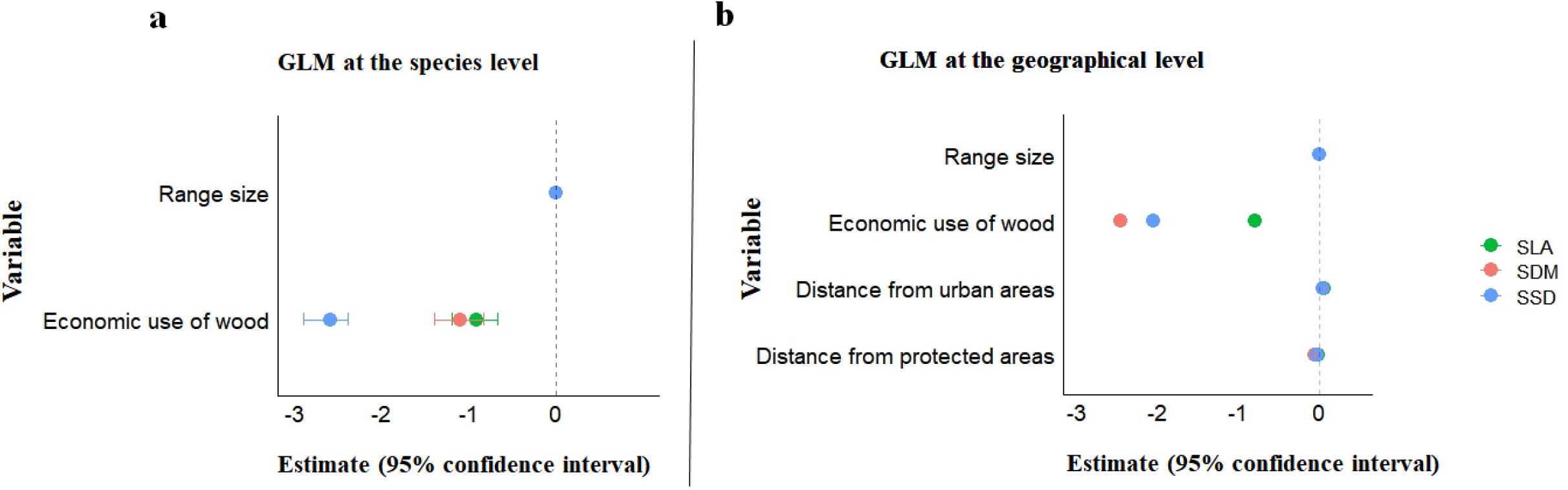
Colored dots are observed slope estimates for traits and horizontal bars are 95% confidence intervals. (a) Trait data gaps at the species level are negatively biased towards species mean range size and economic use of wood for specific leaf area (green), seed dry mass (pink) and stem specific density (blue); (b) Trait data gaps at the geographical level are negatively biased towards species mean range size, economic use of wood and distance from protected areas, and positively biased towards distance from urban areas for specific leaf area (green), seed dry mass (pink) and stem specific density (blue).

In general, trait gap was higher in the eastern Atlantic Forest and lower in the southwestern and central northeast, except for maximum height. The gap of SLA ranged from 57% to 80%, of SDM from 63% to 85% and of SSD from 36% to 62%. The gap of HMAX ranged from nearly zero to 1.2% in the eastern Atlantic Forest (Figure 2). See Appendix 8.2, in Supplementary Information for detailed description of functional structure and dispersion.

**Figure 2.**
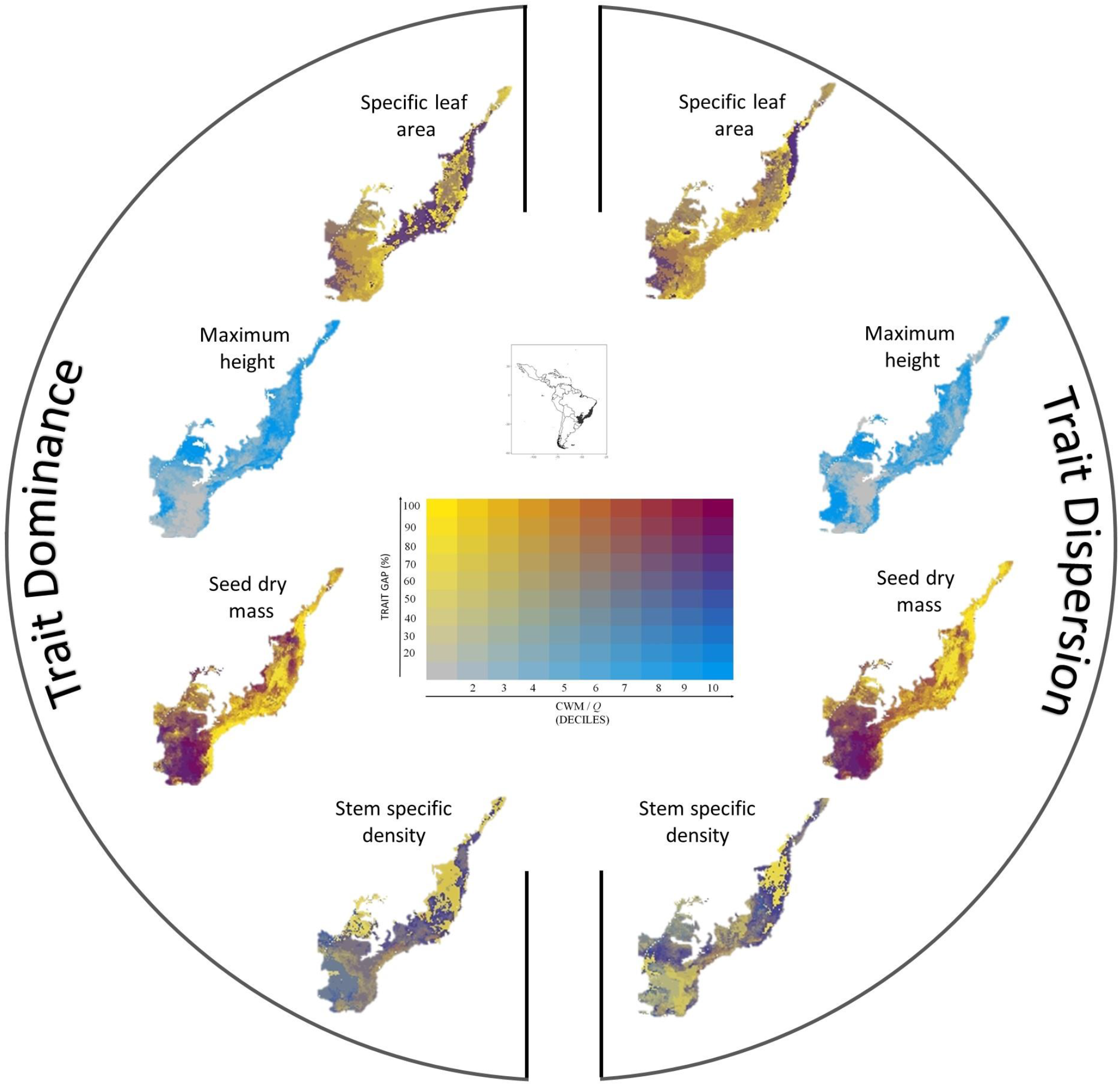
Trait dominance (CWM) and trait dispersion (*Q*) for specific leaf area, maximum height, seed dry mass, and stem specific density in the Atlantic Forest.

## DISCUSSION

Our results showed that the trait gap is higher for Atlantic Forest tree species that have small ranges and no economical value. Also, we tend to know less about Atlantic Forest tree species regarding their traits in places distant from main cities, closest to protected areas, and within communities composed by small-ranged and with less species with economic use. These biases in both species and geographical level might compromise conservation assessments on biodiversity distribution, decline and its relation to ecosystem services [1,10]. Therefore, it is paramount that efforts to fill these gaps increase, in order to better inform theoretical and applied science.

### Trait gaps at the species level

In this study, we showed large trait gaps for key functional traits among Atlantic Forest trees, and that these gaps are biased towards small-ranged species and species with no economic use. Bias related to range size, i.e. the greater the range the more we know, could lead to incomplete knowledge of important roles in ecosystem services, especially if rare species fulfill ecosystem key-roles [69,70], not provided by common widely distributed species traits [71]. Also, we found more trait information for tree species with no economic use in Atlantic Forest, which migh lead to dichotomic paths, since economic use can strengthen arguments for conservation [72], but it can also be a reason for biodiversity decline through overexploitation [73].

We actually found that economic use might indirectly strengthen conservation, as species with economic use of their wood are better studied, even those with small range sizes compared to those with no economic use. Moreover, it is necessary to drive attention to trait gaps of species without economic use in order to fill these gaps. In the future, climate change could alter the type and strength of biodiversity-ecosystem relationship, changing the dominance of species with different ecological requirements, which can include species with missing information [74,75]. Then, species with missing information may not be a research focus now, but can be in the future by ensuring ecosystem services.

The small trait gap for HMAX is a direct consequence of available information from exsiccates in the REFLORA project, and from published books used in this study. Such availability of data is a direct result of HMAX being easily measurable and widely reported [76]. Unfortunately, traits such as SLA, SDM and SSD, that are harder to measure, are the ones usually missing [1] and can not be reported in exsiccates tags. HMAX is the trait synthesizing the major dimension of plant trait variation when both herb and woody life forms of varying habitats are considered all together [77]. In this study, however, we are considering only woody plants of the Atlantic tropical forest. In this case, SLA, SDM and SSD appear to be most relevant traits to functionally distinguish among species [78,79].

### Functional trait gaps at the geographical level

Our study showed that communities closer to urban areas, composed by wide range-sized species and species with economic use of wood presented lower trait gaps. Although the number of species records is usually higher near protected areas [6,33], we did not find evidence that communities closer to these areas are better-known regarding their traits, so that increasing sampling effort near protected areas may not be reflected more in trait knowledge. The substantial amounts of trait gap near protected areas might occur because protected areas historically sheltered rare species with specific habitat needs [80], and limited access to these areas coupled to low detectability could impair rare species sampling [81]. Most of protected areas coverage is in the eastern Atlantic Forest, especially in the Serra do Mar area [38], which is a region with high species richness [82]. Species richness might play an important role in trait knowledge in Atlantic Forest, which in turn might lead to regions with high species richeness that have more unknown species than others, even more if inside protected areas.

Trait-based approaches directed to conservation might be impaired if focal communities have large trait gaps. Eastern Atlantic Forest has high tree and tree-like species richness [82] and is in debt of 70 to 90% of area required to restore native vegetation under the new Brazilian Forest Code [40]. In the southwestern and northwestern Atlantic Forest, which have also high tree and tree-like species richness [82], restoration status is even more critical with over 90% of their areas in debt for native vegetation restoration [40]. The good news for these regions, are their smaller trait gaps. Better mean trait knowledge may be useful in theory-drive restoration [16,83], enabling restoration practices concerning biodiversity-ecosystem function in the southwestern and northwestern Atlantic Forest.

If bias of trait data is restrained towards species that display specific functional traits values, such as in areas with low functional dispersion and high trait gaps, there could be underestimation of functional diversity measures [3]. For instance, if only data available in databases is used to estimate carbon storage, and only species with high wood density have been sampled, we could misinterpret ecosystem functions, by over estimating carbon stock. Thus, trait gaps might also limit the applicability of trait-based approaches in restoration practices under the biodiversity-ecosystem approaches in areas with higher trait gaps, since previous functional trait data will be required for implementing that approach.

### Future Directions

We are approaching the deadline proposed by the Aichi Biodiversity Targets in the Convention on Biological Diversity [84]. The Target 19 of the Strategic Plan for Biodiversity 2011-2020 includes, among others, knowledge improvement in biodiversity-ecosystem functioning. Even with the comprehensive coverage information presented by databases [55,85], here we show the necessity to improve data coverage of Atlantic Forest tree traits, a biodiversity *hotspot* providing ecosystem services for 148 million humans [86]. Moreover, data gaps of traits that are related to climatic change should be considered, as climatic change might impact biodiversity-ecossystem relationships [74,75]. For instance, knowledge about traits such roots and plant-pollinator properties that are related to plant resistence to drought and competitive abilities [87], and food security [88], respectively.

Whilst approaches on ecosystem-biodiversity function and services are under the spotlight, trait knowledge is indispensable to properly consider them in conservation and restoration of biodiversity [89]. If trait gaps are reduced in the future, trait-based information, such as functional diversity and functional structure measures, could be a shortcut to predict ecosystem-biodiversity relationships in conservation planning [90,91].

## ACKNOWLEDGMENTS

This study was financed in part by the Coordenação de Aperfeiçoamento de Pessoal de Nível Superior - Brasil (CAPES) - Finance Code 001. FTB currently holds a post-doctoral fellowship grant from Programa Nacional de Pós-Doutorado from Coordenação de Aperfeiçoamento de Pessoal de Nível Superior (PNPD/CAPES, Grant #88882.306081/2018-1). The authors thank Gabriela S. Müller, Luan H. Burda, Ursula Morais and Josué Araldi for the help in data collection; Isabela Galarda Varassin and Bruno Vilela de Moraes e Silva for their valuable suggestions to the manuscript. This paper is developed in the context of the National Institutes for Science and Technology (INCT) in Ecology, Evolution and Biodiversity Conservation, supported by MCTIC/CNpq (proc. 465610/2014-5) and FAPEG (proc. 201810267000023).

