## Supplementary Information for "Minding The Gap: Range Size And Economic Use Drive Functional Trait Data Shortfall In The Atlantic Forest"

**SUPPLEMENTARY MATERIAL**

***Supplementary Figures***

Trait data gap at the geographical level for specific leaf area, maximum height, seed dry mass and wood density.

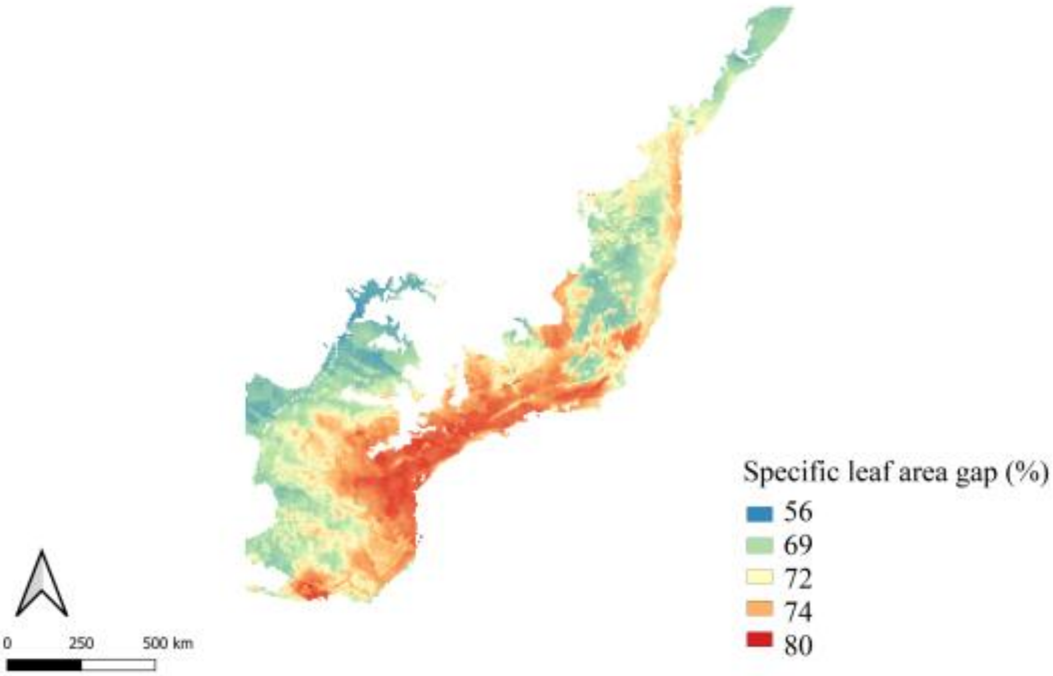


FigureS 1: Specific leaf area gap at the geographical level

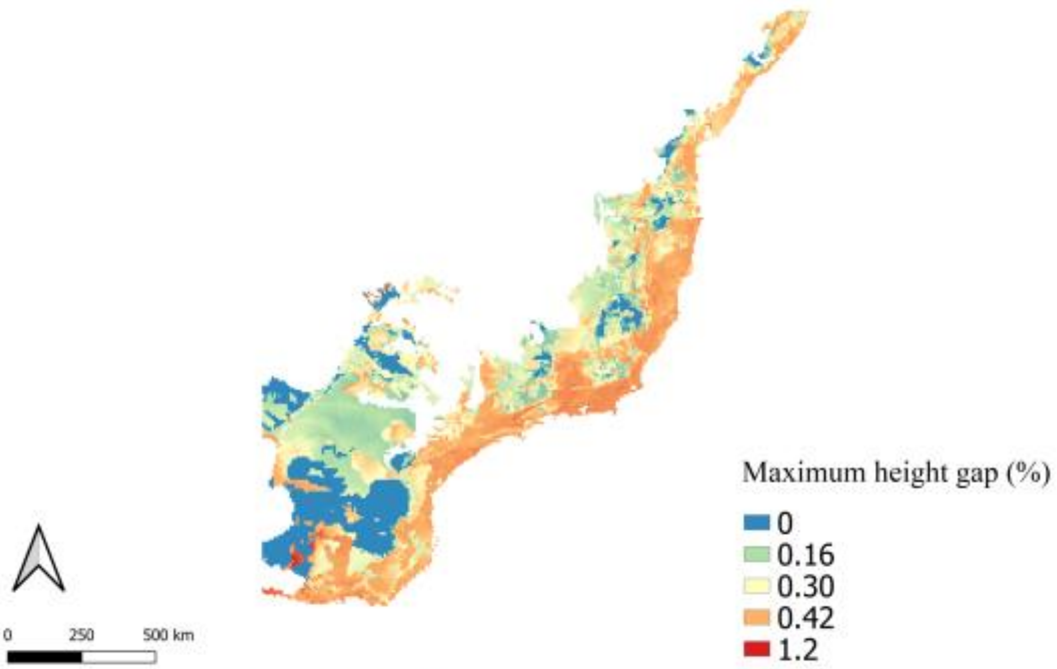


FigureS 2: Maximum height gap at the geographical level.

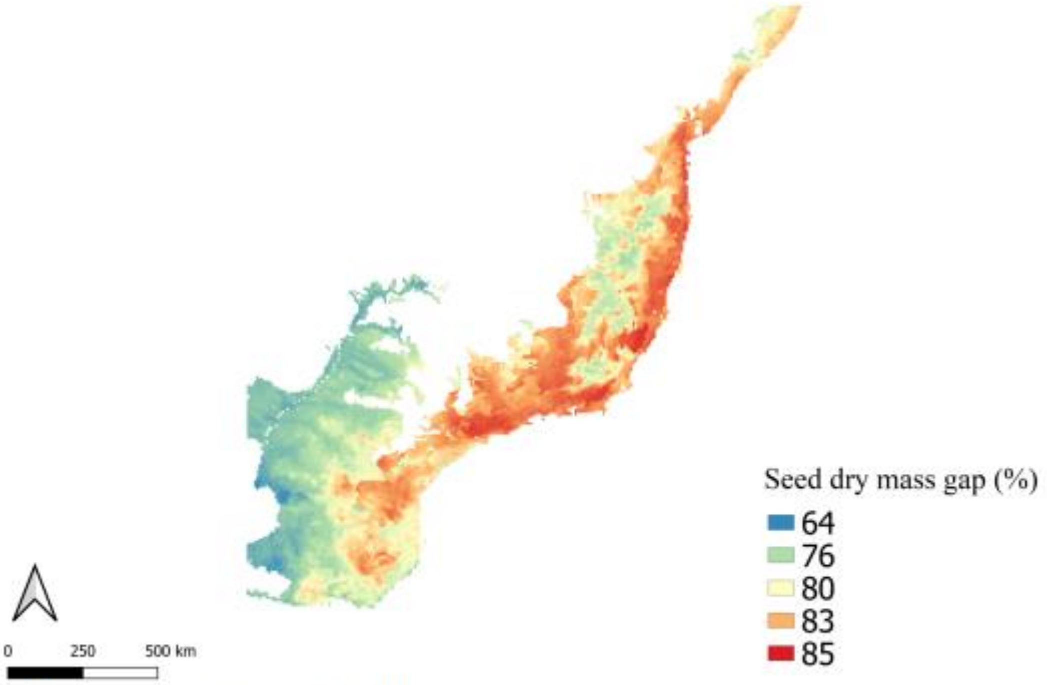


FigureS 3: Seed dry mass gap at the geographical level.

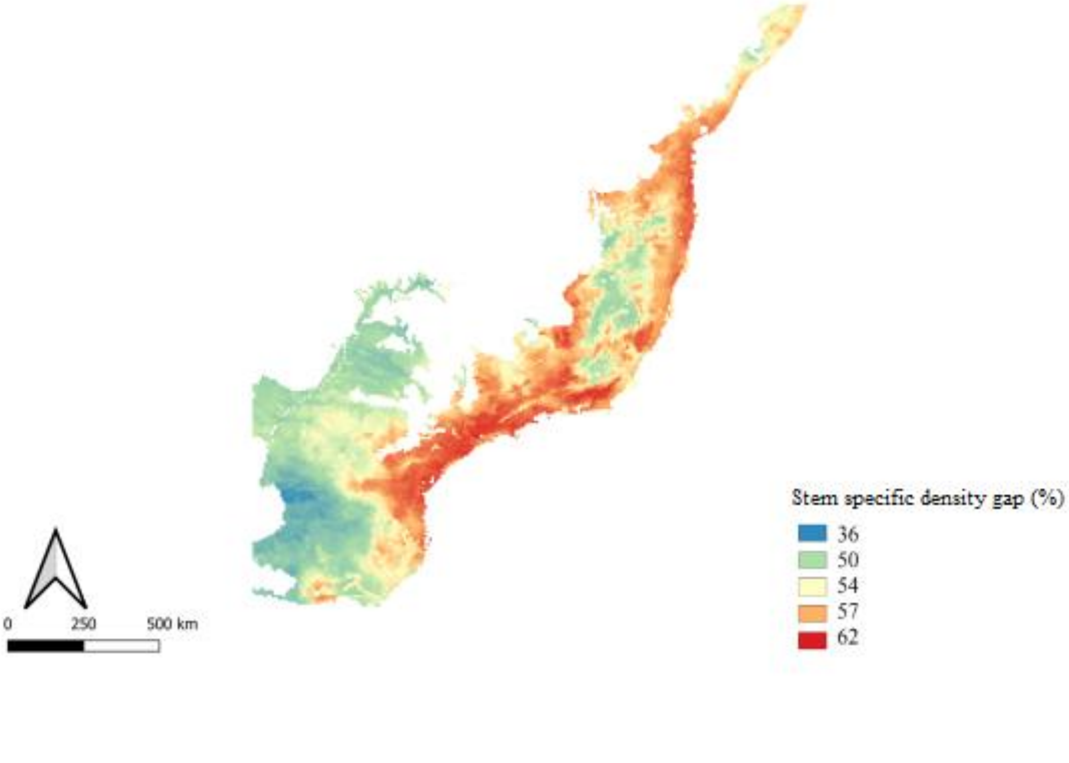

FigureS 4: Stem specific density gap at the geographical level.

Trait dominance (CWM) for specific leaf area, maximum height, seed dry mass and specific stem density.

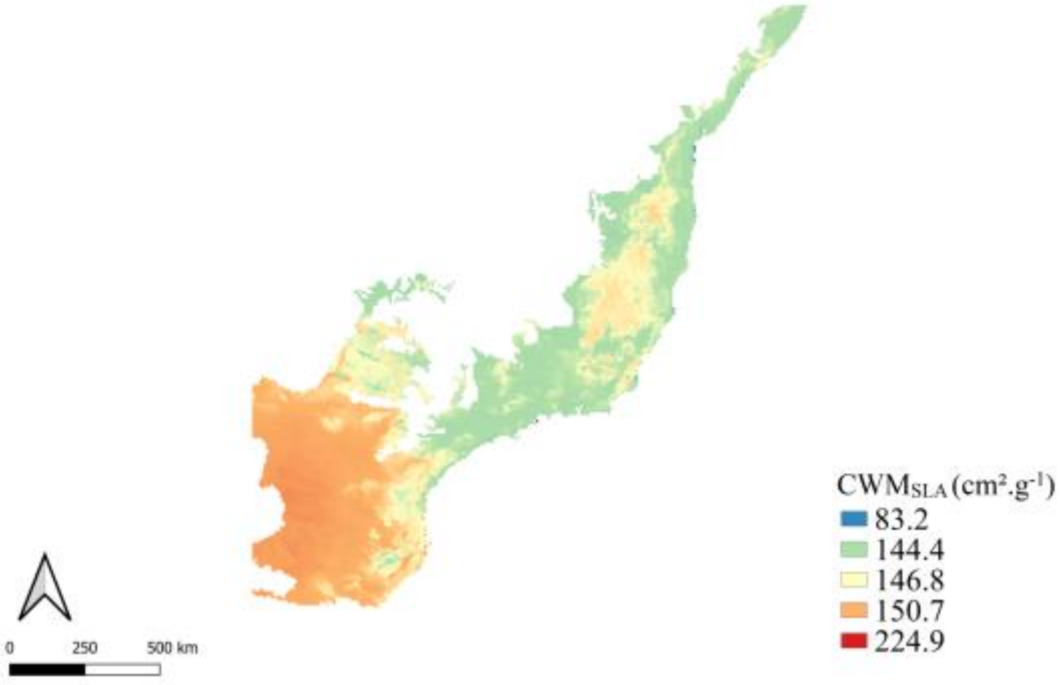


FigureS 5: Trait dispersion of specific leaf area (SLA)

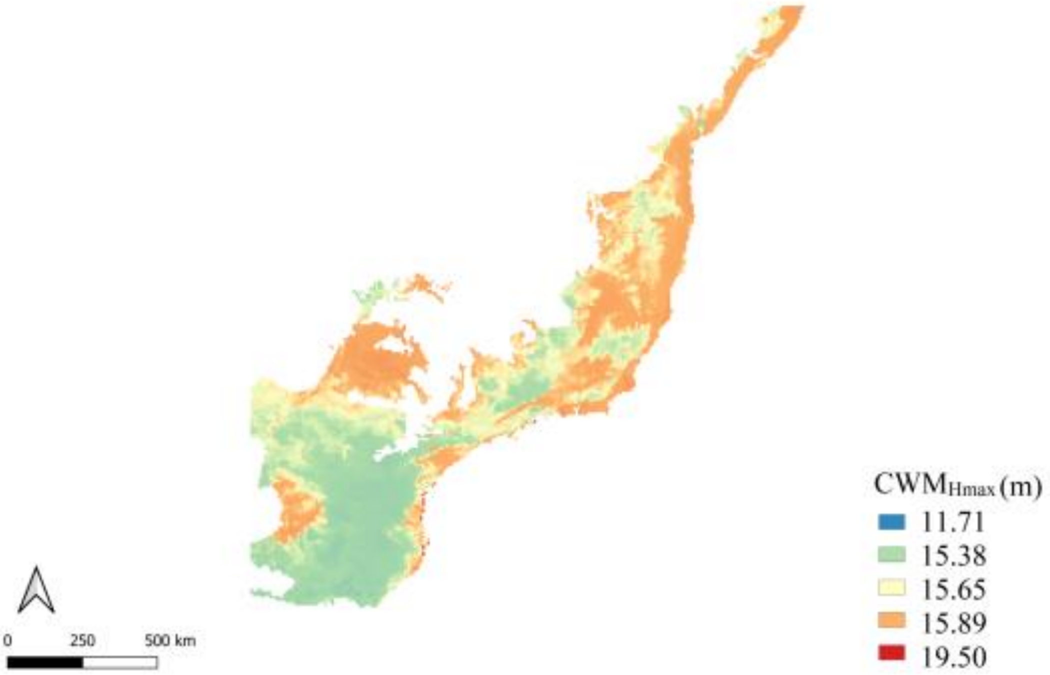


FigureS 6: Trait dominance of maximum height (Hmax)

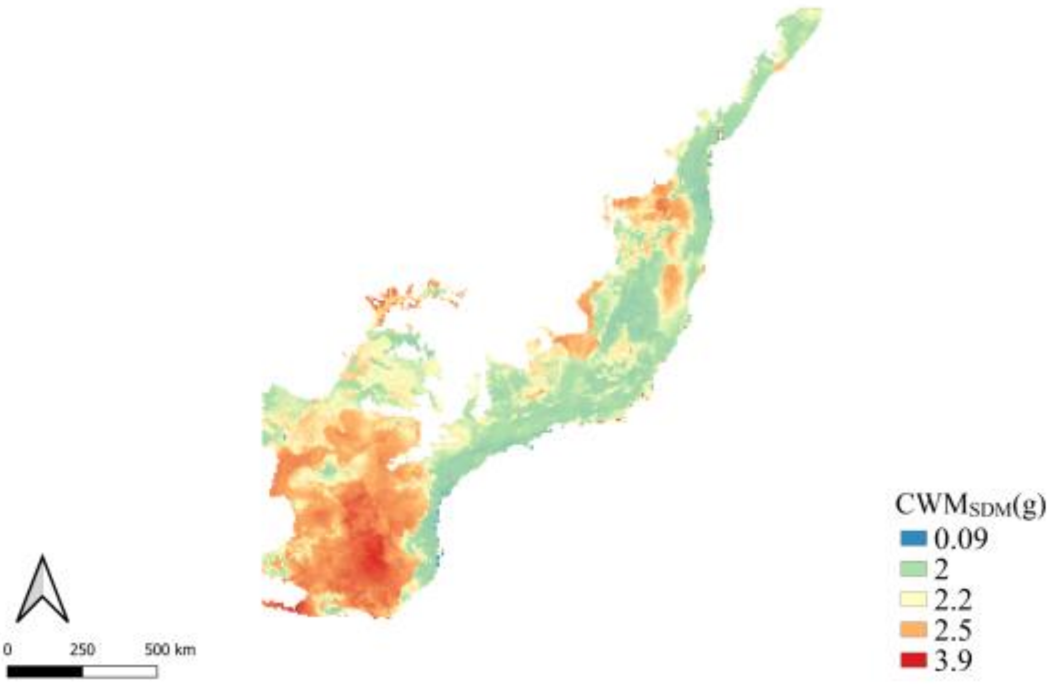


FigureS 7: Trait dominance of seed dry mass (SDM)

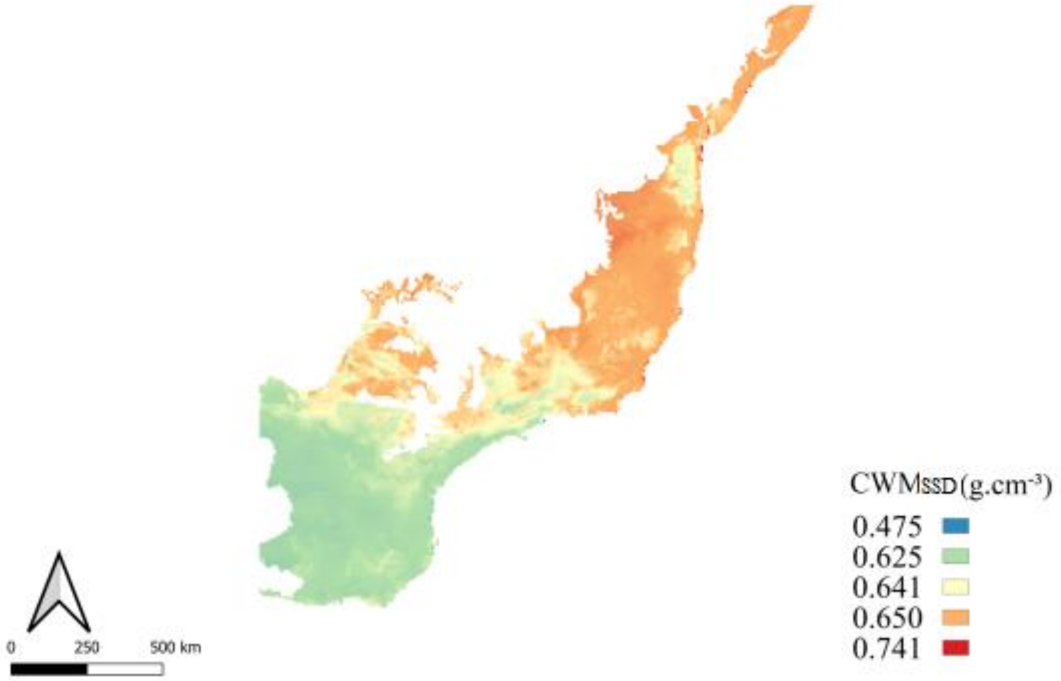

Figure S 8: Trait dominance of stem specific density (SSD)

Trait dispersion ($Q$) for specific leaf area, maximum height, seed dry mass and stem specific density.

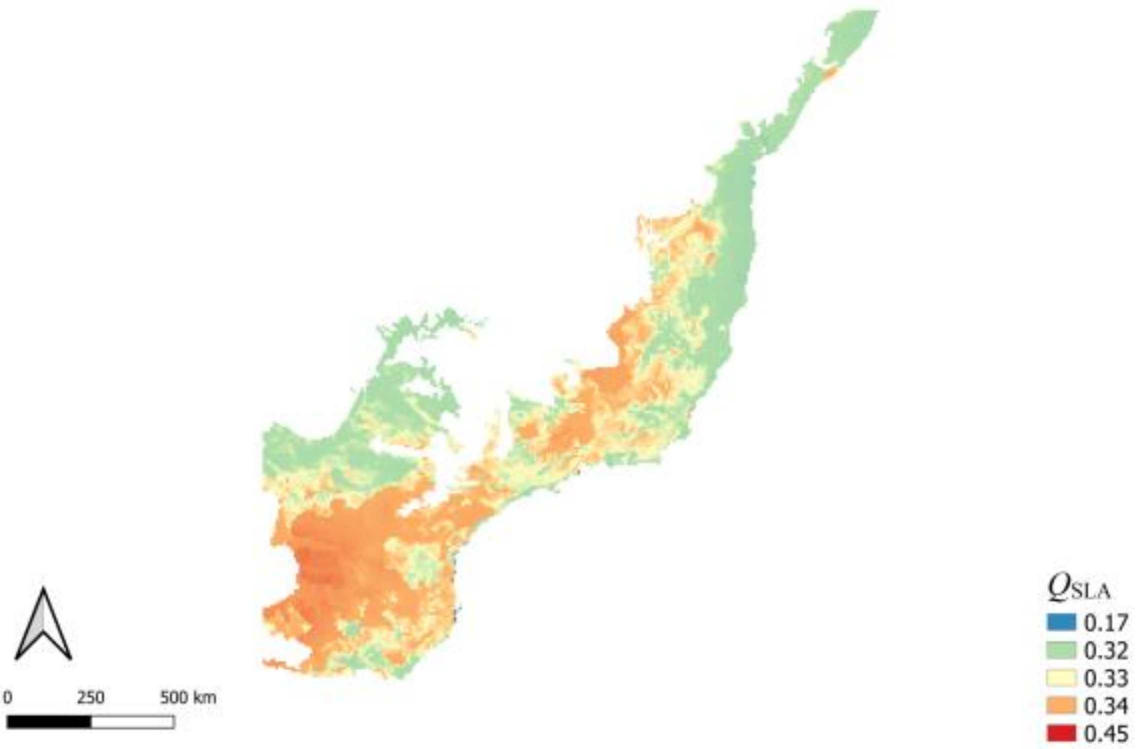


FigureS 9: Trait dispersion of specific leaf area (SLA)

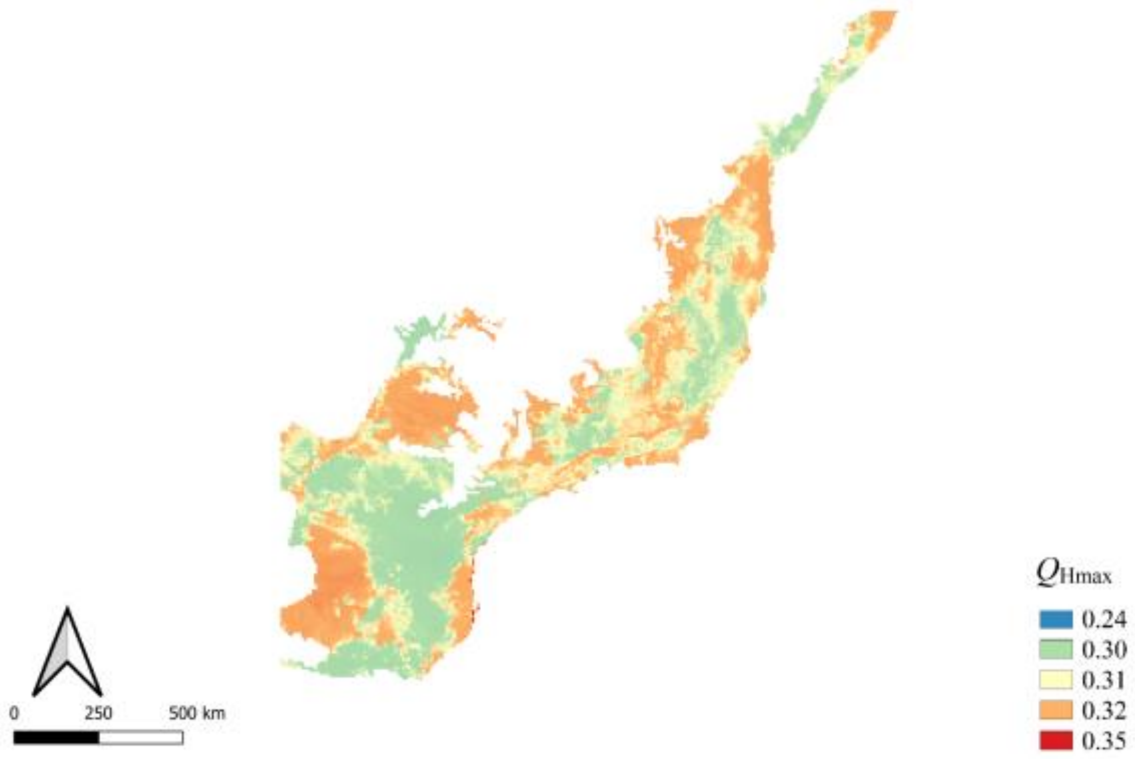


FigureS 10: Trait dispersion of maximum height (Hmax)

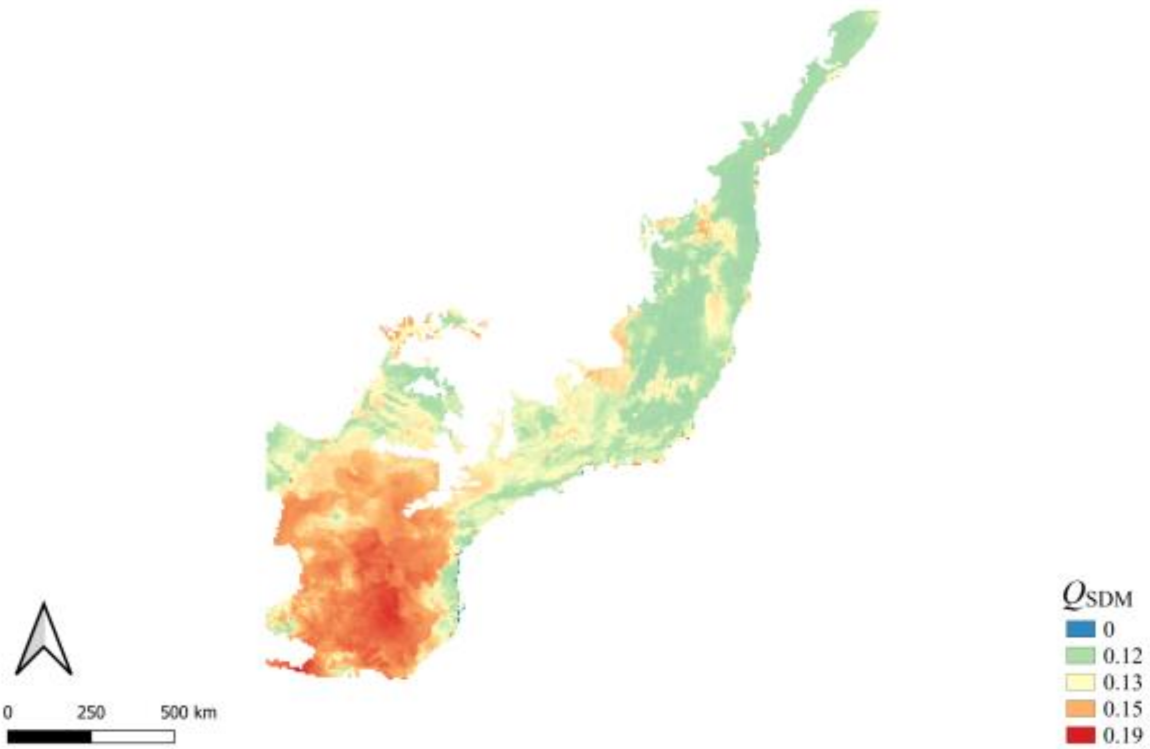


FigureS 11: Trait dispersion of seed dry mass (SDM)

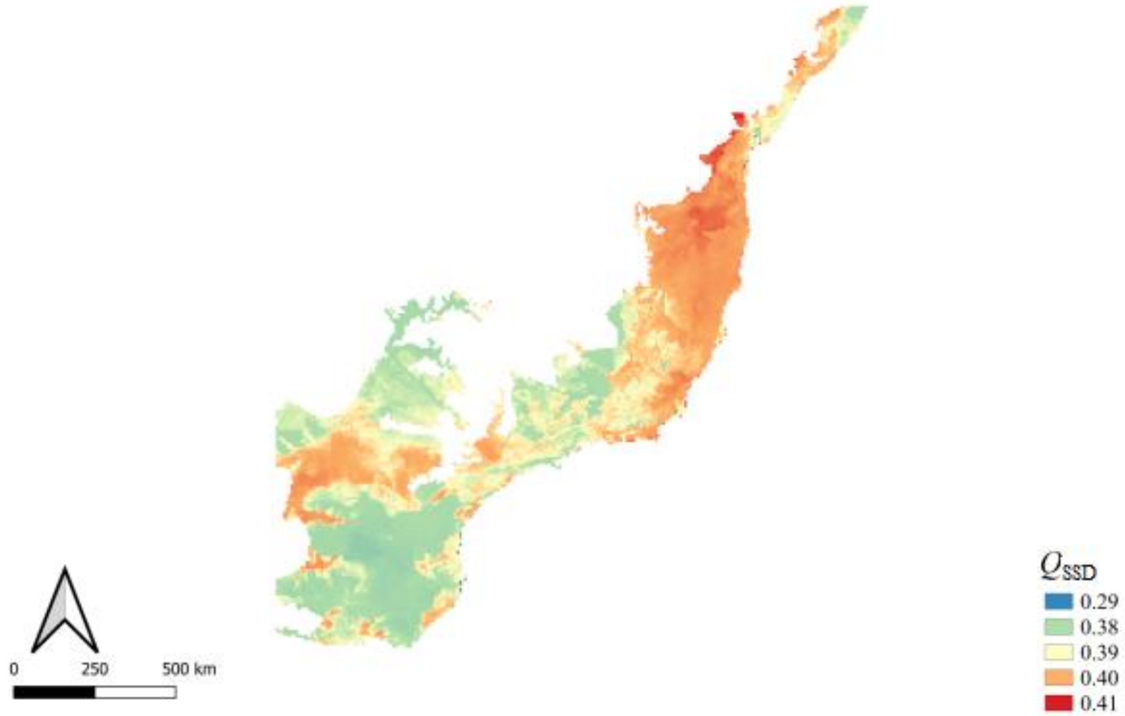

FigureS 12: Trait dispersion of stem specific density (SSD)

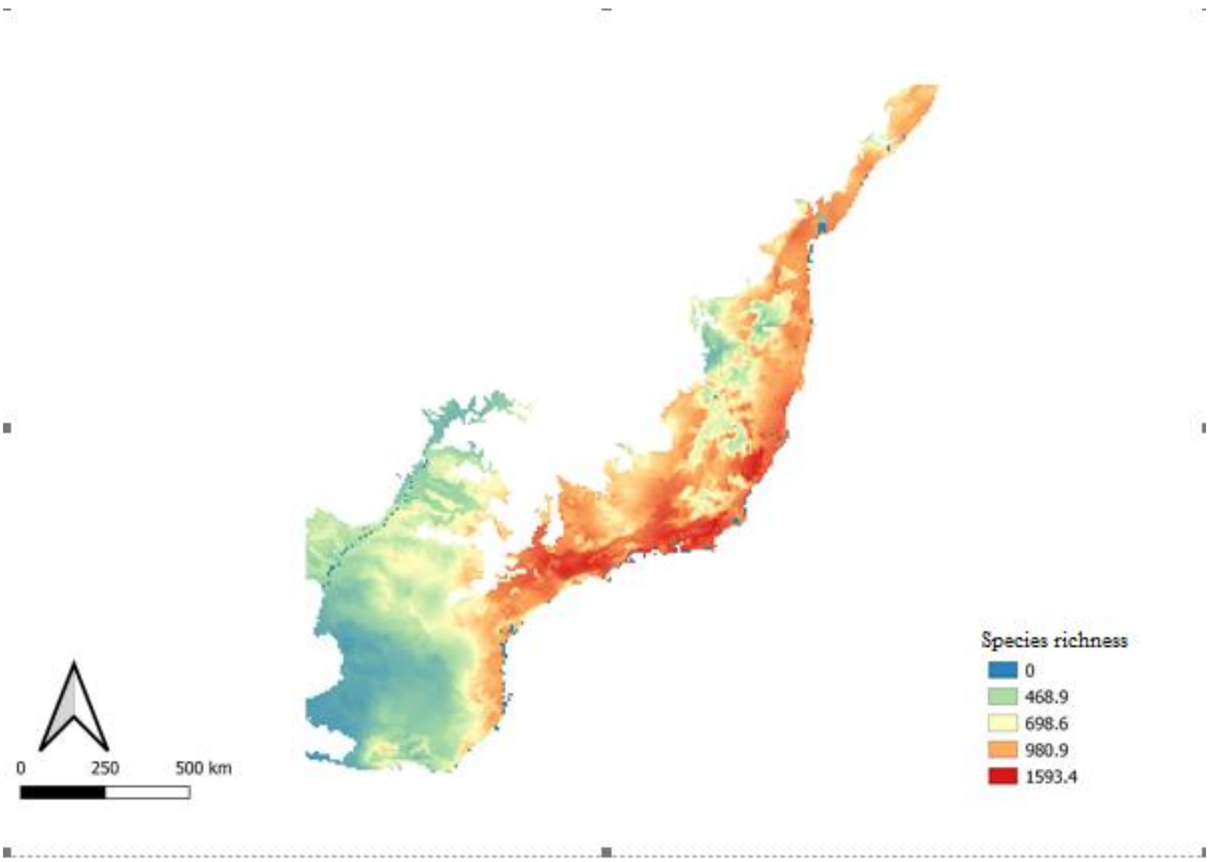


FigureS 13 Atlantic Forest richness of tree species

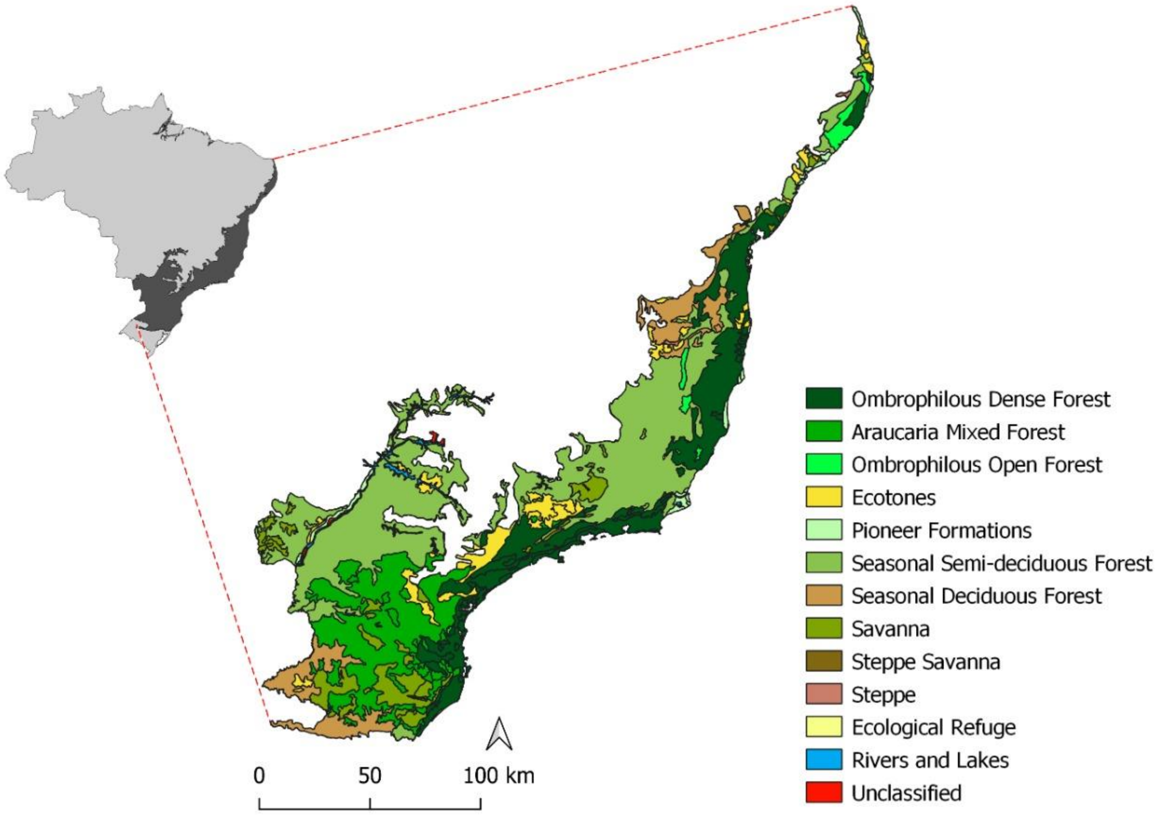


Figure S14 Vegetation cover map of Atlantic Forest from the Brazilian Institute of Geography and Statistics (IBGE– https://ibge.gov.br/).

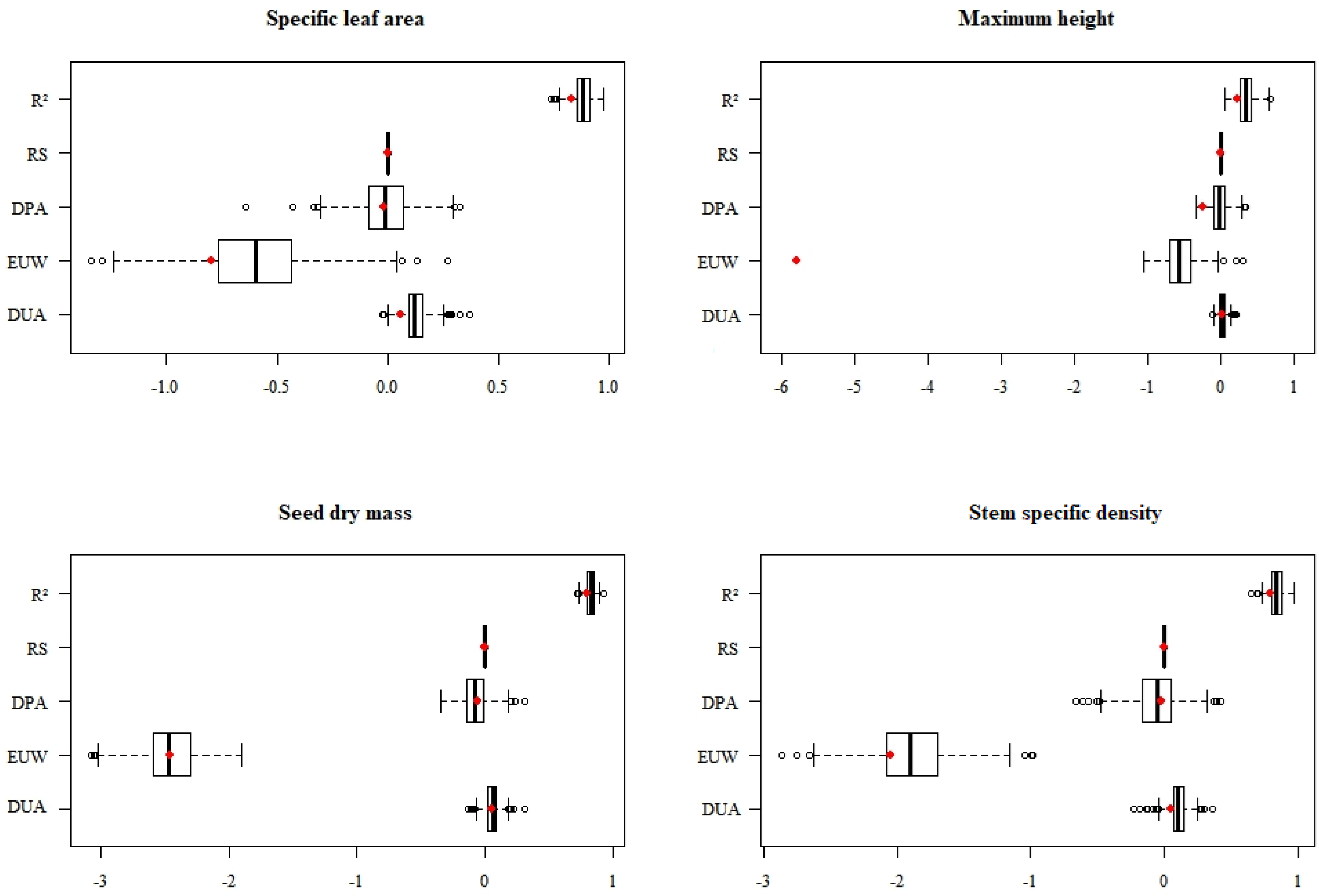


Figure S15 Estimates of 500 fitted GLM from thinned spatial points when Moran's I correlation was low (*I*=0.1). Red dots represent the observed estimates from GLM models with all spatial points. Vertical points are 90% of conficence interval. $R^2$=pseudo-R squared, RS=Range size, DPA=Distance from protected areas, EUW=Economic use of wood, DUA=Distance from urban areas.

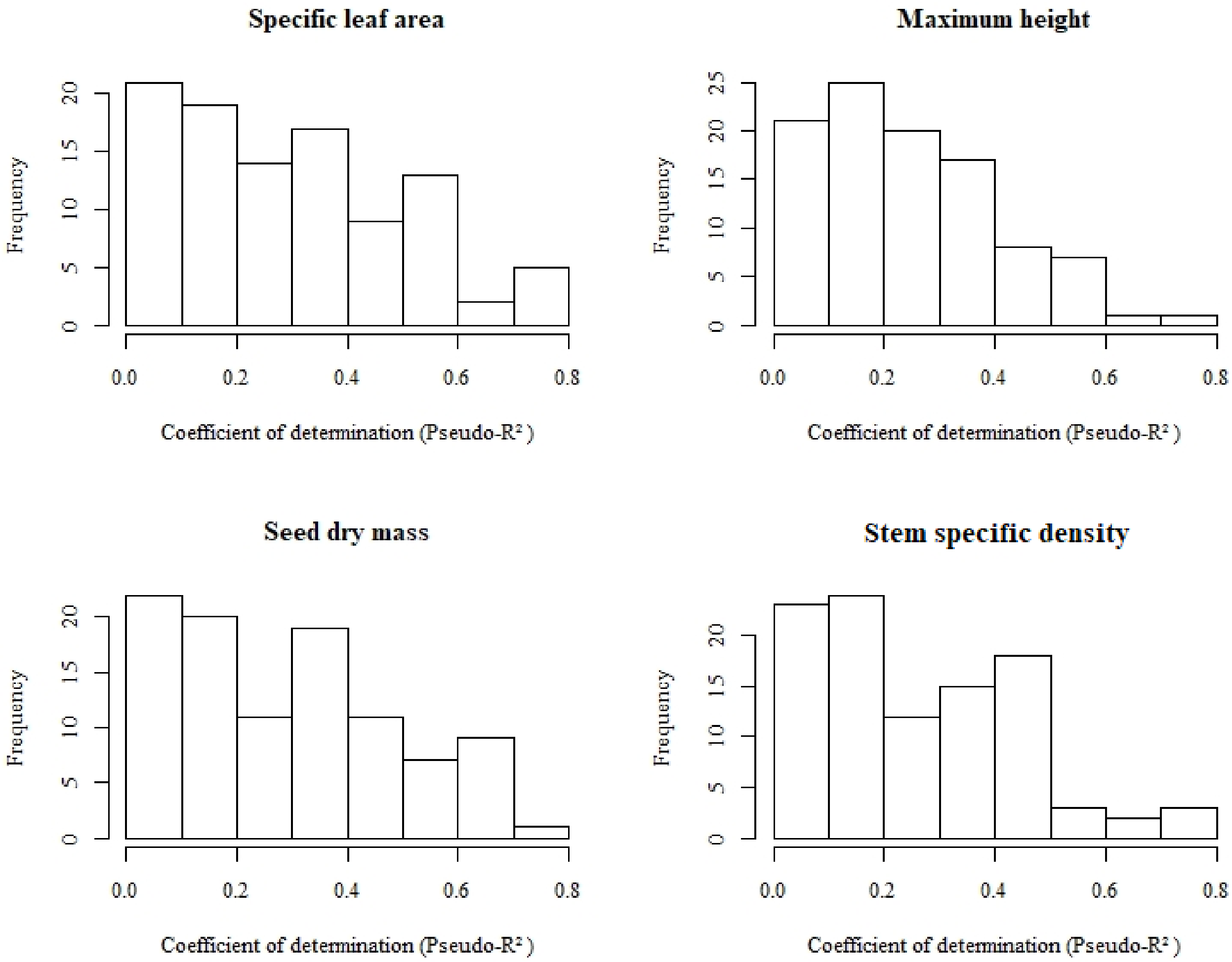


Figure S16: Coefficient of determination (Pseudo-$R^2$) frequency from 100 permutations in trait gap matrix used to model geographical trait gap response to distance from urban areas, distance from protected areas, mean economical use of wood and mean range size.

### *Appendix S1*

We assessed BIEN database through the Bien R package using the function “BIEN.trait.traitbyspecies”. When we used the search tool of Reflora *online herbarium* we obtained organized spreadsheets with all the exsicatas per species in rows, which contained characteristics of the collected individual, as height. We compiled the maximum height of an adult individual registers per species by report.

### *Appendix S2*

Databases used from BIEN and TRY.

*Table S21* Databases for specif leaf area, maximum height, seed dry mass and stem specific density in BIEN database

| Authorship | Contact | Dataset |
|---|---|---|
| Lopez-Gonzalez G | | http://datadryad.org/resource/doi:10.5061/dryad.234 |
| Zanne, A.E., et. al | | http://datadryad.org/repo/handle/10255/dryad.235 |
| T.A. Easdale | | http://dx.doi.org/10.1016/j.ppees.2009.03.001 |
| Ploton P | | http://datadryad.org/resource/doi:10.5061/dryad.f2b52 |
| Goodman RC | | http://datadryad.org/resource/doi:10.5061/dryad.p281g |
| Paine CET | | http://datadryad.org/resource/doi:10.5061/dryad.h9083 |
| Bhaskar R | | http://datadryad.org/resource/doi:10.5061/dryad.6p9v5 |
| Chacón E | | NA |
| Mascaro J | | http://datadryad.org/resource/doi:10.5061/dryad.rs7b0.2 |
| Szefer P | | http://datadryad.org/resource/doi:10.5061/dryad.4b95c.2 |
| Nathan Kraft | | doi:10.1890/09-1672.1 |
| Rasmann S | | http://datadryad.org/resource/doi:10.5061/dryad.8557 |
| L. Poorter | | doi: 10.1890/0012-9658(2006)87[1733:LTAGPO]2.0.CO;2 |
| Maire V | | http://datadryad.org/resource/doi:10.5061/dryad.j42m7.2 |
| L. Poorter | | doi:10.1007/s00442-008-1131-x |
| Benjamin Blonder | | www.amjbot.org/content/suppl/2012/11/07/ajb.1200062.DC1/Blonder_AppS1A_observational_data.csv |
| Price CA | | http://datadryad.org/resource/doi:10.5061/dryad.r3n45 |
| Milla R | | http://datadryad.org/resource/doi:10.5061/dryad.dg85v |
| Michael Kleyer | | http://www.leda-traitbase.org/LEDAportal/ |
| Charles Price | | http://onlinelibrary.wiley.com/doi/10.1111/1365-2435.12298/suppinfo |
| Grootemaat S | | http://datadryad.org/resource/doi:10.5061/dryad.m41f1 |
| Boyero L | | http://datadryad.org/resource/doi:10.5061/dryad.jg8r0 |
| Bufford JL | | http://datadryad.org/resource/doi:10.5061/dryad.b1v2c |
| Perezt F | | http://datadryad.org/resource/doi:10.5061/dryad.d61jk |
| Dana | | NA |

| | | |
|---|---|---|
| Royer | | |
| Liu, K., Eastwood, R.J., Flynn, S., Turner, R.M., and Stuppy, W.H. | | http://www.kew.org/data/sid |
| Letcher SG | | http://datadryad.org/resource/doi:10.5061/dryad.d87v7 |
| Kraft TS | | http://datadryad.org/resource/doi:10.5061/dryad.69ph0 |
| Fricke EC | | http://datadryad.org/resource/doi:10.5061/dryad.90f03 |
| Tim Killeen | NA | NA |
| Susan Letcher | | NA |
| Greg Reams | NA | NA |
| NA | NA | http://www.americanforests.org/resources/bigtrees/register.php |
| L. Poorter | | doi: 10.1093/aob/mcn103 |
| Saara J. DeWalt | | NA |
| NA | NA | NA |
| Kleinschroth F | | http://datadryad.org/resource/doi:10.5061/dryad.51p4f |
| Thomas SC | | http://datadryad.org/resource/doi:10.5061/dryad.bs332 |
| Osuri AM | | http://datadryad.org/resource/doi:10.5061/dryad.7s7r1 |

*Table S2 2*Databases for specif leaf area, maximum height, seed dry mass and stem specific density in TRY database

| Autorship | Dataset ID | Dataset |
|---|---|---|
| Higgins, Steve et. al | 48 | Dispersal Traits Database |
| Niinemets, Ülo et. al | 87 | Global Leaf Robustness and Physiology Database |
| Lloyd, Jon et. al | 34 | The RAINFOR Plant Trait Database |
| Wirth, Christian et. al | 68 | The Functional Ecology of Trees (FET) Database - Jena |
| Pillar, Valerio et. al | 75 | ECOQUA South American Plant Traits Database |
| Sosinski, Enio et. al | 77 | FAPESP Brazil Rainforest Database |
| Wright, S. Joseph et. al | 112 | Panama Plant Traits Database |
| Wright, Ian et. al | 20 | GLOPNET - Global Plant Trait Network Database |
| Wright, Ian et. al | 64 | Neotropic Plant Traits Database |
| Finegan, Bryan et. al | 74 | Costa Rica Rainforest Trees Database |
| Kattge, Jens et. al | 67 | Leaf Physiology Database |
| Domingues, Tomas et. al | 255 | LBA ECO Tapajos: Leaf Characteristics and Photosynthesis |

| | | |
|---|---|---|
| Jackson, Robert et. al | 240 | Nutrient Resorption Efficiency Database |
| Jansen, Steven et. al | 241 | Xylem Functional Traits (XFT) Database |
| Craven, Dylan et. al | 230 | Panama Tree Traits |
| Powers, Jennifer et. al | 263 | Costa Rican Tropical Dry Forest Trees |
| Gonzalez-Melo, Andres et. al | 267 | Functional Traits for Restoration Ecology in the Colombian Amazon |
| Schweingruber, Fritz et. al | 251 | The Xylem/Phloem Database |
| Baraloto, Christopher et. al | 269 | The Bridge Database |
| Lenti, Felipe et. al | 274 | Crown Architecture Database |
| Atkin, Owen et. al | 286 | Global Respiration Database |
| Higuchi, Pedro et. al | 305 | Araucaria Forest Database |
| Holl, Karen et. al | 306 | Plant traits from Costa Rica |
| Mazzochini, Guilherme et. al | 357 | Functional traits of woody species in the Brazilian semi-arid region |
| Maire, Vincent et. al | 342 | Photosynthesis Traits Worldwide |
| Dias, Arildo et. al | 368 | Wood traits of trees and lianas from the Brazilian Atlantic Forest |
| Souza, Alexandre et. al | 369 | Traits and ecological strategies of 66 subtropical tree species in the Brazilian Atlantic Forest |

***Appendix S3***

We used the fallowing literature for functional trait compilation


1. Carvalho PER. 2003 *Espécies Arbóreas Brasileiras - Vol I*. Embrapa.

2. Carvalho PER. 2006 *Espécies Arbóreas Brasileiras - Vol II*. Embrapa.

3. Carvalho PER. 2008 *Espécies Arbóreas Brasileiras - Vol III*. Embrapa.

4. Carvalho PER. 2010 *Espécies Arbóreas Brasileiras - Vol IV*. See Embrapa.

5. Carvalho PER. 2014 *Espécies Arbóreas Brasileiras - Vol V*. Embrapa.

6. Lorenzi H. 2016 *Árvores Brasileiras: Manual de Identificação e Cultivo de Plantas Arbóreas Nativas do Brasil - Vol III*. 2nd edn. Instituto Plantatum.

7. Lorenzi H. 2002 *Árvores Brasileiras: Manual de Identificação e Cultivo de Plantas Arbóreas Nativas do Brasil - Vol II*. 2nd edn. Instituto Plantatum.

8. Lorenzi H. 2002 *Árvores Brasileiras: Manual de Identificação e Cultivo de Plantas Arbóreas Nativas do Brasil - Vol I*. 4th edn. Instituto Plantatum.

9. Durigan G, Baitello JB, Corrêa GAD, Siqueiza MF. 2004 *Plantas do Cerrado Paulista: Imagens de uma Paisagem Ameaçada*. Páginas & Letras Editora e Gráfica.

10. Backes. P, Irgang. B. 2002 *Árvores do Sul: Guia de Identificação & Interesse Ecológico*. 1st edn. Ed. Clube da Árvore.

11. Backes. P, Irgang. B. 2004 *Mata Atlântica: As Árvores e a Paisagem*. 1st edn. Editora Paisagem do Sul.

12. Paula JE, Alves JL de H. 2010 *922 Madeiras Nativas do Brasil*. 1st edn. Cinco Continentes.


***Appendix S4***

```
#### Function adapted from Hijmans in https://stackoverflow.com/questions/54144269/bivariate-choropleth-map-in-r (Acessed september 2020) ###
############# Mapa bivariado para diversos atributos, fixando os valores do eixo Y para todos os atributos entre 0 e 100% #############

library(raster)

#### Functions ####
makeCM <- function(breaks=10, upperleft, upperright, lowerleft, lowerright) {
  m <- matrix(ncol=breaks, nrow=breaks)
  b <- breaks-1
  b <- (0:b)/b
  col1 <- rgb(colorRamp(c(upperleft, lowerleft))(b), max=255)
  col2 <- rgb(colorRamp(c(upperright, lowerright))(b), max=255)
  cm <- apply(cbind(col1, col2), 1, function(i) rgb(colorRamp(i)(b), max=255))
  cm[, ncol(cm):1 ]
} #Generating colors by axis

plotCM <- function(cm, xlab="", ylab="", main="") {
  n <- cm
  n <- matrix(1:length(cm), nrow=nrow(cm), byrow=TRUE)
  r <- raster(n)
  cm <- cm[, ncol(cm):1 ]
```

```
  image(r, col=cm, axes=FALSE, xlab=xlab, ylab=ylab, main=main)
} # figura com a legenda com as cores

#rasterCM <- function(x, y, n) {
  #q1 <- quantile(x, seq(0,1,1/(n)))
  #q2 <- quantile(y, seq(0,1,1/(n)))
  #r1 <- cut(x, q1, include.lowest=TRUE)
  #r2 <- cut(y, q2, include.lowest=TRUE)
  #overlay(r1, r2, fun=function(i, j) {
  #  (j-1) * n + i
  #})} ### Original function

rasterCMmod <- function(x, y, n) {
  q1 <- quantile(x, seq(0,1,1/(n)))
  q2 <- quantile(seq(0,100,1), probs=seq(0,1,1/n))
  r1 <- cut(x, q1, include.lowest=TRUE)
  r2 <- cut(y, q2, include.lowest=TRUE)
  overlay(r1, r2, fun=function(i, j) {
    (j-1) * n + i
  })
}   #Fixing y axis to vary 0 - 100 %

col.trait <- function(out_rasterCMmod,out_makeCM) {
  val<-sort(na.omit(unique(getValues(out_rasterCMmod))))
  cm<- t(out_makeCM)
  n <- matrix(1:length(cm), nrow=nrow(cm), byrow=TRUE)
  cols.plot<-NULL
  for (i in 1:length(val)) {
    cor<-cm[which(n == val[i], arr.ind = T)]
    cols.plot<-c(cols.plot,cor)}
  cor<-cols.plot
} # Selecting colors by gradients
```

```
#####Colors ####

breaks <- 10
#cmat <- makeCM(breaks,"#0096EB", "#820050","grey", "#FFE60F")##mesma do Hidasi
cmat <- makeCM(breaks,"#FFE60F", "#820050","grey", "#0096EB")
plotCM(cmat, "trait cwm/rao range", "trait gap", "")

##### Hmax cwm ####
x.hmax<-raster("raster_hmax_cwm.tif")
y.hmax<-raster("lacuna_hmax.tif")
xy.hmax<-rasterCMmod(x.hmax, y.hmax, 10)
col.hmax<-col.trait(xy.hmax,cmat)
plot(xy.hmax, col=col.hmax, las=1, main="hmax",legend=F)
##### Hmax rao ####
x.hmax_rao<-raster("hmax_rao.tif")
y.hmax<-raster("lacuna_hmax.tif")
xy.hmax_rao<-rasterCMmod(x.hmax_rao, y.hmax, 10)
col.hmax_rao<-col.trait(xy.hmax_rao,cmat)
plot(xy.hmax_rao, col=col.hmax_rao, las=1, main="hmax",legend=F)

##### ssd cwm ####
x. ssd <-raster("raster_ ssd _cwm.tif")
y. ssd <-raster("lacuna_ ssd.tif")
xy. ssd <-rasterCMmod(x. ssd, y. ssd, 10)
col. ssd <-col.trait(xy. ssd,cmat)
plot(xy. ssd, col=col. ssd, las=1, main=" ssd ",legend=F)
##### ssd rao ####
x. ssd _rao<-raster("ssd _rao.tif")
y. ssd <-raster("lacuna_ ssd.tif")
```

```
1.87 xy. ssd _rao<-rasterCMmod(x. ssd _rao, y. ssd, 10)
1.88 col. ssd _rao<-col.trait(xy. ssd _rao,cmat)
1.89 plot(xy. ssd _rao, col=col. ssd _rao, las=1, main=" ssd ",legend=F)
1.90
1.91 ##### sla cwm ####
1.92 x.sla<-raster("raster_sla_cwm.tif")
1.93 y.sla<-raster("lacuna_sla.tif")
1.94 xy.sla<-rasterCMmod(x.sla, y.sla, 10)
1.95 col.sla<-col.trait(xy.sla,cmat)
1.96 plot(xy.sla, col=col.sla, las=1, main="sla",legend=F)
1.97 ##### sla rao ####
1.98 x.sla_rao<-raster("sla_rao.tif")
1.99 y.sla<-raster("lacuna_sla.tif")
1.100 xy.sla_rao<-rasterCMmod(x.sla_rao, y.sla, 10)
1.101 col.sla_rao<-col.trait(xy.sla_rao,cmat)
1.102 plot(xy.sla_rao, col=col.sla_rao, las=1, main="sla",legend=F)
1.103
1.104 ##### sdm cwm ####
1.105 x.sdm<-raster("raster_sdm_cwm.tif")
1.106 y.sdm<-raster("lacuna_sdm.tif")
1.107 xy.sdm<-rasterCMmod(x.sdm, y.sdm, 10)
1.108 col.sdm<-col.trait(xy.sdm,cmat)
1.109 plot(xy.sdm, col=col.sdm, las=1, main="sdm",legend=F)
1.110 ##### sdm rao ####
1.111 x.sdm_rao<-raster("sdm_rao.tif")
1.112 y.sdm<-raster("lacuna_sdm.tif")
1.113 xy.sdm_rao<-rasterCMmod(x.sdm_rao, y.sdm, 10)
1.114 col.sdm_rao<-col.trait(xy.sdm_rao,cmat)
plot(xy.sdm_rao, col=col.sdm_rao, las=1, main="sdm",legend=F, box=F)
```

***Appendix S5***

**Dealing with spatial autocorrelation**

We tested for spatial autocorrelation in the residuals of models with lower AIC values for all traits at geographical level. We calculated Moran's I correlation for different distance classes and chose the distance that spatial autocorrelation was low (I=0.1). Thus, we randomly thinned points by the distance that spatial autocorrelation decreases for each trait and used the thinned points to fit GLM [1], and we repeated this procedure in 500 loops. We further used the resulting estimates and pseudo- $R^2$ to calculate confidence intervals and evaluate if our observed estimates and pseudo - $R^2$ were within it. We used the ncf function from ncf R package to calculate Moran's I correlation coefficients [2].

**Dealing with Type I error**

Since our response variables are mean trait gap values, a spatial pattern could arise by coocuring species in multiple communities, leading to inflated Type I error, that is, it might result in spurious relationships between our response variable and predictor variables [3] Thus, we performed 100 permutations in the trait gap matrix expecting to break coocuring spatial patterns [4]. Then, we tested if coefficients of determination (Pseudo-$R^2$) originated from our models with observed values are higher than expected by chance when compared to the pseudo-$R^2$ originated from resampled values [3,5].

### *Appendix S6*

Table_S 3: Trait gaps for specific leaf area (SLA), maximum height (HMAX), seed dry mass (SDM) and stem specific density (SSD) are signalized as "No" and presence of information is signalized as "Yes".

| Species | SLA | HMAX | SDM | SSD |
|---|---|---|---|---|
| *Abarema brachystachya* | No | Yes | No | Yes |
| *Abarema cochliacarpos* | No | Yes | No | No |
| *Abarema filamentosa* | No | Yes | No | No |
| *Abarema jupunba* | Yes | Yes | No | Yes |
| *Abarema langsdorffii* | No | No | No | No |
| *Abrus precatorius* | Yes | Yes | Yes | No |
| *Abutilon bedfordianum* | No | Yes | No | No |
| *Abutilon rufinerve* | No | Yes | No | No |
| *Acalypha gracilis* | No | Yes | No | No |
| *Acca sellowiana* | Yes | Yes | No | Yes |
| *Achatocarpus praecox* | No | Yes | No | No |
| *Acnistus arborescens* | No | Yes | Yes | No |
| *Acosmium lentiscifolium* | No | Yes | No | Yes |
| *Acrocomia aculeata* | No | Yes | No | No |
| *Actinostemon appendiculatus* | No | Yes | No | No |
| *Actinostemon concolor* | No | Yes | No | No |
| *Actinostemon klotzschii* | No | Yes | No | No |
| *Actinostemon verticillatus* | No | Yes | No | No |
| *Adenocalymma comosum* | No | Yes | No | No |
| *Adenophaedra megalophylla* | No | Yes | No | No |
| *Aegiphila brachiata* | No | Yes | No | No |
| *Aegiphila fluminensis* | No | Yes | No | No |
| *Aegiphila integrifolia* | Yes | Yes | No | Yes |
| *Aegiphila mediterranea* | No | Yes | No | No |
| *Aegiphila obducta* | No | Yes | No | No |
| *Aegiphila pernambucensis* | No | Yes | No | No |
| *Aegiphila verticillata* | No | Yes | No | No |
| *Aegiphila vitelliniflora* | No | Yes | No | No |
| *Agarista eucalyptoides* | No | Yes | No | Yes |
| *Agarista glaberrima* | No | Yes | No | No |
| *Agarista niederleinii* | No | Yes | No | No |

| | | | | |
|---|---|---|---|---|
| *Agarista revoluta* | No | Yes | No | No |
| *Agonandra brasiliensis* | No | Yes | No | Yes |
| *Agonandra excelsa* | No | Yes | No | No |
| *Aiouea acarodomatifera* | No | Yes | No | No |
| *Aiouea saligna* | No | Yes | No | No |
| *Albertinia brasiliensis* | No | Yes | No | No |
| *Albizia edwallii* | No | Yes | No | Yes |
| *Albizia inundata* | No | Yes | No | Yes |
| *Albizia niopoides* | Yes | Yes | Yes | Yes |
| *Albizia pedicellaris* | Yes | Yes | No | Yes |
| *Albizia polycephala* | No | Yes | No | No |
| *Alchornea glandulosa* | Yes | Yes | Yes | Yes |
| *Alchornea sidifolia* | No | Yes | No | Yes |
| *Alchornea triplinervia* | Yes | Yes | Yes | Yes |
| *Algernonia leandrii* | No | Yes | No | No |
| *Alibertia edulis* | Yes | Yes | Yes | Yes |
| *Allagoptera arenaria* | No | Yes | No | No |
| *Allagoptera caudescens* | No | Yes | No | No |
| *Allamanda schottii* | No | Yes | No | No |
| *Allophylus edulis* | Yes | Yes | Yes | Yes |
| *Allophylus guaraniticus* | No | Yes | No | No |
| *Allophylus leucoclados* | No | Yes | No | No |
| *Allophylus petiolulatus* | No | Yes | Yes | Yes |
| *Allophylus puberulus* | No | Yes | No | No |
| *Allophylus semidentatus* | No | Yes | No | No |
| *Allophylus sericeus* | No | Yes | No | Yes |
| *Allophylus strictus* | No | Yes | No | No |
| *Almeidea caerulea* | No | Yes | No | No |
| *Almeidea rubra* | No | Yes | No | No |
| *Aloysia virgata* | No | Yes | Yes | No |
| *Alseis floribunda* | No | Yes | No | Yes |
| *Alseis involuta* | No | Yes | No | No |
| *Alseis pickelii* | No | Yes | No | No |
| *Alsophila capensis* | No | Yes | No | No |
| *Alsophila setosa* | No | Yes | No | No |
| *Alsophila sternbergii* | No | Yes | No | No |
| *Amaioua corymbosa* | Yes | Yes | Yes | No |
| *Amaioua guianensis* | Yes | Yes | No | Yes |

| | | | | |
|---|---|---|---|---|
| *Amaioua intermedia* | Yes | Yes | No | Yes |
| *Amaioua pilosa* | No | Yes | No | No |
| *Amanoa guianensis* | Yes | Yes | No | Yes |
| *Amanoa oblongifolia* | No | Yes | No | Yes |
| *Amburana cearensis* | Yes | Yes | Yes | Yes |
| *Amorimia rigida* | No | Yes | No | No |
| *Amphilophium crucigerum* | No | Yes | No | No |
| *Amphirrhox longifolia* | Yes | Yes | No | Yes |
| *Anacardium occidentale* | Yes | Yes | Yes | Yes |
| *Anadenanthera colubrina* | No | Yes | Yes | Yes |
| *Anadenanthera peregrina* | No | Yes | No | Yes |
| *Anaxagorea dolichocarpa* | No | Yes | No | Yes |
| *Anaxagorea phaeocarpa* | Yes | Yes | No | No |
| *Andira anthelmia* | No | Yes | No | Yes |
| *Andira carvalhoi* | No | Yes | No | No |
| *Andira fraxinifolia* | No | Yes | No | Yes |
| *Andira humilis* | No | Yes | No | No |
| *Andira legalis* | No | Yes | No | Yes |
| *Andira nitida* | No | Yes | No | Yes |
| *Andira ormosioides* | No | Yes | No | No |
| *Andira surinamensis* | Yes | Yes | No | Yes |
| *Andira vermifuga* | No | Yes | No | No |
| *Andradaea floribunda* | No | Yes | No | No |
| *Angostura bracteata* | No | Yes | No | No |
| *Aniba canelilla* | No | Yes | No | Yes |
| *Aniba firmula* | No | Yes | No | Yes |
| *Aniba heringeri* | No | Yes | No | No |
| *Aniba intermedia* | No | Yes | No | No |
| *Aniba viridis* | No | Yes | No | No |
| *Annona acutiflora* | No | Yes | No | No |
| *Annona bahiensis* | No | Yes | No | No |
| *Annona cacans* | No | Yes | No | Yes |
| *Annona crassiflora* | No | Yes | No | Yes |
| *Annona dolabripetala* | No | Yes | No | No |
| *Annona emarginata* | No | Yes | No | Yes |
| *Annona glabra* | Yes | Yes | Yes | Yes |
| *Annona maritima* | No | Yes | No | No |
| *Annona montana* | Yes | Yes | No | No |

| | | | | |
|---|---|---|---|---|
| *Annona mucosa* | No | Yes | No | No |
| *Annona neosericea* | No | Yes | Yes | Yes |
| *Annona pickelii* | No | Yes | No | No |
| *Annona rugulosa* | No | Yes | No | Yes |
| *Annona salzmannii* | No | Yes | No | Yes |
| *Annona sylvatica* | No | Yes | No | No |
| *Anthodiscus amazonicus* | Yes | Yes | No | Yes |
| *Antonia ovata* | Yes | Yes | No | Yes |
| *Aparisthmium cordatum* | Yes | Yes | No | Yes |
| *Apeiba albiflora* | No | Yes | No | Yes |
| *Apeiba tibourbou* | Yes | Yes | Yes | Yes |
| *Aptandra tubicina* | Yes | Yes | No | Yes |
| *Apuleia leiocarpa* | Yes | Yes | No | Yes |
| *Aralia warmingiana* | No | Yes | No | Yes |
| *Arapatiella psilophylla* | No | Yes | No | Yes |
| *Araucaria angustifolia* | Yes | Yes | Yes | Yes |
| *Ardisia guianensis* | Yes | Yes | No | Yes |
| *Asclepias curassavica* | Yes | Yes | No | No |
| *Aspidosperma australe* | No | Yes | No | Yes |
| *Aspidosperma cylindrocarpon* | Yes | Yes | Yes | Yes |
| *Aspidosperma desmanthum* | Yes | Yes | No | Yes |
| *Aspidosperma discolor* | No | Yes | No | Yes |
| *Aspidosperma dispermum* | No | Yes | No | No |
| *Aspidosperma illustre* | No | Yes | No | Yes |
| *Aspidosperma macrocarpon* | No | Yes | No | Yes |
| *Aspidosperma multiflorum* | No | Yes | No | No |
| *Aspidosperma olivaceum* | No | Yes | No | Yes |
| *Aspidosperma parvifolium* | Yes | Yes | No | Yes |
| *Aspidosperma polyneuron* | No | Yes | Yes | Yes |
| *Aspidosperma pyricollum* | No | Yes | No | Yes |
| *Aspidosperma pyrifolium* | Yes | Yes | No | Yes |
| *Aspidosperma ramiflorum* | No | Yes | No | Yes |
| *Aspidosperma riedelii* | No | Yes | No | No |
| *Aspidosperma spruceanum* | Yes | Yes | Yes | Yes |
| *Aspidosperma subincanum* | No | Yes | No | Yes |
| *Aspidosperma tomentosum* | Yes | Yes | No | Yes |
| *Astraea lobata* | No | Yes | No | No |
| *Astrocaryum aculeatisYesum* | No | Yes | No | No |

| | | | | |
|---|---|---|---|---|
| *Astronium concinnum* | No | Yes | No | Yes |
| *Astronium fraxinifolium* | No | Yes | No | Yes |
| *Astronium graveolens* | Yes | Yes | Yes | Yes |
| *Ateleia glazioveana* | No | Yes | No | Yes |
| *Athenaea micrantha* | No | Yes | No | No |
| *Athenaea pogogena* | No | Yes | No | No |
| *Attalea funifera* | No | Yes | No | No |
| *Attalea humilis* | No | Yes | No | No |
| *Augusta longifolia* | No | Yes | No | No |
| *Aureliana fasciculata* | No | Yes | No | No |
| *Aureliana velutina* | No | Yes | No | No |
| *Austrocritonia velutina* | No | Yes | No | No |
| *Austroeupatorium inulaefolium* | No | Yes | No | No |
| *Averrhoidium gardnerianum* | No | Yes | No | No |
| *Azara uruguayensis* | No | Yes | No | No |
| *Baccharis calvescens* | No | Yes | No | No |
| *Baccharis caprariifolia* | No | Yes | No | No |
| *Baccharis dentata* | No | Yes | No | No |
| *Baccharis dracunculifolia* | No | Yes | No | No |
| *Baccharis elaeagnoides* | No | Yes | No | No |
| *Baccharis lateralis* | No | Yes | No | No |
| *Baccharis linearifolia* | No | Yes | No | No |
| *Baccharis oblongifolia* | No | Yes | No | No |
| *Baccharis oxyodonta* | No | Yes | No | No |
| *Baccharis patens* | No | Yes | No | No |
| *Baccharis platypoda* | No | Yes | No | No |
| *Baccharis semiserrata* | No | Yes | No | Yes |
| *Baccharis singularis* | No | Yes | No | No |
| *Baccharis tridentata* | No | Yes | No | No |
| *Baccharis vulneraria* | No | Yes | No | No |
| *Bactris acanthocarpa* | No | Yes | No | No |
| *Bactris bahiensis* | No | Yes | No | No |
| *Bactris caryotifolia* | No | Yes | No | No |
| *Bactris ferruginea* | No | Yes | No | No |
| *Bactris glassmanii* | No | Yes | No | No |
| *Bactris hirta* | No | Yes | No | No |
| *Bactris pickelii* | No | Yes | No | No |
| *Bactris setosa* | No | Yes | No | No |

| | | | | |
|---|---|---|---|---|
| *Bactris vulgaris* | No | Yes | No | No |
| *Bagassa guianensis* | Yes | Yes | No | Yes |
| *Balfourodendron riedelianum* | No | Yes | Yes | Yes |
| *Banara brasiliensis* | No | Yes | No | No |
| *Banara guianensis* | No | Yes | No | Yes |
| *Banara parviflora* | No | Yes | No | No |
| *Banara serrata* | No | Yes | No | No |
| *Banara tomentosa* | No | Yes | No | Yes |
| *Barnebydendron riedelii* | No | Yes | No | No |
| *Basiloxylon brasiliensis* | No | Yes | No | No |
| *Bastardiopsis densiflora* | No | Yes | No | Yes |
| *Bathysa australis* | No | Yes | No | Yes |
| *Bathysa gymnocarpa* | No | Yes | No | No |
| *Bathysa mendoncaei* | Yes | Yes | No | Yes |
| *Bathysa nicholsonii* | No | Yes | No | No |
| *Bathysa stipulata* | No | Yes | No | No |
| *Bauhinia acuruana* | No | Yes | No | No |
| *Bauhinia cheilantha* | No | Yes | No | Yes |
| *Bauhinia dumosa* | No | Yes | No | No |
| *Bauhinia forficata* | No | Yes | Yes | Yes |
| *Bauhinia fusconervis* | No | Yes | No | No |
| *Bauhinia integerrima* | No | Yes | No | No |
| *Bauhinia longifolia* | No | Yes | Yes | Yes |
| *Bauhinia membranacea* | No | Yes | No | No |
| *Bauhinia rufa* | Yes | Yes | No | No |
| *Bauhinia ungulata* | Yes | Yes | Yes | Yes |
| *Beilschmiedia emarginata* | No | Yes | No | Yes |
| *Beilschmiedia taubertiana* | No | Yes | No | No |
| *Berberis laurina* | No | Yes | No | No |
| *Bernardia pulchella* | No | Yes | No | No |
| *Bernardia scabra* | No | Yes | No | No |
| *Bernardinia fluminensis* | No | Yes | No | No |
| *Bixa arborea* | Yes | Yes | No | Yes |
| *Bixa orellana* | Yes | Yes | Yes | Yes |
| *Blanchetiodendron blanchetii* | No | Yes | No | No |
| *Blepharocalyx eggersii* | No | Yes | No | Yes |
| *Blepharocalyx salicifolius* | Yes | Yes | Yes | Yes |
| *Boehmeria caudata* | Yes | Yes | Yes | Yes |

| | | | | |
|---|---|---|---|---|
| *Bonnetia stricta* | No | Yes | No | No |
| *Bougainvillea glabra* | No | Yes | No | No |
| *Bowdichia virgilioides* | Yes | Yes | No | Yes |
| *Brasiliopuntia brasiliensis* | No | Yes | No | No |
| *Bredemeyera brevifolia* | No | Yes | No | No |
| *Bredemeyera disperma* | No | Yes | No | No |
| *Bredemeyera floribunda* | No | Yes | No | No |
| *Bredemeyera laurifolia* | No | Yes | No | No |
| *Brodriguesia santosii* | No | Yes | No | No |
| *BroYesum gaudichaudii* | No | Yes | No | Yes |
| *BroYesum glaziovii* | No | Yes | No | No |
| *BroYesum guianense* | Yes | Yes | Yes | Yes |
| *BroYesum lactescens* | Yes | Yes | No | Yes |
| *BroYesum rubescens* | Yes | Yes | No | Yes |
| *Brunfelsia brasiliensis* | No | Yes | No | No |
| *Brunfelsia clandestina* | No | Yes | No | No |
| *Brunfelsia hydrangeiformis* | No | Yes | No | No |
| *Brunfelsia pauciflora* | No | Yes | Yes | No |
| *Brunfelsia pilosa* | No | Yes | No | No |
| *Brunfelsia uniflora* | No | Yes | No | No |
| *Buchenavia grandis* | Yes | Yes | No | Yes |
| *Buchenavia hoehneana* | No | Yes | No | No |
| *Buchenavia kleinii* | No | Yes | No | No |
| *Buchenavia tetraphylla* | Yes | Yes | No | Yes |
| *Buchenavia tomentosa* | No | Yes | No | No |
| *Bunchosia glandulifera* | No | Yes | No | No |
| *Bunchosia maritima* | No | Yes | No | No |
| *Bunchosia pallescens* | No | Yes | No | No |
| *Butia capitata* | No | Yes | Yes | No |
| *Byrsonima alvimii* | No | Yes | No | No |
| *Byrsonima bahiana* | No | Yes | No | No |
| *Byrsonima blanchetiana* | No | Yes | No | No |
| *Byrsonima cacaophila* | No | Yes | No | No |
| *Byrsonima crassifolia* | Yes | Yes | Yes | Yes |
| *Byrsonima crispa* | Yes | Yes | No | Yes |
| *Byrsonima cydoniifolia* | No | Yes | No | No |
| *Byrsonima gardneriana* | No | Yes | No | No |
| *Byrsonima intermedia* | No | Yes | No | No |

| | | | | |
|---|---|---|---|---|
| *Byrsonima japurensis* | No | Yes | No | Yes |
| *Byrsonima lancifolia* | No | Yes | No | Yes |
| *Byrsonima laxiflora* | No | Yes | No | No |
| *Byrsonima ligustrifolia* | Yes | Yes | No | Yes |
| *Byrsonima myricifolia* | No | Yes | No | No |
| *Byrsonima nitidifolia* | No | Yes | No | No |
| *Byrsonima salzmanniana* | No | Yes | No | No |
| *Byrsonima sericea* | No | Yes | No | Yes |
| *Byrsonima stipulacea* | No | Yes | No | Yes |
| *Byrsonima vacciniifolia* | No | Yes | No | No |
| *Byrsonima variabilis* | No | Yes | No | No |
| *Byrsonima verbascifolia* | No | Yes | No | Yes |
| *Cabralea canjerana* | Yes | Yes | Yes | Yes |
| *Caesalpinia echinata* | No | Yes | No | Yes |
| *Calliandra bella* | No | Yes | No | No |
| *Calliandra brevipes* | No | Yes | No | No |
| *Calliandra foliolosa* | No | Yes | No | No |
| *Calliandra harrisii* | No | Yes | No | No |
| *Calliandra parvifolia* | No | Yes | No | No |
| *Calliandra sessilis* | No | Yes | No | No |
| *Calliandra tweedii* | No | Yes | No | No |
| *Callisthene major* | No | Yes | No | Yes |
| *Callisthene minor* | No | Yes | No | No |
| *Calophyllum brasiliense* | Yes | Yes | Yes | Yes |
| *Calycolpus legrandii* | No | Yes | No | No |
| *Calyptranthes brasiliensis* | No | Yes | No | No |
| *Calyptranthes clusiifolia* | No | Yes | No | Yes |
| *Calyptranthes concinna* | Yes | Yes | No | Yes |
| *Calyptranthes lanceolata* | No | Yes | Yes | No |
| *Calyptranthes lucida* | Yes | Yes | No | Yes |
| *Calyptranthes pulchella* | No | Yes | No | No |
| *Calyptranthes rubella* | No | Yes | No | No |
| *Calyptranthes strigipes* | No | Yes | No | No |
| *Calyptranthes widgreniana* | No | Yes | No | No |
| *Campomanesia adamantium* | No | Yes | No | No |
| *Campomanesia aromatica* | Yes | Yes | No | No |
| *Campomanesia dichotoma* | No | Yes | No | Yes |
| *Campomanesia eugenioides* | No | Yes | No | Yes |

| | | | | |
|---|---|---|---|---|
| *Campomanesia guaviroba* | Yes | Yes | No | Yes |
| *Campomanesia guazumifolia* | No | Yes | No | Yes |
| *Campomanesia ilhoensis* | No | Yes | No | No |
| *Campomanesia laurifolia* | No | Yes | No | Yes |
| *Campomanesia neriiflora* | No | Yes | No | Yes |
| *Campomanesia phaea* | No | Yes | No | No |
| *Campomanesia pubescens* | No | Yes | No | No |
| *Campomanesia reitziana* | No | Yes | No | Yes |
| *Campomanesia rufa* | No | Yes | No | Yes |
| *Campomanesia schlechtendaliana* | No | Yes | No | Yes |
| *Campomanesia velutina* | No | Yes | No | No |
| *Campomanesia xanthocarpa* | Yes | Yes | Yes | Yes |
| *Campuloclinium purpurascens* | No | Yes | No | No |
| *Capparidastrum frondosum* | Yes | Yes | Yes | No |
| *Capsicum annuum* | Yes | Yes | Yes | No |
| *Capsicum flexuosum* | No | Yes | No | No |
| *Capsicum parvifolium* | No | Yes | No | No |
| *Caraipa densifolia* | Yes | Yes | No | Yes |
| *Cariniana estrellensis* | Yes | Yes | Yes | Yes |
| *Cariniana legalis* | No | Yes | Yes | Yes |
| *Carpotroche brasiliensis* | No | Yes | No | No |
| *Caryocar edule* | No | Yes | No | Yes |
| *Casearia aculeata* | Yes | Yes | Yes | Yes |
| *Casearia arborea* | Yes | Yes | Yes | Yes |
| *Casearia bahiensis* | No | Yes | No | No |
| *Casearia commersoniana* | Yes | Yes | Yes | No |
| *Casearia decandra* | Yes | Yes | Yes | Yes |
| *Casearia gossypiosperma* | Yes | Yes | No | Yes |
| *Casearia grandiflora* | Yes | Yes | Yes | Yes |
| *Casearia javitensis* | Yes | Yes | No | Yes |
| *Casearia lasiophylla* | No | Yes | No | Yes |
| *Casearia mariquitensis* | No | Yes | No | No |
| *Casearia obliqua* | Yes | Yes | No | Yes |
| *Casearia oblongifolia* | No | Yes | No | No |
| *Casearia pauciflora* | No | Yes | No | No |
| *Casearia rupestris* | No | Yes | No | No |
| *Casearia selloana* | No | Yes | No | No |
| *Casearia sylvestris* | Yes | Yes | Yes | Yes |

| | | | | |
|---|---|---|---|---|
| *Casearia ulmifolia* | Yes | Yes | No | No |
| *Cassia ferruginea* | No | Yes | No | Yes |
| *Cassia grandis* | No | Yes | Yes | No |
| *Cassia leptophylla* | No | Yes | No | No |
| *Cathedra acuminata* | Yes | Yes | No | No |
| *Cathedra bahiensis* | No | Yes | No | No |
| *Cavanillesia umbellata* | No | Yes | No | Yes |
| *Cecropia glaziovii* | No | Yes | Yes | Yes |
| *Cecropia hololeuca* | No | Yes | Yes | Yes |
| *Cecropia pachystachya* | No | Yes | Yes | Yes |
| *Cecropia palmata* | No | Yes | No | No |
| *Cedrela fissilis* | Yes | Yes | Yes | Yes |
| *Cedrela odorata* | Yes | Yes | Yes | Yes |
| *Ceiba erianthos* | No | Yes | No | Yes |
| *Ceiba glaziovii* | No | Yes | No | Yes |
| *Ceiba speciosa* | Yes | Yes | Yes | Yes |
| *Celtis ehrenbergiana* | No | Yes | Yes | Yes |
| *Celtis iguanaea* | Yes | Yes | Yes | Yes |
| *Centrolobium microchaete* | Yes | Yes | No | Yes |
| *Centrolobium robustum* | No | Yes | No | Yes |
| *Centrolobium sclerophyllum* | No | Yes | No | Yes |
| *Centrolobium tomentosum* | No | Yes | No | Yes |
| *Cephalanthus glabratus* | No | Yes | No | No |
| *Cereus fernambucensis* | No | Yes | No | No |
| *Cereus hildmannianus* | No | Yes | No | No |
| *Cereus jamacaru* | No | Yes | No | No |
| *Cestrum axillare* | No | Yes | No | No |
| *Cestrum bracteatum* | No | Yes | No | No |
| *Cestrum corymbosum* | No | Yes | No | No |
| *Cestrum intermedium* | No | Yes | No | No |
| *Cestrum retrofractum* | No | Yes | No | No |
| *Cestrum salzmannii* | No | Yes | No | No |
| *Cestrum schlechtendalii* | No | Yes | No | No |
| *Cestrum strigilatum* | No | Yes | No | No |
| *Chaetocarpus echinocarpus* | No | Yes | No | Yes |
| *Chaetocarpus myrsinites* | No | Yes | No | No |
| *Chamaecrista apoucouita* | No | Yes | No | Yes |
| *Chamaecrista bahiae* | No | Yes | No | No |

| | | | | |
|---|---|---|---|---|
| *Chamaecrista duartei* | No | Yes | No | No |
| *Chamaecrista ensiformis* | No | Yes | No | No |
| *Chamaecrista ramosa* | No | Yes | No | No |
| *Cheiloclinium cognatum* | Yes | Yes | No | No |
| *Chiococca alba* | No | Yes | No | No |
| *Chionanthus crassifolius* | No | Yes | No | No |
| *Chionanthus filiformis* | No | Yes | No | No |
| *Chionanthus trichotomus* | No | Yes | No | No |
| *Chiropetalum tricoccum* | No | Yes | No | No |
| *Chloroleucon acacioides* | No | Yes | No | No |
| *Chloroleucon dumosum* | No | Yes | No | No |
| *Chloroleucon foliolosum* | No | Yes | No | No |
| *Chloroleucon tortum* | No | Yes | No | No |
| *Chomelia anisomeris* | No | Yes | No | No |
| *Chomelia brasiliana* | No | Yes | No | No |
| *Chomelia intercedens* | No | Yes | No | No |
| *Chomelia obtusa* | No | Yes | No | No |
| *Chomelia parvifolia* | No | Yes | No | No |
| *Chomelia pedunculosa* | No | Yes | No | No |
| *Chomelia pohliana* | No | Yes | No | No |
| *Chomelia pubescens* | No | Yes | No | No |
| *Chromolaena laevigata* | No | Yes | No | No |
| *Chromolaena maximilianii* | No | Yes | No | No |
| *Chromolaena odorata* | No | Yes | Yes | No |
| *Chrysobalanus icaco* | No | Yes | Yes | Yes |
| *Chrysophyllum flexuosum* | Yes | Yes | No | Yes |
| *Chrysophyllum gonocarpum* | No | Yes | Yes | Yes |
| *Chrysophyllum inornatum* | No | Yes | No | Yes |
| *Chrysophyllum lucentifolium* | No | Yes | No | Yes |
| *Chrysophyllum marginatum* | Yes | Yes | Yes | Yes |
| *Chrysophyllum rufum* | No | Yes | No | No |
| *Chrysophyllum splendens* | No | Yes | No | No |
| *Chrysophyllum viride* | No | Yes | Yes | No |
| *Cinnamodendron dinisii* | No | Yes | No | Yes |
| *Cinnamomum amoenum* | No | Yes | No | No |
| *Cinnamomum glaziovii* | No | Yes | No | No |
| *Cinnamomum pseudoglaziovii* | No | Yes | No | No |
| *Cinnamomum sellowianum* | No | Yes | No | No |

| | | | | |
|---|---|---|---|---|
| *Cinnamomum stenophyllum* | No | Yes | No | No |
| *Cinnamomum triplinerve* | Yes | Yes | Yes | Yes |
| *Citharexylum myrianthum* | No | Yes | No | Yes |
| *Citharexylum solanaceum* | No | Yes | No | Yes |
| *Citronella gongonha* | No | Yes | No | Yes |
| *Citronella paniculata* | Yes | Yes | No | Yes |
| *Clarisia ilicifolia* | No | Yes | No | No |
| *Clarisia racemosa* | Yes | Yes | No | Yes |
| *Clavija spinosa* | No | Yes | No | No |
| *Clethra scabra* | Yes | Yes | No | Yes |
| *Clethra uleana* | No | Yes | No | No |
| *Clidemia biserrata* | No | Yes | No | No |
| *Clidemia capilliflora* | No | Yes | No | No |
| *Clidemia capitellata* | No | Yes | No | No |
| *Clidemia debilis* | No | Yes | No | No |
| *Clidemia hirta* | Yes | Yes | No | No |
| *Clidemia sericea* | Yes | Yes | No | No |
| *Clitoria fairchildiana* | No | Yes | No | No |
| *Clusia criuva* | Yes | Yes | No | Yes |
| *Clusia dardanoi* | No | Yes | No | No |
| *Clusia fluminensis* | No | Yes | No | No |
| *Clusia hilariana* | No | No | No | No |
| *Clusia hoffmannseggiana* | No | Yes | No | No |
| *Clusia lanceolata* | No | Yes | No | Yes |
| *Clusia melchiorii* | No | Yes | No | No |
| *Clusia nemorosa* | No | Yes | No | Yes |
| *Clusia panapanari* | No | Yes | Yes | No |
| *Clusia paralicola* | No | Yes | No | No |
| *Clusia spiritu-sanctensis* | No | Yes | No | No |
| *Cnidoscolus oligandrus* | No | Yes | No | No |
| *Cnidoscolus pubescens* | No | Yes | No | Yes |
| *Coccoloba alnifolia* | No | Yes | No | No |
| *Coccoloba declinata* | No | Yes | No | No |
| *Coccoloba densifrons* | Yes | Yes | No | Yes |
| *Coccoloba glaziovii* | No | Yes | No | No |
| *Coccoloba laevis* | No | Yes | No | No |
| *Coccoloba latifolia* | No | Yes | No | Yes |
| *Coccoloba marginata* | No | Yes | No | No |

| | | | | |
|---|---|---|---|---|
| *Coccoloba mollis* | Yes | Yes | Yes | Yes |
| *Coccoloba oblonga* | No | Yes | No | No |
| *Coccoloba ramosisYesa* | No | Yes | No | No |
| *Coccoloba scandens* | No | Yes | No | No |
| *Coccoloba warmingii* | No | Yes | No | No |
| *Cochlospermum regium* | No | Yes | No | No |
| *Cochlospermum vitifolium* | Yes | Yes | Yes | Yes |
| *Colubrina cordifolia* | No | Yes | No | No |
| *Colubrina glandulosa* | Yes | Yes | Yes | Yes |
| *Commiphora leptophloeos* | Yes | Yes | No | Yes |
| *Conchocarpus cuneifolius* | No | Yes | No | No |
| *Conchocarpus cyrtanthus* | No | Yes | No | No |
| *Conchocarpus diadematus* | No | Yes | No | No |
| *Conchocarpus heterophyllus* | No | Yes | No | No |
| *Conchocarpus insignis* | No | Yes | No | No |
| *Conchocarpus longifolius* | No | Yes | No | No |
| *Conchocarpus macrophyllus* | No | Yes | No | No |
| *Condalia buxifolia* | No | Yes | No | No |
| *Connarus blanchetii* | No | Yes | No | No |
| *Connarus detersus* | No | Yes | No | No |
| *Connarus nodosus* | No | Yes | No | No |
| *Connarus portosegurensis* | No | Yes | No | No |
| *Connarus regnellii* | No | Yes | No | No |
| *Connarus rostratus* | No | Yes | No | No |
| *Conocarpus erectus* | No | Yes | No | Yes |
| *Copaifera duckei* | Yes | Yes | No | Yes |
| *Copaifera langsdorffii* | No | Yes | Yes | Yes |
| *Copaifera lucens* | No | Yes | No | No |
| *Copaifera multijuga* | No | Yes | No | Yes |
| *Copaifera trapezifolia* | No | Yes | Yes | Yes |
| *Cordia aberrans* | No | Yes | No | No |
| *Cordia alliodora* | Yes | Yes | Yes | Yes |
| *Cordia americana* | No | Yes | Yes | Yes |
| *Cordia anabaptista* | No | Yes | No | No |
| *Cordia bicolor* | Yes | Yes | Yes | Yes |
| *Cordia ecalyculata* | No | Yes | No | Yes |
| *Cordia exaltata* | Yes | Yes | No | Yes |
| *Cordia lomatoloba* | No | Yes | No | Yes |

| | | | | |
|---|---|---|---|---|
| *Cordia magnoliifolia* | No | Yes | No | Yes |
| *Cordia nodosa* | Yes | Yes | Yes | Yes |
| *Cordia rufescens* | No | Yes | No | No |
| *Cordia sellowiana* | Yes | Yes | No | No |
| *Cordia silvestris* | No | Yes | Yes | Yes |
| *Cordia superba* | No | Yes | Yes | No |
| *Cordia taguahyensis* | No | Yes | No | No |
| *Cordia toqueve* | No | Yes | Yes | No |
| *Cordia trachyphylla* | No | Yes | No | No |
| *Cordia trichoclada* | No | Yes | No | Yes |
| *Cordia trichotoma* | No | Yes | Yes | Yes |
| *Cordiera concolor* | No | Yes | No | No |
| *Cordiera elliptica* | No | Yes | No | No |
| *Cordiera macrophylla* | Yes | Yes | No | Yes |
| *Cordiera myrciifolia* | No | Yes | No | No |
| *Cordiera obtusa* | No | Yes | No | No |
| *Cordiera sessilis* | No | Yes | No | Yes |
| *Cordyline spectabilis* | No | Yes | No | No |
| *Couepia grandiflora* | No | Yes | No | No |
| *Couepia impressa* | No | Yes | No | No |
| *Couepia rufa* | No | Yes | No | No |
| *Couepia schottii* | No | Yes | No | No |
| *Couepia uiti* | No | Yes | No | No |
| *Couepia venosa* | Yes | Yes | No | Yes |
| *Couma rigida* | No | Yes | No | No |
| *Couratari macrosperma* | No | Yes | No | Yes |
| *Coussapoa microcarpa* | No | Yes | No | Yes |
| *Coussapoa pachyphylla* | No | Yes | No | No |
| *Coussarea albescens* | No | Yes | No | No |
| *Coussarea contracta* | No | Yes | Yes | No |
| *Coussarea graciliflora* | No | Yes | No | No |
| *Coussarea hydrangeifolia* | No | Yes | No | Yes |
| *Coussarea ilheotica* | No | Yes | No | No |
| *Coussarea meridionalis* | No | Yes | No | No |
| *Coussarea nodosa* | No | Yes | No | No |
| *Coussarea platyphylla* | No | Yes | No | No |
| *Coutarea hexandra* | Yes | Yes | No | Yes |
| *Crateva tapia* | No | Yes | Yes | Yes |

| | | | | |
|---|---|---|---|---|
| *Critoniopsis quinqueflora* | No | Yes | No | No |
| *Crotalaria vitellina* | No | Yes | Yes | No |
| *Croton astraeatus* | No | Yes | No | No |
| *Croton celtidifolius* | No | Yes | No | Yes |
| *Croton echinocarpus* | No | Yes | No | No |
| *Croton floribundus* | No | Yes | Yes | Yes |
| *Croton heliotropiifolius* | No | Yes | No | No |
| *Croton macrobothrys* | No | Yes | No | No |
| *Croton nepetifolius* | No | Yes | No | No |
| *Croton organensis* | No | Yes | No | No |
| *Croton piptocalyx* | No | Yes | Yes | No |
| *Croton polyandrus* | No | Yes | No | No |
| *Croton salutaris* | No | Yes | No | No |
| *Croton sellowii* | No | Yes | No | No |
| *Croton sincorensis* | No | Yes | No | No |
| *Croton sonderianus* | Yes | Yes | No | Yes |
| *Croton urticifolius* | No | Yes | No | No |
| *Croton urucurana* | No | Yes | Yes | Yes |
| *Cryptocarya aschersoniana* | Yes | Yes | No | Yes |
| *Cryptocarya mandioccana* | No | Yes | No | No |
| *Cryptocarya micrantha* | No | Yes | No | No |
| *Cryptocarya moschata* | No | Yes | Yes | No |
| *Cryptocarya saligna* | No | Yes | No | No |
| *Cupania bracteosa* | No | Yes | No | No |
| *Cupania emarginata* | No | Yes | No | No |
| *Cupania furfuracea* | No | Yes | No | No |
| *Cupania impressinervia* | No | Yes | No | No |
| *Cupania ludowigii* | No | Yes | No | No |
| *Cupania oblongifolia* | No | Yes | Yes | Yes |
| *Cupania paniculata* | No | Yes | No | No |
| *Cupania racemosa* | No | Yes | No | No |
| *Cupania rubiginosa* | No | Yes | No | No |
| *Cupania rugosa* | No | Yes | No | No |
| *Cupania scrobiculata* | Yes | Yes | No | Yes |
| *Cupania vernalis* | Yes | Yes | Yes | Yes |
| *Cupania zanthoxyloides* | No | Yes | No | No |
| *Curatella americana* | Yes | Yes | Yes | Yes |
| *Curitiba prismatica* | No | Yes | No | No |

| | | | | |
|---|---|---|---|---|
| *Cyathea abreviata* | No | Yes | No | No |
| *Cyathea atrovirens* | No | Yes | No | No |
| *Cyathea corcovadensis* | No | Yes | No | No |
| *Cyathea delgadii* | No | Yes | No | No |
| *Cyathea dichromatolepis* | No | Yes | No | No |
| *Cyathea gardneri* | No | Yes | No | No |
| *Cyathea glaziovii* | No | Yes | No | No |
| *Cyathea leucofolis* | No | Yes | No | No |
| *Cyathea microdonta* | No | Yes | No | No |
| *Cyathea phalerata* | No | Yes | No | No |
| *Cyathea praecincta* | No | Yes | No | No |
| *Cyathea villosa* | No | Yes | No | No |
| *Cybianthus amplus* | No | Yes | No | No |
| *Cybianthus brasiliensis* | No | Yes | No | No |
| *Cybianthus cuneifolius* | No | Yes | No | No |
| *Cybianthus densiflorus* | No | Yes | No | No |
| *Cybianthus detergens* | No | No | No | No |
| *Cybianthus fulvopulverulentus* | No | Yes | No | Yes |
| *Cybistax antisyphilitica* | No | Yes | No | Yes |
| *Cyclolobium brasiliense* | No | Yes | No | No |
| *Cymbopetalum brasiliense* | No | Yes | No | Yes |
| *Cynophalla flexuosa* | No | Yes | No | No |
| *Cynophalla hastata* | No | Yes | No | No |
| *Cyrtocarpa caatingae* | No | Yes | No | No |
| *Dahlstedtia pentaphylla* | No | Yes | No | No |
| *Dahlstedtia pinnata* | Yes | Yes | No | Yes |
| *Dalbergia brasiliensis* | No | Yes | No | Yes |
| *Dalbergia ecastaphyllum* | No | Yes | No | No |
| *Dalbergia foliolosa* | No | Yes | No | No |
| *Dalbergia frutescens* | No | Yes | No | Yes |
| *Dalbergia miscolobium* | Yes | Yes | Yes | No |
| *Dalbergia nigra* | No | Yes | No | Yes |
| *Dalbergia villosa* | No | Yes | No | No |
| *Daphnopsis brasiliensis* | No | Yes | No | Yes |
| *Daphnopsis coriacea* | No | Yes | No | No |
| *Daphnopsis fasciculata* | No | Yes | No | No |
| *Daphnopsis martii* | No | Yes | No | No |
| *Daphnopsis racemosa* | No | Yes | No | No |

| | | | | |
|---|---|---|---|---|
| *Daphnopsis schwackeana* | No | Yes | No | No |
| *Daphnopsis sellowiana* | No | Yes | No | No |
| *Dasyphyllum brasiliense* | No | Yes | No | Yes |
| *Dasyphyllum flagellare* | No | Yes | No | No |
| *Dasyphyllum spinescens* | Yes | Yes | No | Yes |
| *Davilla kunthii* | No | Yes | No | No |
| *Davilla macrocarpa* | No | Yes | No | No |
| *Davilla rugosa* | Yes | Yes | No | No |
| *Deguelia costata* | No | Yes | No | No |
| *Dendropanax australis* | No | Yes | No | No |
| *Dendropanax bahiensis* | No | Yes | No | No |
| *Dendropanax cuneatus* | No | Yes | No | No |
| *Dendropanax monogynus* | No | Yes | No | No |
| *Dialium guianense* | Yes | Yes | Yes | Yes |
| *Diatenopteryx sorbifolia* | No | Yes | No | Yes |
| *Dicksonia sellowiana* | No | Yes | No | No |
| *Dictyoloma vandellianum* | No | Yes | No | No |
| *Dilodendron bipinnatum* | No | Yes | No | No |
| *Dimorphandra jorgei* | No | Yes | No | Yes |
| *Dimorphandra mollis* | No | Yes | No | No |
| *Diospyros artanthifolia* | Yes | Yes | No | No |
| *Diospyros brasiliensis* | No | Yes | No | Yes |
| *Diospyros capreifolia* | Yes | Yes | No | Yes |
| *Diospyros hispida* | No | Yes | No | Yes |
| *Diospyros inconstans* | No | Yes | No | Yes |
| *Diospyros sericea* | No | Yes | No | No |
| *Diploon cuspidatum* | Yes | Yes | Yes | Yes |
| *Diplotropis ferruginea* | No | Yes | No | No |
| *Diplotropis incexis* | No | Yes | No | Yes |
| *Diplotropis purpurea* | Yes | Yes | No | Yes |
| *Dipteryx odorata* | Yes | Yes | No | Yes |
| *Discophora guianensis* | Yes | Yes | No | Yes |
| *Dodonaea viscosa* | Yes | Yes | Yes | Yes |
| *Dolichandra unguiculata* | No | Yes | No | No |
| *Drimys angustifolia* | No | Yes | No | Yes |
| *Drimys brasiliensis* | No | Yes | No | Yes |
| *Drypetes sessiliflora* | No | Yes | No | No |
| *Duguetia bahiensis* | No | Yes | No | No |

| | | | | |
|---|---|---|---|---|
| *Duguetia gardneriana* | No | Yes | No | No |
| *Duguetia lanceolata* | No | Yes | No | Yes |
| *Dulacia papillosa* | No | Yes | No | No |
| *Duranta vestita* | No | Yes | No | Yes |
| *Dyssochroma viridiflorum* | No | Yes | No | No |
| *Ecclinusa ramiflora* | Yes | Yes | No | Yes |
| *Emmotum affine* | No | Yes | No | Yes |
| *Emmotum nitens* | No | Yes | No | Yes |
| *Endlicheria glomerata* | No | Yes | No | No |
| *Endlicheria paniculata* | No | Yes | No | Yes |
| *Enterolobium contortisiliquum* | No | Yes | No | Yes |
| *Enterolobium timbouva* | No | Yes | No | Yes |
| *Eremanthus brasiliensis* | No | Yes | No | No |
| *Eremanthus crotonoides* | No | Yes | No | No |
| *Eremanthus erythropappus* | No | Yes | No | Yes |
| *Eremanthus glomerulatus* | No | Yes | No | No |
| *Eremanthus incanus* | No | Yes | No | No |
| *Eriotheca candolleana* | No | Yes | No | Yes |
| *Eriotheca globosa* | Yes | Yes | No | Yes |
| *Eriotheca gracilipes* | No | Yes | No | No |
| *Eriotheca macrophylla* | No | Yes | No | No |
| *Eriotheca obcordata* | No | Yes | No | No |
| *Eriotheca pentaphylla* | Yes | Yes | No | Yes |
| *Erythrina crista-galli* | No | Yes | Yes | Yes |
| *Erythrina falcata* | No | Yes | No | Yes |
| *Erythrina fusca* | Yes | Yes | Yes | Yes |
| *Erythrina poeppigiana* | Yes | Yes | Yes | Yes |
| *Erythrina speciosa* | No | Yes | No | No |
| *Erythrina velutina* | No | Yes | Yes | No |
| *Erythrina verna* | No | Yes | No | No |
| *Erythrochiton brasiliensis* | No | Yes | No | No |
| *Erythroxylum affine* | No | Yes | No | No |
| *Erythroxylum ambiguum* | No | Yes | No | No |
| *Erythroxylum amplifolium* | No | Yes | No | No |
| *Erythroxylum argentinum* | No | Yes | No | Yes |
| *Erythroxylum campestre* | No | Yes | No | No |
| *Erythroxylum citrifolium* | Yes | Yes | No | Yes |
| *Erythroxylum columbinum* | No | Yes | No | No |

| | | | | |
|---|---|---|---|---|
| *Erythroxylum cuneifolium* | No | Yes | No | No |
| *Erythroxylum cuspidifolium* | No | Yes | No | No |
| *Erythroxylum deciduum* | No | Yes | No | Yes |
| *Erythroxylum grandifolium* | No | Yes | No | No |
| *Erythroxylum macrophyllum* | Yes | Yes | No | Yes |
| *Erythroxylum martii* | No | Yes | No | No |
| *Erythroxylum mikanii* | No | Yes | No | No |
| *Erythroxylum mucronatum* | No | Yes | No | No |
| *Erythroxylum ovalifolium* | No | Yes | No | No |
| *Erythroxylum passerinum* | No | Yes | No | No |
| *Erythroxylum pauferrense* | No | Yes | No | No |
| *Erythroxylum pelleterianum* | No | Yes | No | No |
| *Erythroxylum pulchrum* | No | Yes | No | Yes |
| *Erythroxylum revolutum* | No | Yes | No | No |
| *Erythroxylum rimosum* | No | Yes | No | No |
| *Erythroxylum simonis* | No | Yes | No | No |
| *Erythroxylum squamatum* | Yes | Yes | No | Yes |
| *Erythroxylum suberosum* | Yes | Yes | No | No |
| *Erythroxylum subrotundum* | No | Yes | No | No |
| *Erythroxylum subsessile* | No | Yes | No | No |
| *Erythroxylum tenue* | No | Yes | No | No |
| *Erythroxylum vacciniifolium* | No | Yes | No | No |
| *Escallonia bifida* | No | Yes | No | Yes |
| *Eschweilera alvimii* | No | Yes | No | No |
| *Eschweilera apiculata* | No | Yes | No | No |
| *Eschweilera ovata* | No | Yes | No | Yes |
| *Eschweilera tetrapetala* | No | Yes | No | No |
| *Esenbeckia febrifuga* | No | Yes | No | No |
| *Esenbeckia grandiflora* | No | Yes | No | Yes |
| *Esenbeckia leiocarpa* | No | Yes | No | Yes |
| *Eugenia acutata* | No | Yes | Yes | Yes |
| *Eugenia arenaria* | No | Yes | No | No |
| *Eugenia astringens* | No | Yes | No | No |
| *Eugenia aurata* | No | Yes | No | No |
| *Eugenia ayacuchae* | No | Yes | No | No |
| *Eugenia bacopari* | No | Yes | No | No |
| *Eugenia bahiensis* | No | Yes | No | No |
| *Eugenia bimarginata* | No | Yes | No | No |

| | | | | |
|---|---|---|---|---|
| *Eugenia bocainensis* | No | Yes | Yes | No |
| *Eugenia brasiliensis* | No | Yes | Yes | Yes |
| *Eugenia brevistyla* | No | Yes | Yes | No |
| *Eugenia bunchosiifolia* | No | No | No | No |
| *Eugenia burkartiana* | No | Yes | No | No |
| *Eugenia candolleana* | No | Yes | No | Yes |
| *Eugenia capitulifera* | No | Yes | No | No |
| *Eugenia catharinae* | No | Yes | No | No |
| *Eugenia catharinensis* | No | Yes | No | No |
| *Eugenia cerasiflora* | No | Yes | No | Yes |
| *Eugenia cereja* | No | Yes | No | No |
| *Eugenia chlorophylla* | No | Yes | No | No |
| *Eugenia citrifolia* | No | Yes | No | Yes |
| *Eugenia copacabanensis* | No | Yes | No | No |
| *Eugenia cuprea* | No | Yes | Yes | No |
| *Eugenia dodonaeifolia* | No | Yes | No | No |
| *Eugenia dysenterica* | No | Yes | No | Yes |
| *Eugenia egensis* | Yes | Yes | No | No |
| *Eugenia excelsa* | No | Yes | No | No |
| *Eugenia flamingensis* | No | Yes | No | No |
| *Eugenia flavescens* | No | Yes | No | No |
| *Eugenia florida* | Yes | Yes | Yes | Yes |
| *Eugenia francavilleana* | No | Yes | No | No |
| *Eugenia fusca* | No | Yes | No | No |
| *Eugenia gemmiflora* | No | Yes | No | No |
| *Eugenia gracillima* | No | Yes | No | No |
| *Eugenia handroana* | No | Yes | Yes | No |
| *Eugenia handroi* | No | Yes | No | No |
| *Eugenia hiemalis* | No | Yes | No | No |
| *Eugenia hirta* | No | Yes | No | No |
| *Eugenia ilhensis* | No | Yes | No | No |
| *Eugenia involucrata* | No | Yes | Yes | No |
| *Eugenia itapemirimensis* | No | Yes | No | No |
| *Eugenia kleinii* | No | Yes | No | No |
| *Eugenia klotzschiana* | No | Yes | No | No |
| *Eugenia ligustrina* | No | Yes | No | Yes |
| *Eugenia longipedunculata* | No | Yes | No | No |
| *Eugenia luschnathiana* | No | Yes | Yes | No |

| | | | | |
|---|---|---|---|---|
| *Eugenia macrosperma* | No | Yes | No | No |
| *Eugenia magnifica* | No | Yes | No | No |
| *Eugenia malacantha* | No | Yes | No | No |
| *Eugenia mansoi* | No | Yes | No | No |
| *Eugenia melanogyna* | No | Yes | Yes | No |
| *Eugenia modesta* | No | Yes | No | No |
| *Eugenia monosperma* | No | Yes | No | Yes |
| *Eugenia mosenii* | No | Yes | Yes | No |
| *Eugenia multicostata* | No | Yes | Yes | Yes |
| *Eugenia myrcianthes* | No | Yes | No | No |
| *Eugenia neoglomerata* | No | Yes | Yes | No |
| *Eugenia neomyrtifolia* | No | Yes | No | No |
| *Eugenia neoverrucosa* | No | Yes | Yes | No |
| *Eugenia nutans* | No | Yes | No | No |
| *Eugenia oblongata* | No | Yes | Yes | No |
| *Eugenia pisiformis* | No | Yes | No | No |
| *Eugenia platyphylla* | No | Yes | No | No |
| *Eugenia platysema* | No | Yes | No | No |
| *Eugenia pluriflora* | No | Yes | No | Yes |
| *Eugenia prasina* | No | Yes | No | No |
| *Eugenia pruinosa* | No | Yes | No | No |
| *Eugenia pruniformis* | No | Yes | No | No |
| *Eugenia punicifolia* | No | Yes | Yes | No |
| *Eugenia pyriformis* | No | Yes | No | Yes |
| *Eugenia ramboi* | No | Yes | No | No |
| *Eugenia repanda* | No | Yes | No | No |
| *Eugenia rostrata* | No | Yes | No | No |
| *Eugenia rostrifolia* | No | Yes | No | Yes |
| *Eugenia schottiana* | No | Yes | No | No |
| *Eugenia selloi* | No | Yes | No | No |
| *Eugenia sonderiana* | No | Yes | No | Yes |
| *Eugenia speciosa* | No | Yes | No | Yes |
| *Eugenia sphenophylla* | No | Yes | No | No |
| *Eugenia splendens* | No | Yes | No | No |
| *Eugenia stictopetala* | No | Yes | No | No |
| *Eugenia stigmatosa* | No | Yes | No | No |
| *Eugenia subavenia* | No | Yes | No | No |
| *Eugenia suberosa* | No | Yes | No | No |

| | | | | |
|---|---|---|---|---|
| *Eugenia subterminalis* | No | Yes | No | No |
| *Eugenia sulcata* | No | Yes | No | Yes |
| *Eugenia supraaxillaris* | No | Yes | No | No |
| *Eugenia ternatifolia* | No | Yes | No | No |
| *Eugenia umbellata* | No | Yes | No | No |
| *Eugenia umbrosa* | No | Yes | No | No |
| *Eugenia uniflora* | Yes | Yes | Yes | Yes |
| *Eugenia uruguayensis* | Yes | Yes | No | No |
| *Eugenia vattimoana* | No | Yes | No | No |
| *Eugenia verticillata* | No | Yes | No | No |
| *Euplassa cantareirae* | No | Yes | No | Yes |
| *Euplassa legalis* | No | Yes | No | No |
| *Euterpe edulis* | Yes | Yes | Yes | No |
| *Exostyles godoyensis* | No | Yes | No | No |
| *Exostyles venusta* | No | Yes | No | No |
| *Faramea axilliflora* | No | Yes | No | No |
| *Faramea coerulea* | No | Yes | No | No |
| *Faramea corymbosa* | No | Yes | No | No |
| *Faramea hyacinthina* | No | Yes | No | No |
| *Faramea latifolia* | No | Yes | No | No |
| *Faramea martiana* | No | Yes | No | No |
| *Faramea monantha* | No | Yes | No | No |
| *Faramea montevidensis* | No | Yes | No | No |
| *Faramea multiflora* | Yes | Yes | No | No |
| *Faramea nigrescens* | No | Yes | No | No |
| *Faramea occidentalis* | Yes | Yes | Yes | Yes |
| *Faramea oligantha* | No | Yes | No | No |
| *Faramea pachyantha* | Yes | Yes | No | Yes |
| *Faramea truncata* | No | Yes | No | No |
| *Ferdinandusa elliptica* | No | Yes | No | Yes |
| *Ferdinandusa speciosa* | No | Yes | No | No |
| *Ficus adhatodifolia* | No | Yes | No | Yes |
| *Ficus calyptroceras* | No | Yes | No | No |
| *Ficus castellviana* | No | Yes | No | No |
| *Ficus cestrifolia* | No | Yes | No | No |
| *Ficus citrifolia* | Yes | Yes | Yes | Yes |
| *Ficus clusiifolia* | No | Yes | No | Yes |
| *Ficus cyclophylla* | No | Yes | No | No |

| | | | | |
|---|---|---|---|---|
| *Ficus enormis* | No | Yes | No | Yes |
| *Ficus eximia* | No | Yes | No | Yes |
| *Ficus gomelleira* | No | Yes | Yes | Yes |
| *Ficus guaranitica* | No | Yes | No | No |
| *Ficus hirsuta* | No | Yes | No | Yes |
| *Ficus insipida* | Yes | Yes | Yes | Yes |
| *Ficus luschnathiana* | No | Yes | No | Yes |
| *Ficus mariae* | No | Yes | No | No |
| *Ficus maxima* | Yes | Yes | No | Yes |
| *Ficus mexiae* | No | Yes | No | Yes |
| *Ficus nymphaeifolia* | Yes | Yes | No | Yes |
| *Ficus obtusifolia* | Yes | Yes | Yes | Yes |
| *Ficus obtusiuscula* | No | Yes | No | No |
| *Ficus organensis* | No | Yes | No | Yes |
| *Ficus pertusa* | Yes | Yes | No | Yes |
| *Ficus pulchella* | No | Yes | No | No |
| *Ficus trigona* | Yes | Yes | No | No |
| *Ficus trigonata* | Yes | Yes | Yes | Yes |
| *Forsteronia leptocarpa* | No | Yes | No | No |
| *Forsteronia pubescens* | No | Yes | No | No |
| *Forsteronia rufa* | No | Yes | No | No |
| *Galipea jasminiflora* | No | Yes | No | No |
| *Galipea laxiflora* | No | Yes | No | No |
| *Gallesia integrifolia* | Yes | Yes | No | Yes |
| *Gamochaeta americana* | No | Yes | No | No |
| *Garcinia brasiliensis* | Yes | Yes | Yes | Yes |
| *Garcinia gardneriana* | No | Yes | Yes | Yes |
| *Garcinia macrophylla* | Yes | Yes | Yes | Yes |
| *Gaylussacia brasiliensis* | No | Yes | No | No |
| *Geissanthus ambiguus* | No | Yes | No | No |
| *Geissospermum laeve* | Yes | Yes | No | Yes |
| *Genipa americana* | Yes | Yes | Yes | Yes |
| *Geoffroea spinosa* | No | Yes | No | Yes |
| *Geonoma brevispatha* | No | Yes | No | No |
| *Geonoma gamiova* | No | Yes | No | No |
| *Geonoma pauciflora* | No | Yes | No | No |
| *Geonoma pohliana* | No | Yes | No | No |
| *Geonoma schottiana* | No | Yes | No | No |

| | | | | |
|---|---|---|---|---|
| *Glycydendron amazonicum* | Yes | Yes | No | Yes |
| *Gochnatia oligocephala* | No | Yes | No | No |
| *Gochnatia paniculata* | No | Yes | No | No |
| *Gochnatia polymorpha* | No | Yes | No | Yes |
| *Gochnatia pulchra* | No | Yes | No | No |
| *Godmania dardanoi* | No | Yes | No | No |
| *Goniorrhachis marginata* | No | Yes | No | Yes |
| *Gorceixia decurrens* | No | Yes | No | No |
| *Grazielia serrata* | No | Yes | No | No |
| *Grazielodendron rio-docensis* | No | Yes | No | Yes |
| *Griselinia ruscifolia* | No | Yes | No | No |
| *Guapira areolata* | Yes | Yes | No | No |
| *Guapira graciliflora* | No | Yes | No | No |
| *Guapira hirsuta* | No | Yes | No | No |
| *Guapira laxiflora* | No | Yes | No | No |
| *Guapira nitida* | No | Yes | No | No |
| *Guapira noxia* | Yes | Yes | No | No |
| *Guapira obtusata* | No | Yes | No | No |
| *Guapira opposita* | Yes | Yes | Yes | Yes |
| *Guapira pernambucensis* | No | Yes | No | No |
| *Guapira tomentosa* | No | Yes | No | No |
| *Guapira venosa* | No | Yes | No | No |
| *Guarea blanchetii* | No | Yes | No | No |
| *Guarea guidonia* | Yes | Yes | Yes | Yes |
| *Guarea kunthiana* | Yes | Yes | No | Yes |
| *Guarea macrophylla* | Yes | Yes | No | Yes |
| *Guatteria australis* | Yes | Yes | Yes | No |
| *Guatteria campestris* | No | Yes | No | No |
| *Guatteria candolleana* | No | Yes | No | No |
| *Guatteria ferruginea* | No | Yes | No | No |
| *Guatteria macropus* | No | Yes | No | No |
| *Guatteria oligocarpa* | No | Yes | No | No |
| *Guatteria pogonopus* | No | Yes | No | No |
| *Guatteria pohliana* | No | Yes | No | No |
| *Guatteria schomburgkiana* | Yes | Yes | No | Yes |
| *Guatteria sellowiana* | No | Yes | No | No |
| *Guatteria villosisYesa* | No | Yes | No | No |
| *Guazuma crinita* | No | Yes | No | Yes |

| | | | | |
|---|---|---|---|---|
| *Guazuma ulmifolia* | Yes | Yes | Yes | Yes |
| *Guettarda angelica* | No | Yes | No | No |
| *Guettarda platyphylla* | No | Yes | No | No |
| *Guettarda platypoda* | No | Yes | No | No |
| *Guettarda pohliana* | No | Yes | No | No |
| *Guettarda sericea* | No | Yes | No | No |
| *Guettarda uruguensis* | No | Yes | Yes | Yes |
| *Guettarda viburnoides* | No | Yes | No | No |
| *Guilandina bonduc* | No | Yes | No | No |
| *Gustavia augusta* | Yes | Yes | No | Yes |
| *Gymnanthes gaudichaudii* | No | Yes | No | No |
| *Gymnanthes glabrata* | No | Yes | No | No |
| *Hamelia patens* | Yes | Yes | Yes | No |
| *Hancornia speciosa* | No | Yes | No | No |
| *Handroanthus albus* | No | Yes | No | Yes |
| *Handroanthus chrysotrichus* | No | Yes | No | Yes |
| *Handroanthus heptaphyllus* | No | Yes | No | Yes |
| *Handroanthus impetiginosus* | Yes | Yes | Yes | Yes |
| *Handroanthus ochraceus* | No | Yes | No | Yes |
| *Handroanthus pulcherrimus* | No | Yes | No | No |
| *Handroanthus serratifolius* | No | Yes | No | Yes |
| *Handroanthus umbellatus* | No | Yes | No | No |
| *Handroanthus vellosoi* | No | Yes | No | Yes |
| *Harleyodendron unifoliolatum* | No | Yes | No | Yes |
| *Hedyosmum brasiliense* | No | Yes | No | Yes |
| *Heisteria perianthomega* | No | Yes | No | No |
| *Heisteria silvianii* | No | Yes | Yes | Yes |
| *Helicostylis tomentosa* | Yes | Yes | No | Yes |
| *Helicteres brevispira* | No | Yes | No | No |
| *Helicteres eichleri* | No | Yes | No | No |
| *Helicteres guazumifolia* | No | Yes | No | No |
| *Helicteres heptandra* | No | Yes | No | No |
| *Helicteres ovata* | No | Yes | No | No |
| *Helicteres pentandra* | No | Yes | No | No |
| *Helietta apiculata* | No | Yes | No | Yes |
| *Heliocarpus popayanensis* | Yes | Yes | Yes | Yes |
| *Hennecartia omphalandra* | No | Yes | No | Yes |
| *Henriettea glabra* | No | Yes | No | No |

| | | | | |
|---|---|---|---|---|
| *Henriettea saldanhaei* | No | Yes | No | No |
| *Henriettea succosa* | No | Yes | No | No |
| *Heterocondylus alatus* | No | Yes | No | No |
| *Heterocondylus vitalbae* | Yes | Yes | No | No |
| *Hevea brasiliensis* | Yes | Yes | Yes | Yes |
| *Hieronyma alchorneoides* | Yes | Yes | Yes | Yes |
| *Hieronyma oblonga* | Yes | Yes | No | Yes |
| *Himatanthus articulatus* | Yes | Yes | No | Yes |
| *Himatanthus bracteatus* | No | Yes | No | Yes |
| *Himatanthus obovatus* | No | Yes | No | Yes |
| *Himatanthus phagedaenicus* | No | Yes | No | Yes |
| *Hirtella angustifolia* | No | Yes | No | No |
| *Hirtella bahiensis* | No | Yes | No | No |
| *Hirtella bicornis* | Yes | Yes | No | Yes |
| *Hirtella ciliata* | No | Yes | No | Yes |
| *Hirtella glandulosa* | Yes | Yes | No | Yes |
| *Hirtella gracilipes* | No | Yes | No | No |
| *Hirtella hebeclada* | Yes | Yes | No | Yes |
| *Hirtella martiana* | No | Yes | No | No |
| *Hirtella racemosa* | Yes | Yes | Yes | Yes |
| *Hirtella triandra* | Yes | Yes | Yes | Yes |
| *Holocalyx balansae* | No | Yes | No | Yes |
| *Hornschuchia bryotrophe* | No | Yes | No | No |
| *Hornschuchia myrtillus* | No | Yes | No | No |
| *Hortia brasiliana* | No | Yes | No | No |
| *Huberia ovalifolia* | No | Yes | No | No |
| *Humiria balsamifera* | Yes | Yes | No | Yes |
| *Humiriastrum dentatum* | No | Yes | No | Yes |
| *Humiriastrum glaziovii* | No | Yes | No | No |
| *Humiriastrum spiritu-sancti* | No | Yes | No | No |
| *Hydrogaster trinervis* | No | Yes | No | Yes |
| *Hymenaea aurea* | No | Yes | No | No |
| *Hymenaea courbaril* | Yes | Yes | Yes | Yes |
| *Hymenaea martiana* | No | Yes | No | Yes |
| *Hymenaea oblongifolia* | Yes | Yes | No | Yes |
| *Hymenaea rubriflora* | No | Yes | No | No |
| *Hymenaea stigonocarpa* | No | Yes | No | Yes |
| *Hymenolobium alagoanum* | No | Yes | No | No |

| | | | | |
|---|---|---|---|---|
| *Hymenolobium janeirense* | No | Yes | No | No |
| *Hyptidendron asperrimum* | No | Yes | No | Yes |
| *Ilex brevicuspis* | Yes | Yes | No | Yes |
| *Ilex cerasifolia* | No | Yes | No | Yes |
| *Ilex chamaedryfolia* | No | Yes | No | No |
| *Ilex conocarpa* | No | Yes | No | No |
| *Ilex dumosa* | Yes | Yes | No | No |
| *Ilex integerrima* | No | Yes | No | No |
| *Ilex lundii* | No | Yes | No | No |
| *Ilex microdonta* | Yes | Yes | No | No |
| *Ilex paraguariensis* | Yes | Yes | No | Yes |
| *Ilex psammophila* | No | Yes | No | No |
| *Ilex pseudobuxus* | No | Yes | No | No |
| *Ilex taubertiana* | No | Yes | No | No |
| *Ilex theezans* | No | Yes | No | Yes |
| *Indigofera suffruticosa* | No | Yes | Yes | No |
| *Inga barbata* | No | Yes | No | No |
| *Inga blanchetiana* | No | Yes | No | No |
| *Inga bullata* | No | Yes | No | No |
| *Inga capitata* | Yes | Yes | No | Yes |
| *Inga cayennensis* | Yes | Yes | No | Yes |
| *Inga cylindrica* | Yes | Yes | No | Yes |
| *Inga edulis* | Yes | Yes | Yes | Yes |
| *Inga edwallii* | No | Yes | No | No |
| *Inga exfoliata* | No | Yes | No | No |
| *Inga flagelliformis* | No | Yes | No | No |
| *Inga gracilifolia* | Yes | Yes | No | No |
| *Inga grazielae* | No | Yes | No | No |
| *Inga heterophylla* | Yes | Yes | No | Yes |
| *Inga hispida* | No | Yes | No | No |
| *Inga ingoides* | No | Yes | No | Yes |
| *Inga laurina* | Yes | Yes | Yes | Yes |
| *Inga lenticellata* | No | Yes | No | No |
| *Inga lentiscifolia* | No | Yes | No | No |
| *Inga leptantha* | No | Yes | No | No |
| *Inga marginata* | Yes | Yes | Yes | Yes |
| *Inga maritima* | No | Yes | No | No |
| *Inga pleiogyna* | No | Yes | No | No |

| | | | | |
|---|---|---|---|---|
| *Inga sellowiana* | No | Yes | No | No |
| *Inga sessilis* | Yes | Yes | No | Yes |
| *Inga striata* | Yes | Yes | No | Yes |
| *Inga subnuda* | Yes | Yes | No | Yes |
| *Inga tenuis* | No | Yes | No | No |
| *Inga thibaudiana* | Yes | Yes | Yes | Yes |
| *Inga vera* | Yes | Yes | No | Yes |
| *Inga virescens* | No | Yes | No | Yes |
| *Inga vulpina* | No | Yes | No | No |
| *Ixora brevifolia* | No | Yes | No | Yes |
| *Ixora gardneriana* | No | Yes | No | Yes |
| *Ixora muelleri* | No | Yes | No | No |
| *Ixora venulosa* | No | Yes | No | No |
| *Jacaranda bracteata* | No | Yes | No | No |
| *Jacaranda caroba* | No | Yes | No | No |
| *Jacaranda cuspidifolia* | No | Yes | No | No |
| *Jacaranda jasminoides* | No | Yes | No | No |
| *Jacaranda macrantha* | No | Yes | No | No |
| *Jacaranda micrantha* | No | Yes | No | Yes |
| *Jacaranda obovata* | No | Yes | No | No |
| *Jacaranda puberula* | Yes | Yes | No | Yes |
| *Jacaratia heptaphylla* | No | Yes | No | No |
| *Jacaratia spinosa* | Yes | Yes | Yes | Yes |
| *Jacquinia armillaris* | Yes | Yes | No | No |
| *Jatropha mollissima* | No | Yes | No | No |
| *Jatropha mutabilis* | No | Yes | No | No |
| *Joannesia princeps* | No | Yes | No | Yes |
| *Jodina rhombifolia* | No | Yes | No | Yes |
| *Kaunia rufescens* | No | Yes | No | No |
| *Kielmeyera albopunctata* | No | Yes | No | No |
| *Kielmeyera coriacea* | Yes | Yes | No | No |
| *Kielmeyera lathrophyton* | No | Yes | No | Yes |
| *Kielmeyera membranacea* | No | Yes | No | No |
| *Kielmeyera neglecta* | No | Yes | No | No |
| *Kielmeyera petiolaris* | No | Yes | No | No |
| *Kielmeyera rubriflora* | No | Yes | No | Yes |
| *Kielmeyera rugosa* | No | Yes | No | No |
| *Lacistema aggregatum* | Yes | Yes | Yes | Yes |

| | | | | |
|---|---|---|---|---|
| *Lacistema hasslerianum* | No | Yes | No | No |
| *Lacistema lucidum* | No | Yes | No | No |
| *Lacistema pubescens* | Yes | Yes | No | Yes |
| *Lacistema robustum* | No | Yes | No | No |
| *Lacmellea aculeata* | Yes | Yes | No | No |
| *Lacunaria crenata* | Yes | Yes | No | Yes |
| *Ladenbergia hexandra* | No | Yes | No | No |
| *Lafoensia glyptocarpa* | No | Yes | No | Yes |
| *Lafoensia pacari* | No | Yes | No | Yes |
| *Lafoensia vandelliana* | No | Yes | No | No |
| *Laguncularia racemosa* | No | Yes | No | Yes |
| *Lamanonia ternata* | No | Yes | No | Yes |
| *Lantana camara* | Yes | Yes | Yes | Yes |
| *Lantana fucata* | No | Yes | No | No |
| *Lantana radula* | No | Yes | No | No |
| *Lantana undulata* | No | Yes | No | No |
| *Laplacea fructicosa* | No | Yes | No | No |
| *Leandra acutiflora* | No | Yes | No | No |
| *Leandra aurea* | No | Yes | No | No |
| *Leandra australis* | No | Yes | No | No |
| *Leandra barbinervis* | No | Yes | No | No |
| *Leandra bergiana* | No | Yes | No | No |
| *Leandra carassana* | No | Yes | No | No |
| *Leandra clidemioides* | No | Yes | No | No |
| *Leandra cuneata* | No | Yes | No | No |
| *Leandra dubia* | No | Yes | No | No |
| *Leandra fragilis* | No | Yes | No | No |
| *Leandra gardneriana* | No | Yes | No | No |
| *Leandra glazioviana* | No | Yes | No | No |
| *Leandra ionopogon* | No | Yes | No | No |
| *Leandra laevigata* | No | Yes | No | No |
| *Leandra melastomoides* | No | Yes | No | No |
| *Leandra nianga* | No | Yes | No | No |
| *Leandra regnellii* | No | Yes | No | No |
| *Leandra reversa* | No | Yes | No | No |
| *Leandra rhamnifolia* | No | Yes | No | No |
| *Leandra rufescens* | No | Yes | No | No |
| *Leandra sericea* | No | Yes | No | No |

| | | | | |
|---|---|---|---|---|
| *Leandra variabilis* | No | Yes | No | No |
| *Lecythis lanceolata* | No | Yes | No | Yes |
| *Lecythis lurida* | Yes | Yes | No | Yes |
| *Lecythis pisonis* | Yes | Yes | No | Yes |
| *Lepidaploa argyrotricha* | No | Yes | No | No |
| *Lepidaploa balansae* | No | Yes | No | No |
| *Lepidaploa chalybaea* | No | Yes | No | No |
| *Lepidaploa cotoneaster* | No | Yes | No | No |
| *Leptolobium bijugum* | No | Yes | No | No |
| *Leptolobium dasycarpum* | No | Yes | No | No |
| *Leptolobium elegans* | No | Yes | No | No |
| *Lessingianthus cataractarum* | No | Yes | No | No |
| *Lessingianthus macrophyllus* | No | Yes | No | No |
| *Leucochloron incuriale* | No | Yes | No | No |
| *Libidibia ferrea* | No | Yes | No | No |
| *Licania discolor* | No | Yes | No | Yes |
| *Licania heteromorpha* | Yes | Yes | Yes | Yes |
| *Licania hoehnei* | Yes | Yes | No | Yes |
| *Licania hypoleuca* | Yes | Yes | Yes | Yes |
| *Licania kunthiana* | Yes | Yes | No | Yes |
| *Licania leptostachya* | No | Yes | No | No |
| *Licania littoralis* | No | Yes | No | Yes |
| *Licania micrantha* | Yes | Yes | No | Yes |
| *Licania octandra* | Yes | Yes | No | Yes |
| *Licania rigida* | No | Yes | No | Yes |
| *Licania spicata* | No | Yes | No | No |
| *Licania tomentosa* | No | Yes | No | Yes |
| *Licaria armeniaca* | Yes | Yes | No | Yes |
| *Licaria bahiana* | No | Yes | No | No |
| *Licaria chrysophylla* | Yes | Yes | No | Yes |
| *Licaria guianensis* | Yes | Yes | No | Yes |
| *Lippia brasiliensis* | No | Yes | No | No |
| *Lithrea brasiliensis* | No | Yes | No | No |
| *Lithrea molleoides* | No | Yes | No | No |
| *Lonchocarpus campestris* | Yes | Yes | No | Yes |
| *Lonchocarpus cultratus* | No | Yes | No | No |
| *Lonchocarpus muehlbergianus* | No | Yes | No | Yes |
| *Lonchocarpus nitidus* | No | Yes | No | Yes |

| | | | | |
|---|---|---|---|---|
| *Lonchocarpus sericeus* | Yes | Yes | Yes | Yes |
| *Lonchocarpus subglaucescens* | No | Yes | No | No |
| *Luehea candicans* | No | Yes | No | No |
| *Luehea cymulosa* | No | Yes | No | Yes |
| *Luehea divaricata* | Yes | Yes | No | Yes |
| *Luehea grandiflora* | No | Yes | No | No |
| *Luehea ochrophylla* | No | Yes | No | No |
| *Luehea paniculata* | No | Yes | No | Yes |
| *Luehea speciosa* | Yes | Yes | Yes | Yes |
| *Luetzelburgia guaissara* | No | Yes | No | No |
| *Mabea fistulifera* | No | Yes | No | No |
| *Mabea glaziovii* | No | Yes | No | No |
| *Mabea occidentalis* | Yes | Yes | Yes | Yes |
| *Mabea piriri* | Yes | Yes | No | Yes |
| *Machaerium acutifolium* | Yes | Yes | No | Yes |
| *Machaerium fulvovenosum* | No | Yes | No | Yes |
| *Machaerium hatschbachii* | No | Yes | No | No |
| *Machaerium incorruptibile* | No | Yes | No | Yes |
| *Machaerium nyctitans* | No | Yes | No | No |
| *Machaerium paraguariense* | No | Yes | No | Yes |
| *Machaerium pedicellatum* | No | Yes | No | No |
| *Machaerium punctatum* | No | Yes | No | No |
| *Machaerium salzmannii* | No | Yes | No | No |
| *Machaerium scleroxylon* | No | Yes | No | Yes |
| *Machaerium stipitatum* | No | Yes | No | Yes |
| *Machaerium villosum* | No | Yes | No | Yes |
| *Machaonia acuminata* | No | Yes | No | No |
| *Maclura tinctoria* | Yes | Yes | No | Yes |
| *Macoubea guianensis* | Yes | Yes | No | Yes |
| *Macrocarpaea obtusifolia* | No | Yes | No | No |
| *Macrolobium latifolium* | No | Yes | No | No |
| *Macrolobium rigidum* | No | Yes | No | No |
| *Macropeplus dentatus* | No | Yes | No | No |
| *Macropeplus ligustrinus* | No | Yes | No | No |
| *Macrothumia kuhlmannii* | No | Yes | No | No |
| *Macrotorus utriculatus* | No | Yes | No | No |
| *Magnolia ovata* | No | Yes | Yes | Yes |
| *Malouetia arborea* | No | Yes | No | No |

| | | | | |
|---|---|---|---|---|
| *Manihot anomala* | No | Yes | No | No |
| *Manihot caerulescens* | No | Yes | No | No |
| *Manihot carthaginensis* | No | Yes | No | No |
| *Manihot grahamii* | No | Yes | Yes | No |
| *Manihot pilosa* | No | Yes | No | No |
| *Manihot tripartita* | No | Yes | No | No |
| *Manilkara longifolia* | No | Yes | No | No |
| *Manilkara maxima* | No | Yes | No | No |
| *Manilkara rufula* | No | Yes | No | No |
| *Manilkara salzmannii* | No | Yes | No | Yes |
| *Manilkara subsericea* | No | Yes | No | Yes |
| *Maprounea brasiliensis* | No | Yes | No | No |
| *Maprounea guianensis* | Yes | Yes | No | Yes |
| *Margaritaria nobilis* | Yes | Yes | Yes | Yes |
| *Margaritopsis astrellantha* | No | Yes | No | No |
| *Margaritopsis cephalantha* | No | Yes | No | No |
| *Margaritopsis chaenotricha* | No | Yes | No | No |
| *Margaritopsis cymuligera* | No | Yes | No | No |
| *Marlierea clausseniana* | No | Yes | No | No |
| *Marlierea eugenioides* | No | Yes | No | No |
| *Marlierea eugeniopsoides* | No | Yes | Yes | No |
| *Marlierea excoriata* | No | Yes | No | No |
| *Marlierea laevigata* | No | Yes | No | No |
| *Marlierea neuwiedeana* | No | Yes | No | No |
| *Marlierea obscura* | Yes | Yes | Yes | Yes |
| *Marlierea obversa* | No | Yes | No | No |
| *Marlierea racemosa* | No | Yes | No | No |
| *Marlierea regeliana* | No | Yes | No | No |
| *Marlierea reitzii* | No | Yes | No | No |
| *Marlierea silvatica* | No | Yes | No | Yes |
| *Marlierea suaveolens* | No | Yes | Yes | No |
| *Marlierea tomentosa* | Yes | Yes | Yes | Yes |
| *Martiodendron mediterraneum* | No | Yes | No | No |
| *Matayba arborescens* | Yes | Yes | No | Yes |
| *Matayba discolor* | No | Yes | No | Yes |
| *Matayba elaeagnoides* | Yes | Yes | Yes | Yes |
| *Matayba guianensis* | Yes | Yes | No | Yes |
| *Matayba inelegans* | Yes | Yes | No | Yes |

| | | | | |
|---|---|---|---|---|
| *Matayba intermedia* | No | Yes | No | Yes |
| *Matayba juglandifolia* | No | Yes | No | No |
| *Matayba mollis* | No | Yes | No | No |
| *Mattfeldanthus andrade-limae* | No | Yes | No | No |
| *Mauritia flexuosa* | No | Yes | No | No |
| *Maytenus ardisiifolia* | No | Yes | No | No |
| *Maytenus boaria* | No | Yes | Yes | Yes |
| *Maytenus brasiliensis* | No | Yes | No | No |
| *Maytenus cassineformis* | No | Yes | No | No |
| *Maytenus cestrifolia* | No | Yes | No | No |
| *Maytenus communis* | No | Yes | No | No |
| *Maytenus dasyclada* | No | Yes | No | No |
| *Maytenus distichophylla* | No | Yes | No | No |
| *Maytenus erythroxyla* | No | Yes | No | No |
| *Maytenus evonymoides* | No | Yes | No | No |
| *Maytenus floribunda* | No | Yes | No | No |
| *Maytenus glaucescens* | No | Yes | No | No |
| *Maytenus gonoclada* | No | Yes | Yes | Yes |
| *Maytenus ilicifolia* | No | Yes | No | Yes |
| *Maytenus littoralis* | No | Yes | No | No |
| *Maytenus obtusifolia* | No | Yes | No | Yes |
| *Maytenus opaca* | No | Yes | No | No |
| *Maytenus patens* | No | Yes | No | No |
| *Maytenus rigida* | No | Yes | No | Yes |
| *Maytenus schumanniana* | No | Yes | No | No |
| *Melanopsidium nigrum* | No | Yes | No | No |
| *Melanoxylon brauna* | No | Yes | No | Yes |
| *Meliosma sellowii* | No | Yes | No | Yes |
| *Meriania calyptrata* | No | Yes | No | No |
| *Meriania claussenii* | No | Yes | No | No |
| *Meriania glabra* | No | Yes | No | No |
| *Meriania paniculata* | No | Yes | No | No |
| *Meriania tetramera* | No | Yes | No | No |
| *Mesocapparis lineata* | No | Yes | No | No |
| *Metrodorea maracasana* | No | Yes | No | No |
| *Metrodorea nigra* | No | Yes | No | No |
| *Metrodorea stipularis* | No | Yes | No | Yes |
| *Metternichia princeps* | No | Yes | No | Yes |

| | | | | |
|---|---|---|---|---|
| *Miconia albicans* | Yes | Yes | Yes | No |
| *Miconia amacurensis* | No | Yes | No | No |
| *Miconia amoena* | No | Yes | No | No |
| *Miconia argyrophylla* | No | Yes | No | No |
| *Miconia brasiliensis* | No | Yes | No | No |
| *Miconia brevipes* | No | Yes | No | No |
| *Miconia brunnea* | No | Yes | No | No |
| *Miconia budlejoides* | No | Yes | No | No |
| *Miconia cabucu* | No | Yes | No | Yes |
| *Miconia calvescens* | Yes | Yes | No | Yes |
| *Miconia centrodesma* | Yes | Yes | No | No |
| *Miconia chamissois* | No | Yes | No | No |
| *Miconia chartacea* | Yes | Yes | No | No |
| *Miconia ciliata* | No | Yes | Yes | No |
| *Miconia cinerascens* | No | Yes | No | No |
| *Miconia cinnamomifolia* | No | Yes | No | Yes |
| *Miconia collatata* | No | Yes | No | No |
| *Miconia compressa* | No | Yes | No | No |
| *Miconia corallina* | No | Yes | No | No |
| *Miconia cubatanensis* | No | Yes | Yes | No |
| *Miconia cuspidata* | No | Yes | No | No |
| *Miconia discolor* | No | Yes | No | No |
| *Miconia dodecandra* | No | Yes | No | No |
| *Miconia eichleri* | No | Yes | No | No |
| *Miconia elegans* | No | Yes | No | No |
| *Miconia fasciculata* | No | Yes | No | No |
| *Miconia ferruginata* | Yes | Yes | No | No |
| *Miconia formosa* | No | Yes | No | No |
| *Miconia holosericea* | Yes | Yes | Yes | Yes |
| *Miconia hyemalis* | No | Yes | No | Yes |
| *Miconia hypoleuca* | No | Yes | No | No |
| *Miconia ibaguensis* | No | Yes | No | Yes |
| *Miconia inconspicua* | No | Yes | No | No |
| *Miconia jucunda* | No | Yes | No | No |
| *Miconia latecrenata* | No | Yes | No | No |
| *Miconia lepidota* | Yes | Yes | No | Yes |
| *Miconia ligustroides* | Yes | Yes | No | Yes |
| *Miconia longicuspis* | No | Yes | No | No |

| | | | | |
|---|---|---|---|---|
| *Miconia macrothyrsa* | No | Yes | No | No |
| *Miconia minutiflora* | Yes | Yes | No | No |
| *Miconia mirabilis* | No | Yes | No | Yes |
| *Miconia nervosa* | Yes | Yes | No | No |
| *Miconia octopetala* | No | Yes | No | No |
| *Miconia paniculata* | No | Yes | No | No |
| *Miconia paucidens* | No | Yes | No | No |
| *Miconia pepericarpa* | No | Yes | No | No |
| *Miconia petropolitana* | No | Yes | No | No |
| *Miconia polyandra* | No | Yes | No | No |
| *Miconia prasina* | Yes | Yes | Yes | Yes |
| *Miconia pusilliflora* | No | Yes | No | No |
| *Miconia pyrifolia* | Yes | Yes | No | Yes |
| *Miconia rimalis* | No | Yes | No | No |
| *Miconia rubiginosa* | Yes | Yes | Yes | No |
| *Miconia ruficalyx* | No | Yes | No | No |
| *Miconia saldanhae* | No | Yes | No | No |
| *Miconia sclerophylla* | No | Yes | No | No |
| *Miconia sellowiana* | No | Yes | No | No |
| *Miconia serialis* | No | Yes | No | No |
| *Miconia serrulata* | No | Yes | No | No |
| *Miconia splendens* | No | Yes | No | Yes |
| *Miconia stenostachya* | No | Yes | No | No |
| *Miconia theizans* | No | Yes | No | No |
| *Miconia tomentosa* | Yes | Yes | Yes | Yes |
| *Miconia trianae* | No | Yes | No | No |
| *Miconia tristis* | No | Yes | No | No |
| *Miconia urophylla* | No | Yes | No | No |
| *Miconia valtheri* | No | Yes | No | Yes |
| *Micrandra elata* | Yes | Yes | No | Yes |
| *Micropholis crassipedicellata* | No | Yes | No | Yes |
| *Micropholis gardneriana* | No | Yes | No | No |
| *Micropholis gnaphaloclados* | No | Yes | No | No |
| *Micropholis guyanensis* | Yes | Yes | No | Yes |
| *Micropholis venulosa* | Yes | Yes | No | Yes |
| *Mimosa adenophylla* | No | Yes | No | No |
| *Mimosa arenosa* | Yes | Yes | Yes | Yes |
| *Mimosa artemisiana* | No | Yes | No | Yes |

| | | | | |
|---|---|---|---|---|
| *Mimosa bimucronata* | No | Yes | No | Yes |
| *Mimosa caesalpiniifolia* | No | Yes | No | No |
| *Mimosa furfuracea* | No | Yes | No | No |
| *Mimosa gemmulata* | No | Yes | No | No |
| *Mimosa pigra* | No | Yes | Yes | Yes |
| *Mimosa scabrella* | No | Yes | Yes | Yes |
| *Mimosa tenuiflora* | Yes | Yes | Yes | Yes |
| *Moldenhawera blanchetiana* | No | Yes | No | No |
| *Moldenhawera emarginata* | No | Yes | No | No |
| *Moldenhawera floribunda* | No | Yes | No | Yes |
| *Mollinedia argyrogyna* | No | Yes | No | No |
| *Mollinedia blumenaviana* | No | Yes | No | No |
| *Mollinedia boracensis* | No | Yes | No | No |
| *Mollinedia calodonta* | No | Yes | No | No |
| *Mollinedia clavigera* | No | Yes | No | No |
| *Mollinedia elegans* | No | Yes | No | No |
| *Mollinedia engleriana* | No | Yes | No | No |
| *Mollinedia fruticulosa* | No | Yes | No | No |
| *Mollinedia gilgiana* | No | Yes | No | No |
| *Mollinedia glabra* | No | Yes | No | No |
| *Mollinedia lanceolata* | No | Yes | No | No |
| *Mollinedia longifolia* | No | Yes | No | No |
| *Mollinedia oligantha* | No | Yes | No | No |
| *Mollinedia ovata* | No | Yes | No | No |
| *Mollinedia schottiana* | Yes | Yes | No | Yes |
| *Mollinedia triflora* | No | Yes | No | No |
| *Mollinedia uleana* | No | Yes | Yes | No |
| *Mollinedia widgrenii* | No | Yes | No | Yes |
| *Molopanthera paniculata* | No | Yes | No | No |
| *Monilicarpa brasiliana* | No | Yes | No | No |
| *Mouriri arborea* | No | Yes | No | No |
| *Mouriri chamissoana* | No | Yes | Yes | No |
| *Mouriri glazioviana* | No | Yes | No | Yes |
| *Myracrodruon urundeuva* | No | Yes | No | Yes |
| *Myrceugenia acutiflora* | No | Yes | No | No |
| *Myrceugenia alpigena* | No | Yes | No | No |
| *Myrceugenia bracteosa* | No | Yes | No | No |
| *Myrceugenia campestris* | No | Yes | No | No |

| | | | | |
|---|---|---|---|---|
| *Myrceugenia cucullata* | No | Yes | No | No |
| *Myrceugenia euosma* | No | Yes | No | Yes |
| *Myrceugenia glaucescens* | No | Yes | No | No |
| *Myrceugenia kleinii* | No | Yes | No | No |
| *Myrceugenia mesomischa* | No | Yes | No | No |
| *Myrceugenia miersiana* | Yes | Yes | No | Yes |
| *Myrceugenia myrcioides* | No | Yes | No | Yes |
| *Myrceugenia myrtoides* | No | Yes | No | No |
| *Myrceugenia ovalifolia* | No | Yes | No | No |
| *Myrceugenia ovata* | Yes | Yes | No | No |
| *Myrceugenia oxysepala* | No | Yes | No | No |
| *Myrceugenia pilotantha* | No | Yes | No | No |
| *Myrceugenia reitzii* | No | Yes | No | No |
| *Myrceugenia rufescens* | No | Yes | No | No |
| *Myrcia aethusa* | No | Yes | No | No |
| *Myrcia amazonica* | No | Yes | No | Yes |
| *Myrcia anacardiifolia* | No | Yes | Yes | No |
| *Myrcia anceps* | No | Yes | No | No |
| *Myrcia bergiana* | No | Yes | No | No |
| *Myrcia bicolor* | No | Yes | No | No |
| *Myrcia brasiliensis* | Yes | Yes | No | No |
| *Myrcia cerqueiria* | No | Yes | No | No |
| *Myrcia citrifolia* | No | No | Yes | No |
| *Myrcia cordiifolia* | No | Yes | No | No |
| *Myrcia crocea* | No | Yes | No | No |
| *Myrcia dichrophylla* | No | Yes | No | No |
| *Myrcia eriocalyx* | No | Yes | No | No |
| *Myrcia eriopus* | No | Yes | No | No |
| *Myrcia eximia* | No | Yes | No | No |
| *Myrcia fenzliana* | No | Yes | No | No |
| *Myrcia flagellaris* | No | Yes | No | No |
| *Myrcia glabra* | No | Yes | No | No |
| *Myrcia grandifolia* | No | Yes | No | No |
| *Myrcia guianensis* | Yes | Yes | No | Yes |
| *Myrcia hartwegiana* | No | Yes | No | No |
| *Myrcia hatschbachii* | No | Yes | No | Yes |
| *Myrcia hebepetala* | No | Yes | No | No |
| *Myrcia heringii* | No | No | No | No |

| | | | | |
|---|---|---|---|---|
| *Myrcia ilheosensis* | No | Yes | No | No |
| *Myrcia insularis* | No | Yes | No | No |
| *Myrcia lajeana* | No | Yes | No | No |
| *Myrcia laruotteana* | No | Yes | No | No |
| *Myrcia laxiflora* | No | Yes | No | No |
| *Myrcia mischophylla* | No | Yes | No | No |
| *Myrcia montana* | No | Yes | No | No |
| *Myrcia multiflora* | No | Yes | No | No |
| *Myrcia mutabilis* | No | Yes | No | No |
| *Myrcia neoblanchetiana* | No | Yes | No | No |
| *Myrcia neobullata* | No | Yes | No | No |
| *Myrcia oblongata* | No | Yes | No | No |
| *Myrcia obovata* | No | Yes | No | No |
| *Myrcia oligantha* | No | Yes | No | No |
| *Myrcia ovata* | No | Yes | No | No |
| *Myrcia palustris* | No | Yes | No | Yes |
| *Myrcia pubescens* | No | Yes | No | No |
| *Myrcia pubiflora* | No | Yes | No | No |
| *Myrcia pubipetala* | No | Yes | Yes | Yes |
| *Myrcia pulchra* | No | Yes | No | No |
| *Myrcia racemosa* | Yes | Yes | No | No |
| *Myrcia retorta* | No | Yes | No | No |
| *Myrcia rufipes* | No | Yes | No | No |
| *Myrcia rupicola* | No | Yes | No | No |
| *Myrcia selloi* | No | Yes | No | No |
| *Myrcia silvatica* | No | Yes | No | No |
| *Myrcia spectabilis* | No | Yes | No | Yes |
| *Myrcia splendens* | Yes | Yes | No | Yes |
| *Myrcia tenuivenosa* | No | Yes | No | No |
| *Myrcia thyrsoidea* | No | Yes | No | No |
| *Myrcia tijucensis* | No | Yes | No | No |
| *Myrcia tomentosa* | Yes | Yes | Yes | Yes |
| *Myrcia undulata* | No | Yes | No | No |
| *Myrcia variabilis* | No | Yes | No | No |
| *Myrcia venulosa* | No | Yes | No | No |
| *Myrcia vittoriana* | No | Yes | No | No |
| *Myrcianthes gigantea* | Yes | Yes | No | Yes |
| *Myrcianthes pungens* | Yes | Yes | Yes | Yes |

| | | | | |
|---|---|---|---|---|
| *Myrciaria cuspidata* | No | Yes | No | Yes |
| *Myrciaria delicatula* | No | Yes | No | Yes |
| *Myrciaria disticha* | No | Yes | No | No |
| *Myrciaria ferruginea* | No | Yes | No | No |
| *Myrciaria floribunda* | Yes | Yes | No | Yes |
| *Myrciaria glanduliflora* | No | Yes | No | No |
| *Myrciaria glazioviana* | No | Yes | No | No |
| *Myrciaria glomerata* | No | Yes | No | No |
| *Myrciaria plinioides* | No | Yes | No | Yes |
| *Myrciaria strigipes* | No | Yes | No | No |
| *Myrciaria tenella* | No | Yes | No | No |
| *Myrocarpus fastigiatus* | No | Yes | No | Yes |
| *Myrocarpus frondosus* | Yes | Yes | No | Yes |
| *Myroxylon peruiferum* | No | Yes | Yes | Yes |
| *Myrrhinium atropurpureum* | Yes | Yes | No | Yes |
| *Myrsine altomontana* | No | Yes | No | No |
| *Myrsine balansae* | No | Yes | No | No |
| *Myrsine coriacea* | Yes | Yes | Yes | Yes |
| *Myrsine gardneriana* | No | Yes | No | No |
| *Myrsine guianensis* | No | Yes | No | No |
| *Myrsine hermogenesii* | No | Yes | No | No |
| *Myrsine laetevirens* | Yes | Yes | Yes | Yes |
| *Myrsine lancifolia* | No | Yes | No | No |
| *Myrsine lineata* | No | Yes | No | No |
| *Myrsine loefgrenii* | No | Yes | No | No |
| *Myrsine matensis* | No | Yes | No | No |
| *Myrsine monticola* | No | Yes | No | No |
| *Myrsine parvifolia* | No | Yes | No | No |
| *Myrsine parvula* | No | Yes | No | No |
| *Myrsine umbellata* | Yes | Yes | Yes | Yes |
| *Myrsine venosa* | No | Yes | No | No |
| *Myrsine villosisYesa* | No | Yes | No | No |
| *Naucleopsis oblongifolia* | No | Yes | No | No |
| *Nectandra angustifolia* | No | Yes | No | Yes |
| *Nectandra barbellata* | No | Yes | No | No |
| *Nectandra cissiflora* | Yes | Yes | Yes | Yes |
| *Nectandra cuspidata* | Yes | Yes | No | Yes |
| *Nectandra globosa* | No | Yes | No | Yes |

| | | | | |
|---|---|---|---|---|
| *Nectandra grandiflora* | No | Yes | No | Yes |
| *Nectandra lanceolata* | No | Yes | No | Yes |
| *Nectandra leucantha* | No | Yes | No | No |
| *Nectandra megapotamica* | Yes | Yes | Yes | Yes |
| *Nectandra membranacea* | Yes | Yes | No | Yes |
| *Nectandra nitidula* | No | Yes | No | Yes |
| *Nectandra oppositifolia* | Yes | Yes | No | Yes |
| *Nectandra psammophila* | No | Yes | No | No |
| *Nectandra puberula* | No | Yes | No | Yes |
| *Nectandra reticulata* | No | Yes | Yes | Yes |
| *Neea floribunda* | Yes | Yes | No | Yes |
| *Neea macrophylla* | Yes | Yes | No | No |
| *Neea parviflora* | No | Yes | No | No |
| *Neea schwackeana* | No | Yes | No | No |
| *Neocalyptrocalyx longifolium* | No | Yes | No | No |
| *Neocalyptrocalyx nectareus* | No | Yes | No | No |
| *Neomitranthes cordifolia* | No | Yes | No | No |
| *Neomitranthes glomerata* | No | Yes | Yes | No |
| *Neomitranthes obscura* | No | Yes | No | Yes |
| *Neomitranthes obtusa* | No | Yes | No | No |
| *Neomitranthes warmingiana* | No | Yes | No | No |
| *Neoptychocarpus apodanthus* | No | Yes | No | No |
| *Neoraputia alba* | No | Yes | No | Yes |
| *Neoraputia magnifica* | No | Yes | No | No |
| *Neoraputia trifoliata* | No | Yes | No | No |
| *Noticastrum calvatum* | No | Yes | No | No |
| *Noticastrum decumbens* | No | Yes | No | No |
| *Notopleura tapajozensis* | No | Yes | No | No |
| *Ocotea aciphylla* | Yes | Yes | No | Yes |
| *Ocotea acutifolia* | No | Yes | No | Yes |
| *Ocotea bicolor* | No | Yes | No | No |
| *Ocotea brachybotra* | No | Yes | No | No |
| *Ocotea canaliculata* | No | Yes | No | Yes |
| *Ocotea catharinensis* | No | Yes | No | Yes |
| *Ocotea cernua* | Yes | Yes | Yes | Yes |
| *Ocotea complicata* | No | No | No | No |
| *Ocotea corymbosa* | No | Yes | No | No |
| *Ocotea daphnifolia* | No | Yes | No | No |

| | | | | |
|---|---|---|---|---|
| *Ocotea deflexa* | No | Yes | No | No |
| *Ocotea densiflora* | No | Yes | No | No |
| *Ocotea diospyrifolia* | No | Yes | No | Yes |
| *Ocotea dispersa* | Yes | Yes | Yes | Yes |
| *Ocotea divaricata* | No | Yes | No | No |
| *Ocotea duckei* | No | Yes | No | No |
| *Ocotea elegans* | No | Yes | No | Yes |
| *Ocotea floribunda* | Yes | Yes | No | Yes |
| *Ocotea gardneri* | No | Yes | No | No |
| *Ocotea glauca* | No | Yes | No | No |
| *Ocotea glaziovii* | No | Yes | No | No |
| *Ocotea glomerata* | No | Yes | No | Yes |
| *Ocotea indecora* | Yes | Yes | No | No |
| *Ocotea insignis* | No | Yes | No | No |
| *Ocotea lanata* | No | Yes | No | No |
| *Ocotea lancifolia* | No | Yes | No | No |
| *Ocotea laxa* | No | Yes | No | No |
| *Ocotea leucoxylon* | Yes | Yes | No | Yes |
| *Ocotea lobbii* | No | Yes | No | No |
| *Ocotea longifolia* | Yes | Yes | No | No |
| *Ocotea magnilimba* | No | Yes | No | No |
| *Ocotea mandioccana* | No | Yes | No | Yes |
| *Ocotea minarum* | No | Yes | No | Yes |
| *Ocotea montana* | No | Yes | No | No |
| *Ocotea nectandrifolia* | No | Yes | No | No |
| *Ocotea neesiana* | No | Yes | No | Yes |
| *Ocotea nitida* | No | Yes | No | No |
| *Ocotea notata* | No | Yes | No | No |
| *Ocotea nunesiana* | No | Yes | No | No |
| *Ocotea nutans* | No | Yes | No | No |
| *Ocotea odorifera* | No | Yes | No | Yes |
| *Ocotea oppositifolia* | No | Yes | No | No |
| *Ocotea paranapiacabensis* | No | Yes | No | No |
| *Ocotea pauciflora* | No | Yes | No | Yes |
| *Ocotea percurrens* | Yes | Yes | No | Yes |
| *Ocotea pomaderroides* | No | Yes | No | No |
| *Ocotea porosa* | No | Yes | No | Yes |
| *Ocotea puberula* | Yes | Yes | Yes | Yes |

| | | | | |
|---|---|---|---|---|
| *Ocotea pulchella* | Yes | Yes | No | Yes |
| *Ocotea pulchra* | No | Yes | No | No |
| *Ocotea silvestris* | No | Yes | No | No |
| *Ocotea spectabilis* | No | Yes | No | No |
| *Ocotea spixiana* | No | Yes | No | Yes |
| *Ocotea tabacifolia* | No | Yes | No | No |
| *Ocotea teleiandra* | No | Yes | Yes | No |
| *Ocotea tenuiflora* | No | Yes | No | No |
| *Ocotea tristis* | No | Yes | No | No |
| *Ocotea vaccinioides* | No | Yes | No | No |
| *Ocotea velloziana* | No | Yes | No | No |
| *Ocotea velutina* | No | Yes | No | No |
| *Ocotea venulosa* | No | Yes | No | No |
| *Ocotea villosa* | No | Yes | No | No |
| *Opuntia ficus-indica* | No | Yes | Yes | No |
| *Oreopanax capitatus* | No | Yes | No | No |
| *Oreopanax fulvus* | No | Yes | No | Yes |
| *Ormosia arborea* | No | Yes | No | Yes |
| *Ormosia bahiensis* | No | Yes | No | No |
| *Ormosia fastigiata* | No | Yes | No | No |
| *Ossaea amygdaloides* | No | Yes | No | No |
| *Ossaea angustifolia* | No | Yes | No | No |
| *Ossaea confertiflora* | No | Yes | No | No |
| *Ossaea marginata* | No | Yes | No | No |
| *Ossaea sanguinea* | No | Yes | No | No |
| *Ouratea castaneifolia* | No | Yes | No | Yes |
| *Ouratea crassa* | No | Yes | No | No |
| *Ouratea cuspidata* | No | Yes | No | No |
| *Ouratea fieldingiana* | No | Yes | No | No |
| *Ouratea hexasperma* | Yes | Yes | No | No |
| *Ouratea multiflora* | No | Yes | No | No |
| *Ouratea parviflora* | Yes | Yes | No | Yes |
| *Ouratea salicifolia* | No | Yes | No | No |
| *Ouratea sellowii* | No | Yes | No | No |
| *Ouratea semiserrata* | No | Yes | No | No |
| *Ouratea vaccinioides* | No | Yes | No | No |
| *Oxandra nitida* | No | Yes | No | No |
| *Pachira aquatica* | Yes | Yes | Yes | Yes |

| | | | | |
|---|---|---|---|---|
| *Pachira stenopetala* | No | Yes | No | No |
| *Pachystroma longifolium* | No | Yes | No | No |
| *Palicourea australis* | No | Yes | No | No |
| *Palicourea blanchetiana* | No | Yes | No | No |
| *Palicourea crocea* | No | Yes | No | Yes |
| *Palicourea croceoides* | No | Yes | No | No |
| *Palicourea guianensis* | Yes | Yes | Yes | Yes |
| *Palicourea macrobotrys* | No | Yes | No | No |
| *Palicourea marcgravii* | No | Yes | No | No |
| *Palicourea rigida* | No | Yes | No | No |
| *Palicourea tetraphylla* | No | Yes | No | No |
| *Panopsis rubescens* | Yes | Yes | No | Yes |
| *Paradrypetes ilicifolia* | No | Yes | No | No |
| *Paralychnophora santosii* | No | Yes | No | No |
| *Parapiptadenia pterosperma* | No | Yes | No | Yes |
| *Parapiptadenia rigida* | No | Yes | No | Yes |
| *Paratecoma peroba* | No | Yes | No | Yes |
| *Parinari excelsa* | Yes | Yes | Yes | Yes |
| *Parinari littoralis* | No | Yes | No | No |
| *Parinari obtusifolia* | No | Yes | No | No |
| *Parkia pendula* | No | Yes | Yes | Yes |
| *Pausandra morisiana* | Yes | Yes | No | Yes |
| *Pausandra trianae* | Yes | Yes | No | Yes |
| *Paypayrola blanchetiana* | No | Yes | No | No |
| *Paypayrola grandiflora* | No | Yes | No | Yes |
| *Peltastes peltatus* | No | Yes | No | No |
| *Peltogyne confertiflora* | No | Yes | No | Yes |
| *Peltogyne pauciflora* | No | Yes | No | No |
| *Peltogyne venosa* | Yes | Yes | No | Yes |
| *Peltophorum dubium* | No | Yes | No | Yes |
| *Pera anisotricha* | No | Yes | No | No |
| *Pera glabrata* | Yes | Yes | No | Yes |
| *Pera heteranthera* | No | Yes | No | Yes |
| *Pereskia grandifolia* | No | Yes | No | No |
| *Persea alba* | No | Yes | No | No |
| *Persea aurata* | No | Yes | No | No |
| *Persea caesia* | No | Yes | No | No |
| *Persea fulva* | No | Yes | No | No |

| | | | | |
|---|---|---|---|---|
| *Persea major* | No | Yes | No | Yes |
| *Persea rufotomentosa* | No | Yes | No | No |
| *Persea splendens* | No | Yes | No | No |
| *Persea venosa* | No | Yes | No | Yes |
| *Persea willdenovii* | No | Yes | No | Yes |
| *Phyllanthus acuminatus* | No | Yes | No | No |
| *Phyllanthus juglandifolius* | No | Yes | No | No |
| *Phyllostemonodaphne geminiflora* | No | Yes | No | No |
| *Phytolacca dioica* | No | Yes | Yes | Yes |
| *Picramnia andrade-limae* | No | Yes | No | No |
| *Picramnia bahiensis* | No | Yes | No | No |
| *Picramnia ciliata* | No | Yes | No | No |
| *Picramnia excelsa* | No | Yes | No | No |
| *Picramnia gardneri* | No | Yes | No | No |
| *Picramnia glazioviana* | No | Yes | No | No |
| *Picramnia parvifolia* | No | Yes | No | No |
| *Picramnia ramiflora* | No | Yes | No | No |
| *Picramnia sellowii* | Yes | Yes | No | Yes |
| *Picrasma crenata* | No | Yes | No | Yes |
| *Pilocarpus microphyllus* | No | Yes | No | No |
| *Pilocarpus pauciflorus* | No | Yes | No | No |
| *Pilocarpus pennatifolius* | Yes | Yes | No | Yes |
| *Pilocarpus riedelianus* | No | Yes | No | No |
| *Pilocarpus spicatus* | No | Yes | No | No |
| *Pilosocereus arrabidae* | No | Yes | No | No |
| *Pimenta pseudocaryophyllus* | No | Yes | No | Yes |
| *Piper aduncum* | Yes | Yes | No | Yes |
| *Piper amalago* | Yes | Yes | Yes | No |
| *Piper amplum* | No | Yes | No | No |
| *Piper anonifolium* | No | Yes | No | No |
| *Piper arboreum* | Yes | Yes | Yes | No |
| *Piper bowiei* | No | Yes | No | No |
| *Piper caldense* | No | Yes | No | No |
| *Piper cernuum* | No | Yes | No | No |
| *Piper corcovadensis* | No | Yes | No | No |
| *Piper crassinervium* | Yes | Yes | Yes | No |
| *Piper cuyabanum* | No | Yes | No | No |
| *Piper dilatatum* | No | Yes | Yes | No |

| | | | | |
|---|---|---|---|---|
| *Piper divaricatum* | No | Yes | No | No |
| *Piper eucalyptophyllum* | No | Yes | No | No |
| *Piper gaudichaudianum* | No | Yes | No | No |
| *Piper glabratum* | No | Yes | No | No |
| *Piper hayneanum* | No | Yes | No | No |
| *Piper hispidum* | Yes | Yes | Yes | No |
| *Piper ilheusense* | No | Yes | No | No |
| *Piper lhotzkyanum* | No | Yes | No | No |
| *Piper malacophyllum* | No | Yes | No | No |
| *Piper marginatum* | No | Yes | Yes | No |
| *Piper miquelianum* | No | Yes | No | No |
| *Piper mollicomum* | No | Yes | No | No |
| *Piper mosenii* | No | Yes | No | No |
| *Piper obliquum* | No | Yes | No | No |
| *Piper ovatum* | No | Yes | No | No |
| *Piper richardiifolium* | No | Yes | No | No |
| *Piper tuberculatum* | No | Yes | Yes | No |
| *Piper umbellatum* | Yes | Yes | Yes | No |
| *Piper vicosanum* | No | Yes | No | No |
| *Piper viminifolium* | No | Yes | No | No |
| *Piper xylosteoides* | No | Yes | No | No |
| *Piptadenia gonoacantha* | No | Yes | No | Yes |
| *Piptadenia paniculata* | Yes | Yes | No | Yes |
| *Piptadenia stipulacea* | Yes | Yes | No | Yes |
| *Piptadenia trisperma* | No | Yes | No | No |
| *Piptadenia viridiflora* | No | Yes | No | Yes |
| *Piptocarpha angustifolia* | No | Yes | No | Yes |
| *Piptocarpha axillaris* | No | Yes | No | Yes |
| *Piptocarpha densifolia* | No | Yes | No | No |
| *Piptocarpha macropoda* | No | Yes | No | No |
| *Piptocarpha oblonga* | No | Yes | No | No |
| *Piptocarpha organensis* | No | Yes | No | No |
| *Piptocarpha regnellii* | No | Yes | No | No |
| *Piptocarpha rotundifolia* | Yes | Yes | No | Yes |
| *Piptocarpha sellowii* | No | Yes | No | No |
| *Pisonia ambigua* | No | Yes | No | Yes |
| *Pityrocarpa moniliformis* | No | Yes | No | Yes |
| *Plathymenia reticulata* | No | Yes | No | Yes |

| | | | | |
|---|---|---|---|---|
| *Platycyamus regnellii* | No | Yes | No | Yes |
| *Platymiscium floribundum* | No | Yes | No | Yes |
| *Platymiscium pubescens* | No | Yes | No | No |
| *Platypodium elegans* | Yes | Yes | Yes | Yes |
| *Plenckia populnea* | No | Yes | No | No |
| *Plinia callosa* | No | Yes | No | No |
| *Plinia cauliflora* | No | Yes | No | No |
| *Plinia complanata* | No | Yes | Yes | No |
| *Plinia cordifolia* | No | Yes | No | No |
| *Plinia edulis* | No | Yes | No | Yes |
| *Plinia peruviana* | No | Yes | No | No |
| *Plinia pseudodichasiantha* | No | Yes | No | No |
| *Plinia rivularis* | Yes | Yes | No | Yes |
| *Pluchea sagittalis* | No | Yes | No | No |
| *Podocarpus lambertii* | Yes | Yes | No | Yes |
| *Podocarpus sellowii* | No | Yes | No | Yes |
| *Poecilanthe falcata* | No | Yes | No | No |
| *Poecilanthe parviflora* | No | Yes | No | Yes |
| *Poecilanthe ulei* | No | Yes | No | Yes |
| *Poeppigia procera* | Yes | Yes | No | Yes |
| *Pogonophora schomburgkiana* | Yes | Yes | No | Yes |
| *Poincianella gardneriana* | No | Yes | No | No |
| *Poincianella pluviosa* | No | Yes | No | No |
| *Poincianella pyramidalis* | No | Yes | No | No |
| *Porcelia macrocarpa* | No | Yes | No | Yes |
| *Posoqueria acutifolia* | No | Yes | No | Yes |
| *Posoqueria latifolia* | Yes | Yes | Yes | Yes |
| *Posoqueria longiflora* | Yes | Yes | No | Yes |
| *Pourouma bicolor* | Yes | Yes | Yes | Yes |
| *Pourouma cecropiifolia* | Yes | Yes | No | Yes |
| *Pourouma guianensis* | Yes | Yes | Yes | Yes |
| *Pourouma mollis* | Yes | Yes | No | Yes |
| *Pourouma velutina* | No | Yes | No | No |
| *Pouteria bangii* | Yes | Yes | No | Yes |
| *Pouteria beaurepairei* | No | Yes | No | No |
| *Pouteria bilocularis* | Yes | Yes | No | Yes |
| *Pouteria bullata* | No | Yes | No | Yes |
| *Pouteria caimito* | Yes | Yes | No | Yes |

| | | | | |
|---|---|---|---|---|
| *Pouteria cuspidata* | Yes | Yes | Yes | Yes |
| *Pouteria durlandii* | Yes | Yes | Yes | Yes |
| *Pouteria filipes* | Yes | Yes | No | Yes |
| *Pouteria gardneri* | No | Yes | No | Yes |
| *Pouteria gardneriana* | No | Yes | No | Yes |
| *Pouteria glomerata* | Yes | Yes | Yes | Yes |
| *Pouteria grandiflora* | No | Yes | No | Yes |
| *Pouteria guianensis* | Yes | Yes | Yes | Yes |
| *Pouteria macahensis* | No | Yes | No | No |
| *Pouteria macrophylla* | Yes | Yes | Yes | Yes |
| *Pouteria procera* | Yes | Yes | No | No |
| *Pouteria psammophila* | No | Yes | No | No |
| *Pouteria ramiflora* | No | Yes | No | No |
| *Pouteria reticulata* | Yes | Yes | Yes | Yes |
| *Pouteria torta* | Yes | Yes | No | Yes |
| *Pouteria venosa* | Yes | Yes | No | Yes |
| *Pradosia lactescens* | No | Yes | No | No |
| *Prockia crucis* | No | Yes | No | Yes |
| *Protium aracouchini* | Yes | Yes | No | Yes |
| *Protium bahianum* | No | Yes | No | No |
| *Protium brasiliense* | Yes | Yes | No | Yes |
| *Protium heptaphyllum* | Yes | Yes | No | Yes |
| *Protium icicariba* | No | Yes | No | No |
| *Protium kleinii* | No | Yes | No | Yes |
| *Protium ovatum* | No | Yes | No | No |
| *Protium sagotianum* | Yes | Yes | No | Yes |
| *Protium spruceanum* | Yes | Yes | No | Yes |
| *Protium warmingianum* | No | Yes | No | No |
| *Protium widgrenii* | No | Yes | Yes | No |
| *Prunus brasiliensis* | No | Yes | No | Yes |
| *Prunus myrtifolia* | Yes | Yes | No | Yes |
| *Prunus subcoriacea* | No | Yes | No | No |
| *Pseudobombax grandiflorum* | No | Yes | No | Yes |
| *Pseudobombax longiflorum* | No | Yes | No | No |
| *Pseudobombax marginatum* | No | Yes | No | Yes |
| *Pseudobombax tomentosum* | No | Yes | No | No |
| *Pseudolmedia macrophylla* | Yes | Yes | No | Yes |
| *Pseudopiptadenia bahiana* | No | Yes | No | No |

| | | | | |
|---|---|---|---|---|
| *Pseudopiptadenia contorta* | No | Yes | No | Yes |
| *Pseudopiptadenia inaequalis* | No | Yes | No | No |
| *Pseudopiptadenia leptostachya* | No | Yes | No | No |
| *Pseudopiptadenia psilostachya* | Yes | Yes | No | Yes |
| *Pseudopiptadenia warmingii* | No | Yes | No | Yes |
| *Pseudoxandra bahiensis* | No | Yes | No | No |
| *Pseudoxandra spiritus-sancti* | No | Yes | No | No |
| *Psidium acutangulum* | No | Yes | Yes | Yes |
| *Psidium appendiculatum* | No | Yes | No | No |
| *Psidium brownianum* | No | Yes | No | No |
| *Psidium cattleianum* | Yes | Yes | No | Yes |
| *Psidium decussatum* | No | Yes | No | No |
| *Psidium firmum* | No | Yes | No | No |
| *Psidium grandifolium* | No | Yes | No | No |
| *Psidium longipetiolatum* | No | Yes | No | Yes |
| *Psidium myrsinites* | No | Yes | No | No |
| *Psidium myrtoides* | No | Yes | No | No |
| *Psidium nutans* | No | Yes | No | No |
| *Psidium oligospermum* | No | Yes | No | No |
| *Psidium ovale* | No | Yes | No | No |
| *Psidium rufum* | No | Yes | No | Yes |
| *Psidium sartorianum* | No | Yes | No | Yes |
| *Psychotria alba* | No | Yes | No | No |
| *Psychotria appendiculata* | No | Yes | No | No |
| *Psychotria bahiensis* | No | Yes | No | No |
| *Psychotria brachyceras* | No | Yes | No | No |
| *Psychotria brachypoda* | No | Yes | Yes | No |
| *Psychotria bracteocardia* | No | Yes | No | No |
| *Psychotria capitata* | No | Yes | No | No |
| *Psychotria carthagenensis* | Yes | Yes | Yes | Yes |
| *Psychotria colorata* | No | Yes | No | No |
| *Psychotria deflexa* | Yes | Yes | Yes | No |
| *Psychotria forsteronioides* | No | Yes | No | No |
| *Psychotria fractistipula* | No | Yes | No | No |
| *Psychotria glaziovii* | No | Yes | No | No |
| *Psychotria gracilenta* | No | Yes | Yes | No |
| *Psychotria hastisepala* | No | Yes | No | No |
| *Psychotria hoffmannseggiana* | No | Yes | Yes | No |

| | | | | |
|---|---|---|---|---|
| *Psychotria iodotricha* | No | Yes | No | No |
| *Psychotria jambosioides* | No | Yes | No | No |
| *Psychotria laciniata* | No | Yes | No | No |
| *Psychotria leiocarpa* | No | Yes | No | No |
| *Psychotria leitana* | No | Yes | No | No |
| *Psychotria mapourioides* | No | Yes | Yes | No |
| *Psychotria myriantha* | No | Yes | No | No |
| *Psychotria nemorosa* | No | Yes | No | No |
| *Psychotria nuda* | Yes | Yes | No | Yes |
| *Psychotria officinalis* | Yes | Yes | No | No |
| *Psychotria patentinervia* | No | Yes | No | No |
| *Psychotria platypoda* | No | Yes | No | No |
| *Psychotria pleiocephala* | No | Yes | No | No |
| *Psychotria purpurascens* | No | Yes | No | No |
| *Psychotria racemosa* | Yes | Yes | Yes | No |
| *Psychotria ruelliifolia* | No | Yes | No | No |
| *Psychotria schlechtendaliana* | No | Yes | No | No |
| *Psychotria stachyoides* | No | Yes | No | No |
| *Psychotria subtriflora* | No | Yes | No | No |
| *Psychotria suterella* | No | Yes | Yes | Yes |
| *Psychotria tenerior* | No | Yes | No | No |
| *Psychotria tenuifolia* | Yes | Yes | Yes | No |
| *Psychotria vellosiana* | No | Yes | No | Yes |
| *Pterocarpus rohrii* | Yes | Yes | Yes | Yes |
| *Pterodon emarginatus* | Yes | Yes | No | Yes |
| *Pterogyne nitens* | No | Yes | No | Yes |
| *Qualea cordata* | No | Yes | No | No |
| *Qualea cryptantha* | No | Yes | No | No |
| *Qualea dichotoma* | Yes | Yes | No | No |
| *Qualea multiflora* | No | Yes | No | Yes |
| *Qualea selloi* | No | Yes | No | No |
| *Quararibea penduliflora* | No | Yes | No | No |
| *Quararibea turbinata* | Yes | Yes | No | Yes |
| *Quiina glaziovii* | No | Yes | Yes | No |
| *Quiina paraensis* | No | Yes | No | No |
| *Quillaja brasiliensis* | No | Yes | No | Yes |
| *Ramisia brasiliensis* | No | Yes | No | Yes |
| *Randia armata* | Yes | Yes | Yes | Yes |

| | | | | |
|---|---|---|---|---|
| *Randia ferox* | No | Yes | No | Yes |
| *Rauia nodosa* | No | Yes | No | No |
| *Rauia resinosa* | No | Yes | No | No |
| *Raulinoreitzia leptophlebia* | No | Yes | No | No |
| *Rauvolfia bahiensis* | No | Yes | No | No |
| *Rauvolfia grandiflora* | No | Yes | No | No |
| *Rauvolfia ligustrina* | No | Yes | No | No |
| *Rauvolfia sellowii* | No | Yes | No | Yes |
| *Ravenia infelix* | No | Yes | No | No |
| *Remijia ferruginea* | No | Yes | No | No |
| *Rhamnidium elaeocarpum* | Yes | Yes | No | Yes |
| *Rhamnus sphaerosperma* | No | Yes | No | Yes |
| *Rhodostemonodaphne capixabensis* | No | Yes | No | No |
| *Rhodostemonodaphne macrocalyx* | No | Yes | No | No |
| *Richeria grandis* | Yes | Yes | No | Yes |
| *Rinorea bahiensis* | No | Yes | No | Yes |
| *Rinorea guianensis* | Yes | Yes | No | Yes |
| *Ronabea latifolia* | No | Yes | No | No |
| *Roucheria columbiana* | Yes | Yes | No | Yes |
| *Roupala conYesilis* | No | Yes | No | No |
| *Roupala montana* | Yes | Yes | No | Yes |
| *Roupala paulensis* | No | Yes | No | No |
| *Roupala rhombifolia* | No | Yes | No | No |
| *Rourea doniana* | No | Yes | No | No |
| *Rourea induta* | No | Yes | No | No |
| *Rourea martiana* | No | Yes | No | No |
| *Rubus brasiliensis* | No | Yes | No | No |
| *Rubus erythrocladus* | No | Yes | No | No |
| *Rubus sellowii* | No | Yes | No | No |
| *Rudgea coriacea* | No | Yes | No | No |
| *Rudgea coronata* | No | Yes | No | No |
| *Rudgea crassifolia* | No | Yes | No | No |
| *Rudgea gardenioides* | No | Yes | No | No |
| *Rudgea interrupta* | No | Yes | No | No |
| *Rudgea jacobinensis* | No | Yes | No | No |
| *Rudgea jasminoides* | No | Yes | Yes | No |
| *Rudgea parquioides* | No | Yes | No | No |
| *Rudgea recurva* | No | Yes | No | No |

| | | | | |
|---|---|---|---|---|
| *Rudgea reticulata* | No | Yes | No | No |
| *Rudgea vellerea* | No | Yes | No | No |
| *Rudgea viburnoides* | No | Yes | No | Yes |
| *Ruprechtia laxiflora* | Yes | Yes | Yes | Yes |
| *Rustia formosa* | Yes | Yes | No | Yes |
| *Sacoglottis guianensis* | Yes | Yes | No | Yes |
| *Sacoglottis mattogrossensis* | Yes | Yes | No | Yes |
| *Salacia arborea* | No | No | No | No |
| *Salacia elliptica* | No | Yes | No | Yes |
| *Salacia grandifolia* | Yes | Yes | No | Yes |
| *Salix humboldtiana* | No | Yes | Yes | Yes |
| *Salzmannia nitida* | No | Yes | No | No |
| *Samanea inopinata* | No | Yes | No | No |
| *Samanea tubulosa* | No | Yes | No | Yes |
| *Sambucus australis* | No | Yes | No | No |
| *Sapindus saponaria* | Yes | Yes | Yes | Yes |
| *Sapium argutum* | No | Yes | No | No |
| *Sapium glandulosum* | Yes | Yes | Yes | Yes |
| *Sapium haematospermum* | No | Yes | Yes | Yes |
| *Sapium sellowianum* | No | Yes | No | No |
| *Sarcaulus brasiliensis* | Yes | Yes | No | Yes |
| *Savia dictyocarpa* | No | Yes | No | Yes |
| *Schefflera angustisYesa* | Yes | Yes | No | Yes |
| *Schefflera calva* | Yes | Yes | No | Yes |
| *Schefflera macrocarpa* | Yes | Yes | No | Yes |
| *Schefflera morototoni* | Yes | Yes | Yes | Yes |
| *Schefflera selloi* | No | Yes | No | No |
| *Schefflera varisiana* | No | Yes | No | No |
| *Schefflera vinosa* | Yes | Yes | No | No |
| *Schinus engleri* | No | Yes | No | Yes |
| *Schinus polygamus* | No | Yes | No | Yes |
| *Schinus terebinthifolius* | Yes | Yes | No | No |
| *Schinus weinmannifolius* | No | Yes | No | No |
| *Schistostemon retusum* | No | Yes | No | Yes |
| *Schizocalyx cuspidatus* | No | Yes | No | No |
| *Schizolobium parahyba* | Yes | Yes | Yes | Yes |
| *Schoepfia brasiliensis* | No | Yes | No | No |
| *Schwartzia adamantium* | No | Yes | No | No |

| | | | | |
|---|---|---|---|---|
| *Scutia arenicola* | No | Yes | No | No |
| *Scutia buxifolia* | Yes | Yes | No | Yes |
| *Sebastiania argutidens* | No | Yes | No | No |
| *Sebastiania brasiliensis* | Yes | Yes | No | Yes |
| *Sebastiania commersoniana* | Yes | Yes | No | Yes |
| *Sebastiania edwalliana* | No | Yes | No | No |
| *Sebastiania jacobinensis* | No | Yes | No | No |
| *Sebastiania schottiana* | No | Yes | No | No |
| *Sebastiania serrata* | No | Yes | No | No |
| *Seguieria americana* | No | Yes | No | No |
| *Seguieria langsdorffii* | No | Yes | No | Yes |
| *Senefeldera verticillata* | No | Yes | No | No |
| *Senegalia lacerans* | No | Yes | No | No |
| *Senegalia langsdorffii* | No | Yes | No | No |
| *Senegalia martiusiana* | No | Yes | No | No |
| *Senegalia monacantha* | No | Yes | No | No |
| *Senegalia piauhiensis* | No | Yes | No | No |
| *Senegalia polyphylla* | No | Yes | No | Yes |
| *Senegalia recurva* | No | Yes | No | No |
| *Senegalia riparia* | No | Yes | No | No |
| *Senegalia tenuifolia* | No | Yes | No | Yes |
| *Senegalia tucumanensis* | No | Yes | No | Yes |
| *Senna affinis* | No | Yes | No | No |
| *Senna alata* | No | Yes | Yes | No |
| *Senna appendiculata* | No | Yes | No | No |
| *Senna aversiflora* | No | Yes | No | No |
| *Senna cana* | No | Yes | No | No |
| *Senna corymbosa* | No | Yes | Yes | No |
| *Senna georgica* | No | Yes | No | No |
| *Senna macranthera* | Yes | Yes | Yes | Yes |
| *Senna multijuga* | No | Yes | Yes | Yes |
| *Senna obtusifolia* | No | Yes | Yes | No |
| *Senna pendula* | No | Yes | No | No |
| *Senna pinheiroi* | No | Yes | No | No |
| *Senna quinquangulata* | No | Yes | No | No |
| *Senna rizzinii* | No | Yes | No | No |
| *Senna silvestris* | Yes | Yes | Yes | No |
| *Senna spectabilis* | No | Yes | Yes | Yes |

| | | | | |
|---|---|---|---|---|
| *Senna splendida* | No | Yes | No | No |
| *Senna velutina* | No | Yes | No | No |
| *Sessea brasiliensis* | No | Yes | No | No |
| *Sessea regnellii* | No | Yes | No | Yes |
| *Sideroxylon obtusifolium* | No | Yes | Yes | Yes |
| *Simaba cedron* | Yes | Yes | No | Yes |
| *Simaba ferruginea* | No | Yes | No | No |
| *Simaba floribunda* | No | Yes | No | No |
| *Simaba guianensis* | No | Yes | No | No |
| *Simarouba amara* | Yes | Yes | Yes | Yes |
| *Simarouba versicolor* | No | Yes | No | Yes |
| *Simira corumbensis* | No | Yes | No | No |
| *Simira glaziovii* | No | Yes | No | No |
| *Simira rubescens* | Yes | Yes | No | Yes |
| *Simira sampaioana* | No | Yes | No | No |
| *Siparuna bifida* | No | Yes | No | No |
| *Siparuna brasiliensis* | No | Yes | No | No |
| *Siparuna cymosa* | No | Yes | No | No |
| *Siparuna glycycarpa* | No | Yes | No | Yes |
| *Siparuna guianensis* | Yes | Yes | Yes | No |
| *Siparuna reginae* | No | Yes | No | No |
| *Siphoneugena crassifolia* | No | Yes | No | No |
| *Siphoneugena densiflora* | No | Yes | No | Yes |
| *Siphoneugena dussii* | No | Yes | No | No |
| *Siphoneugena reitzii* | Yes | Yes | No | No |
| *Sloanea garckeana* | Yes | Yes | No | No |
| *Sloanea guianensis* | Yes | Yes | Yes | Yes |
| *Sloanea hirsuta* | No | Yes | No | Yes |
| *Sloanea lasiocoma* | No | Yes | No | Yes |
| *Sloanea obtusifolia* | Yes | Yes | No | No |
| *Solanum alatirameum* | No | Yes | No | No |
| *Solanum argenteum* | No | Yes | No | No |
| *Solanum asperum* | Yes | Yes | No | No |
| *Solanum asterophorum* | No | Yes | No | No |
| *Solanum atropurpureum* | No | Yes | No | No |
| *Solanum bullatum* | No | Yes | No | Yes |
| *Solanum caavurana* | No | Yes | No | No |
| *Solanum campaniforme* | No | Yes | No | No |

| | | | | |
|---|---|---|---|---|
| *Solanum cernuum* | No | Yes | No | No |
| *Solanum cinnamomeum* | No | Yes | No | No |
| *Solanum compressum* | No | Yes | No | No |
| *Solanum concinnum* | No | Yes | No | No |
| *Solanum cordioides* | No | Yes | No | No |
| *Solanum corymbiflorum* | No | Yes | No | No |
| *Solanum crinitum* | No | Yes | No | Yes |
| *Solanum didymum* | No | Yes | No | No |
| *Solanum diploconos* | No | Yes | No | No |
| *Solanum evonymoides* | No | Yes | No | No |
| *Solanum granulosoleprosum* | No | Yes | No | No |
| *Solanum hexandrum* | No | Yes | No | No |
| *Solanum insidiosum* | No | Yes | No | No |
| *Solanum latiflorum* | No | Yes | No | No |
| *Solanum leucodendron* | No | Yes | No | No |
| *Solanum martii* | No | Yes | No | No |
| *Solanum mauritianum* | No | Yes | Yes | Yes |
| *Solanum melissarum* | No | Yes | No | No |
| *Solanum pabstii* | Yes | Yes | No | No |
| *Solanum paludosum* | No | Yes | No | No |
| *Solanum paniculatum* | No | Yes | No | No |
| *Solanum paranense* | No | Yes | No | No |
| *Solanum polytrichum* | No | Yes | No | No |
| *Solanum pseudoquina* | Yes | Yes | No | Yes |
| *Solanum ramulosum* | No | Yes | No | No |
| *Solanum reflexiflorum* | No | Yes | No | No |
| *Solanum restingae* | No | Yes | No | No |
| *Solanum rhytidoandrum* | No | Yes | No | No |
| *Solanum robustum* | No | Yes | No | No |
| *Solanum rufescens* | No | Yes | No | No |
| *Solanum rugosum* | No | Yes | No | No |
| *Solanum sanctae-catharinae* | No | Yes | No | No |
| *Solanum schwackei* | No | Yes | No | No |
| *Solanum scuticum* | No | Yes | No | No |
| *Solanum sooretamum* | No | Yes | No | No |
| *Solanum stipulaceum* | No | Yes | No | No |
| *Solanum stipulatum* | No | Yes | No | No |
| *Solanum stramoniifolium* | No | Yes | No | No |

| | | | | |
|---|---|---|---|---|
| *Solanum swartzianum* | No | Yes | No | No |
| *Solanum sycocarpum* | No | Yes | No | No |
| *Solanum thomasiifolium* | No | Yes | No | No |
| *Solanum trachytrichium* | No | Yes | No | No |
| *Solanum variabile* | No | Yes | No | Yes |
| *Solanum vellozianum* | No | Yes | No | No |
| *Sophora tomentosa* | No | Yes | Yes | No |
| *Sorocea bonplandii* | Yes | Yes | Yes | Yes |
| *Sorocea guilleminiana* | No | Yes | No | No |
| *Sorocea hilarii* | No | Yes | No | No |
| *Sorocea jureiana* | No | Yes | No | No |
| *Sorocea muriculata* | Yes | Yes | No | No |
| *Sorocea racemosa* | No | Yes | No | No |
| *Sparattanthelium tupiniquinorum* | No | Yes | No | No |
| *Sparattosperma leucanthum* | No | Yes | No | Yes |
| *Spirotheca rivieri* | No | Yes | No | No |
| *Spondias mombin* | Yes | Yes | Yes | Yes |
| *Spondias tuberosa* | No | Yes | No | No |
| *Spondias venulosa* | No | Yes | No | Yes |
| *Stachyarrhena harleyi* | No | Yes | No | No |
| *Stephanopodium blanchetianum* | No | Yes | No | No |
| *Sterculia apetala* | Yes | Yes | Yes | Yes |
| *Sterculia excelsa* | No | Yes | No | Yes |
| *Sterculia striata* | No | Yes | No | Yes |
| *Stifftia chrysantha* | No | Yes | No | No |
| *Stillingia oppositifolia* | No | Yes | No | No |
| *Struchium sparganophorum* | No | Yes | No | No |
| *Strychnos bahiensis* | No | Yes | No | No |
| *Strychnos brasiliensis* | No | Yes | Yes | Yes |
| *Strychnos parvifolia* | No | Yes | No | No |
| *Strychnos pseudoquina* | No | Yes | No | No |
| *Strychnos trinervis* | No | Yes | No | No |
| *Stryphnodendron adstringens* | No | Yes | No | Yes |
| *Stryphnodendron polyphyllum* | No | Yes | No | No |
| *Stryphnodendron pulcherrimum* | No | Yes | No | Yes |
| *Stylogyne lhotzkyana* | No | Yes | No | No |
| *Stylogyne pauciflora* | No | Yes | No | No |
| *Styrax acuminatus* | No | Yes | No | No |

| | | | | |
|---|---|---|---|---|
| *Styrax camporum* | Yes | Yes | No | No |
| *Styrax ferrugineus* | No | Yes | No | No |
| *Styrax glabratus* | No | No | No | No |
| *Styrax latifolius* | No | Yes | No | No |
| *Styrax leprosus* | No | Yes | No | Yes |
| *Styrax maninul* | No | Yes | No | No |
| *Styrax martii* | No | Yes | No | No |
| *Styrax pohlii* | No | Yes | No | No |
| *Swartzia acutifolia* | No | Yes | No | Yes |
| *Swartzia apetala* | No | Yes | No | No |
| *Swartzia dipetala* | No | Yes | No | No |
| *Swartzia flaemingii* | No | Yes | No | Yes |
| *Swartzia langsdorffii* | No | Yes | No | No |
| *Swartzia macrostachya* | No | Yes | No | Yes |
| *Swartzia multijuga* | Yes | Yes | No | No |
| *Swartzia myrtifolia* | No | Yes | No | Yes |
| *Swartzia oblata* | No | Yes | No | Yes |
| *Swartzia pickelii* | No | Yes | No | No |
| *Swartzia pilulifera* | No | Yes | No | No |
| *Swartzia polita* | No | Yes | No | No |
| *Swartzia polyphylla* | Yes | Yes | No | Yes |
| *Swartzia reticulata* | No | Yes | No | No |
| *Swartzia riedelii* | No | Yes | No | No |
| *Swartzia Yesplex* | Yes | Yes | Yes | Yes |
| *Sweetia fruticosa* | Yes | Yes | No | Yes |
| *Syagrus botryophora* | No | Yes | No | No |
| *Syagrus flexuosa* | No | Yes | No | No |
| *Syagrus oleracea* | No | Yes | Yes | No |
| *Syagrus pseudococos* | Yes | Yes | No | No |
| *Syagrus romanzoffiana* | No | Yes | Yes | Yes |
| *Syagrus schizophylla* | No | Yes | No | No |
| *Symphonia globulifera* | Yes | Yes | Yes | Yes |
| *Symphyopappus casarettoi* | No | Yes | No | No |
| *Symphyopappus itatiayensis* | No | Yes | No | No |
| *Symphyopappus lymansmithii* | No | Yes | No | No |
| *Symplocos celastrinea* | No | Yes | No | No |
| *Symplocos corymboclados* | No | Yes | No | No |
| *Symplocos estrellensis* | No | Yes | No | No |

| | | | | |
|---|---|---|---|---|
| *Symplocos glandulosomarginata* | No | Yes | No | No |
| *Symplocos guianensis* | No | Yes | No | No |
| *Symplocos laxiflora* | No | Yes | No | No |
| *Symplocos nitens* | No | Yes | No | No |
| *Symplocos nitidiflora* | No | Yes | No | No |
| *Symplocos oblongifolia* | No | Yes | No | No |
| *Symplocos pubescens* | No | Yes | No | No |
| *Symplocos tenuifolia* | No | Yes | No | No |
| *Symplocos tetrandra* | Yes | Yes | No | No |
| *Symplocos trachycarpos* | No | Yes | No | No |
| *Symplocos uniflora* | No | Yes | No | Yes |
| *Tabebuia aurea* | Yes | Yes | Yes | Yes |
| *Tabebuia cassinoides* | No | Yes | No | Yes |
| *Tabebuia elliptica* | No | Yes | No | Yes |
| *Tabebuia obtusifolia* | No | Yes | No | No |
| *Tabebuia roseoalba* | No | Yes | No | No |
| *Tabebuia stenocalyx* | No | Yes | No | Yes |
| *Tabernaemontana catharinensis* | Yes | Yes | No | Yes |
| *Tabernaemontana flavicans* | No | Yes | No | Yes |
| *Tabernaemontana hystrix* | No | Yes | No | No |
| *Tabernaemontana laeta* | Yes | Yes | No | Yes |
| *Tabernaemontana linkii* | No | Yes | No | No |
| *Tabernaemontana rupicola* | No | Yes | No | No |
| *Tabernaemontana salzmannii* | No | Yes | No | Yes |
| *Tachigali aurea* | No | Yes | No | No |
| *Tachigali densiflora* | No | Yes | No | No |
| *Tachigali denudata* | Yes | Yes | No | Yes |
| *Tachigali multijuga* | No | Yes | No | Yes |
| *Tachigali paratyensis* | No | Yes | No | No |
| *Tachigali pilgeriana* | No | Yes | No | No |
| *Tachigali rugosa* | Yes | Yes | No | Yes |
| *Talisia cerasina* | Yes | Yes | No | No |
| *Talisia cupularis* | Yes | Yes | No | No |
| *Talisia esculenta* | No | Yes | No | Yes |
| *Talisia macrophylla* | No | Yes | No | Yes |
| *Tapirira guianensis* | Yes | Yes | Yes | Yes |
| *Tapirira obtusa* | Yes | Yes | No | Yes |
| *Tapura amazonica* | Yes | Yes | No | No |

| | | | | |
|---|---|---|---|---|
| *Tephrosia candida* | No | Yes | Yes | No |
| *Terminalia argentea* | No | Yes | No | Yes |
| *Terminalia dichotoma* | No | Yes | No | Yes |
| *Terminalia fagifolia* | No | Yes | No | Yes |
| *Terminalia glabrescens* | No | Yes | No | No |
| *Terminalia januarensis* | Yes | Yes | No | No |
| *Terminalia mameluco* | No | Yes | No | No |
| *Terminalia phaeocarpa* | No | Yes | No | No |
| *Terminalia triflora* | Yes | Yes | Yes | Yes |
| *Ternstroemia alnifolia* | No | Yes | No | No |
| *Ternstroemia brasiliensis* | No | Yes | No | Yes |
| *Tetracera breyniana* | No | Yes | No | No |
| *Tetragastris catuaba* | No | Yes | No | No |
| *Tetrastylidium grandifolium* | No | Yes | Yes | No |
| *Tetrorchidium rubrivenium* | Yes | Yes | Yes | Yes |
| *Thyrsodium spruceanum* | Yes | Yes | No | Yes |
| *Tibouchina arborea* | No | Yes | No | Yes |
| *Tibouchina candolleana* | No | Yes | No | Yes |
| *Tibouchina clavata* | No | Yes | No | No |
| *Tibouchina elegans* | No | Yes | No | No |
| *Tibouchina estrellensis* | No | Yes | No | No |
| *Tibouchina fissinervia* | No | Yes | No | No |
| *Tibouchina fothergillae* | No | Yes | No | No |
| *Tibouchina granulosa* | No | Yes | No | No |
| *Tibouchina heteromalla* | No | Yes | No | No |
| *Tibouchina maximiliana* | No | Yes | No | No |
| *Tibouchina mosenii* | No | Yes | No | No |
| *Tibouchina mutabilis* | No | Yes | No | Yes |
| *Tibouchina pulchra* | No | Yes | No | No |
| *Tibouchina regnellii* | No | Yes | No | No |
| *Tibouchina reitzii* | No | Yes | No | No |
| *Tibouchina sellowiana* | No | Yes | No | Yes |
| *Tibouchina semidecandra* | No | Yes | No | No |
| *Tibouchina stenocarpa* | Yes | Yes | No | No |
| *Tibouchina trichopoda* | No | Yes | No | No |
| *Tibouchina urvilleana* | No | Yes | No | No |
| *Tithonia diversifolia* | No | Yes | No | No |
| *Tocoyena brasiliensis* | No | Yes | No | No |

| | | | | |
|---|---|---|---|---|
| *Tocoyena bullata* | No | Yes | No | No |
| *Tocoyena formosa* | Yes | Yes | No | No |
| *Tocoyena sellowiana* | No | Yes | No | No |
| *Tontelea attenuata* | No | Yes | No | No |
| *Tournefortia bicolor* | No | Yes | Yes | No |
| *Tournefortia candidula* | No | Yes | No | No |
| *Tournefortia paniculata* | No | Yes | No | No |
| *Tournefortia rubicunda* | No | Yes | No | No |
| *Tovomita brasiliensis* | No | Yes | No | No |
| *Tovomita brevistaminea* | No | Yes | No | No |
| *Tovomita mangle* | No | Yes | No | No |
| *Tovomitopsis paniculata* | No | Yes | No | No |
| *Tovomitopsis saldanhae* | No | Yes | No | No |
| *Trema micrantha* | Yes | Yes | Yes | Yes |
| *Trembleya parviflora* | No | Yes | No | No |
| *Trichilia casaretti* | No | Yes | No | Yes |
| *Trichilia catigua* | No | Yes | No | Yes |
| *Trichilia claussenii* | No | Yes | No | No |
| *Trichilia elegans* | No | Yes | No | No |
| *Trichilia emarginata* | No | Yes | No | No |
| *Trichilia hirta* | No | Yes | No | Yes |
| *Trichilia lepidota* | No | Yes | No | Yes |
| *Trichilia martiana* | Yes | Yes | No | Yes |
| *Trichilia pallens* | No | Yes | No | No |
| *Trichilia pallida* | Yes | Yes | Yes | Yes |
| *Trichilia pleeana* | Yes | Yes | No | Yes |
| *Trichilia pseudostipularis* | No | Yes | No | No |
| *Trichilia quadrijuga* | Yes | Yes | No | Yes |
| *Trichilia ramalhoi* | No | Yes | No | No |
| *Trichilia silvatica* | No | Yes | No | Yes |
| *Triplaris americana* | Yes | Yes | No | Yes |
| *Triplaris gardneriana* | No | Yes | No | Yes |
| *Triplaris weigeltiana* | No | Yes | No | Yes |
| *Trixis antimenorrhoea* | No | Yes | No | No |
| *Trixis divaricata* | No | Yes | No | No |
| *Trixis praestans* | No | Yes | No | No |
| *Unonopsis bahiensis* | No | Yes | No | No |
| *Unonopsis guatterioides* | Yes | Yes | No | Yes |

| | | | | |
|---|---|---|---|---|
| *Unonopsis stipitata* | No | Yes | No | Yes |
| *Urbanodendron verrucosum* | No | Yes | No | No |
| *Urera aurantiaca* | No | Yes | No | No |
| *Urera baccifera* | Yes | Yes | Yes | Yes |
| *Urera caracasana* | Yes | Yes | Yes | Yes |
| *Urera nitida* | No | Yes | No | No |
| *Vachellia farnesiana* | No | Yes | No | No |
| *Vantanea compacta* | No | Yes | No | Yes |
| *Vantanea obovata* | No | Yes | No | No |
| *Varronia curassavica* | No | Yes | No | No |
| *Varronia leucocephala* | No | Yes | No | No |
| *Varronia multispicata* | No | Yes | No | No |
| *Varronia polycephala* | No | Yes | No | No |
| *Varronia tarodaea* | No | Yes | No | No |
| *Vassobia breviflora* | Yes | Yes | Yes | Yes |
| *Vatairea heteroptera* | No | Yes | No | No |
| *Vatairea macrocarpa* | No | Yes | No | Yes |
| *Vataireopsis araroba* | No | Yes | No | Yes |
| *Verbenoxylum reitzii* | No | Yes | No | No |
| *Verbesina macrophylla* | No | Yes | No | No |
| *Vernonanthura beyrichii* | No | Yes | No | No |
| *Vernonanthura brasiliana* | No | Yes | No | No |
| *Vernonanthura discolor* | No | Yes | No | Yes |
| *Vernonanthura divaricata* | No | Yes | No | No |
| *Vernonanthura ferruginea* | No | Yes | No | No |
| *Vernonanthura petiolaris* | No | Yes | No | No |
| *Vernonanthura phosphorica* | No | Yes | No | No |
| *Vernonanthura puberula* | No | Yes | No | Yes |
| *Vernonanthura subverticillata* | No | Yes | No | No |
| *Vernonanthura tweedieana* | No | Yes | No | No |
| *Vernonanthura vinhae* | No | Yes | No | No |
| *Vernonia rubriramea* | No | Yes | No | No |
| *Virola bicuhyba* | Yes | Yes | Yes | Yes |
| *Virola gardneri* | No | Yes | Yes | Yes |
| *Virola officinalis* | No | Yes | No | No |
| *Virola sebifera* | Yes | Yes | Yes | Yes |
| *Vismia baccifera* | Yes | Yes | Yes | Yes |
| *Vismia brasiliensis* | No | Yes | No | Yes |

| | | | | |
|---|---|---|---|---|
| *Vismia guianensis* | No | Yes | No | Yes |
| *Vismia latifolia* | No | Yes | No | Yes |
| *Vismia macrophylla* | Yes | Yes | Yes | Yes |
| *Vismia magnoliifolia* | No | No | No | No |
| *Vismia martiana* | No | Yes | No | No |
| *Vitex capitata* | No | Yes | Yes | No |
| *Vitex cymosa* | No | Yes | Yes | Yes |
| *Vitex megapotamica* | No | Yes | No | Yes |
| *Vitex orinocensis* | No | Yes | No | Yes |
| *Vitex polygama* | No | Yes | No | No |
| *Vitex rufescens* | No | Yes | No | No |
| *Vitex sellowiana* | No | Yes | No | Yes |
| *Vitex triflora* | Yes | Yes | No | Yes |
| *Vochysia bifalcata* | No | Yes | No | Yes |
| *Vochysia emarginata* | No | Yes | No | No |
| *Vochysia laurifolia* | No | Yes | No | No |
| *Vochysia magnifica* | No | Yes | No | Yes |
| *Vochysia oppugnata* | No | Yes | No | No |
| *Vochysia rectiflora* | No | Yes | No | No |
| *Vochysia riedeliana* | No | Yes | No | No |
| *Vochysia thyrsoidea* | No | Yes | No | Yes |
| *Vochysia tucanorum* | No | Yes | No | Yes |
| *Waltheria cinerescens* | No | Yes | No | No |
| *Weinmannia humilis* | No | Yes | No | No |
| *Weinmannia paulliniifolia* | No | Yes | No | Yes |
| *Weinmannia pinnata* | Yes | Yes | No | No |
| *Wunderlichia mirabilis* | No | Yes | No | No |
| *Ximenia americana* | Yes | Yes | Yes | Yes |
| *Xylopia aromatica* | Yes | Yes | Yes | Yes |
| *Xylopia brasiliensis* | No | Yes | No | Yes |
| *Xylopia emarginata* | No | Yes | No | Yes |
| *Xylopia frutescens* | Yes | Yes | Yes | Yes |
| *Xylopia laevigata* | No | Yes | No | No |
| *Xylopia langsdorfiana* | No | Yes | No | No |
| *Xylopia nitida* | Yes | Yes | No | Yes |
| *Xylopia ochrantha* | No | Yes | No | No |
| *Xylopia sericea* | Yes | Yes | No | Yes |
| *Xylosma ciliatifolia* | No | Yes | No | Yes |

| | | | | |
|---|---|---|---|---|
| *Xylosma glaberrimum* | No | Yes | No | No |
| *Xylosma prockia* | No | Yes | No | No |
| *Xylosma pseudosalzmannii* | No | Yes | No | No |
| *Xylosma tweedianum* | No | No | No | No |
| *Xylosma venosa* | No | Yes | No | No |
| *Zanthoxylum acuminatum* | Yes | Yes | No | Yes |
| *Zanthoxylum caribaeum* | No | Yes | Yes | Yes |
| *Zanthoxylum fagara* | Yes | Yes | Yes | Yes |
| *Zanthoxylum gardneri* | No | Yes | No | No |
| *Zanthoxylum kleinii* | No | Yes | No | No |
| *Zanthoxylum monogynum* | No | Yes | No | Yes |
| *Zanthoxylum petiolare* | No | Yes | No | No |
| *Zanthoxylum rhoifolium* | Yes | Yes | Yes | Yes |
| *Zanthoxylum riedelianum* | No | Yes | No | Yes |
| *Zanthoxylum syncarpum* | No | Yes | No | No |
| *Zanthoxylum tingoassuiba* | No | Yes | No | No |
| *Zeyheria tuberculosa* | No | Yes | No | Yes |
| *Ziziphus joazeiro* | No | Yes | No | No |
| *Ziziphus platyphylla* | No | Yes | No | No |
| *Ziziphus undulata* | No | Yes | No | No |
| *Zollernia glabra* | No | Yes | No | No |
| *Zollernia ilicifolia* | Yes | Yes | No | Yes |
| *Zollernia modesta* | No | Yes | No | No |
| *Zollernia paraensis* | No | Yes | No | Yes |
| *Zygia latifolia* | No | Yes | No | Yes |

### *Appendix S7*

#### Species level

Trait gaps at the species level were investigated in relation to range size and economic use of wood in different combination. Here model selection was performed with minimum AIC values within $\Delta AIC \leq 2$. Models were generated with general linear models (GLM) using binomial distribution.

Table_S 4: Specific leaf area gap at the species level models with different variable combinations, Akaike's (AIC) values, degrees of freedom (df) and $\Delta AIC_i$.

| Variables | df | AIC | $\Delta AIC_i$ |
|---|---|---|---|

| | | | |
|---|---|---|---|
| range size + economic use of wood | 3 | 1700.9 | 0 |
| range size | 2 | 1747.9 | 46.9 |
| economic use of wood | 2 | 2148.7 | 447.8 |
| Only the intercept | 1 | 2212.5 | 511.5 |

Table_S 5: Maximum height gap at the species level models with different variable combinations, Akaike's (AIC) values, degrees of freedom (df) and ΔAICi.

| Variables | df | AIC | $\Delta AIC_i$ |
|---|---|---|---|
| range size | 2 | 139.3 | 0 |
| range size + economic use of wood | 3 | 140.1 | 0.85 |
| Only the intercept | 1 | 140.9 | 1.61 |
| economic use of wood | 2 | 141.2 | 1.88 |

Table_S 6: Seed dry mass gap at the species level models with different variable combinations, Akaike's (AIC) values, degrees of freedom (df) and ΔAICi.

| Variables | df | AIC | $\Delta AIC_i$ |
|---|---|---|---|
| range size + economic use of wood | 3 | 1474.5 | 0 |
| range size | 2 | 1534.2 | 59.6 |
| economic use of wood | 2 | 1617.5 | 142.9 |
| Only the intercept | 1 | 1689.6 | 215.1 |

Table_S 7: Stem specific density gap at the species level models with different variable combinations, Akaike's (AIC) values, degrees of freedom (df) and ΔAICi.

| Variables | df | AIC | $\Delta AIC_i$ |
|---|---|---|---|
| range size + economic use of wood | 3 | 2144.5 | 0 |
| economic use of wood | 2 | 2364.5 | 220.0 |
| range size | 2 | 2676.9 | 532.3 |
| Only the intercept | 1 | 2919.8 | 775.3 |

Detailed GLM outputs of are showed below.

Table_S 8: Generalized linear models (GLM) fitted values for specific leaf area (SLA), seed dry mass (SDM) and wood density (SSD) at the species level.

| | Estimate (β) | Standard Error | z value | *P* |
|---|---|---|---|---|
| **SLA trait gap** | | | | |
| (Intercept) | 2.966 | 0.112 | 26.571 | < 0.001 |
| Range size | $-3.06 \times 10^{-6}$ | $1.61 \times 10^{-7}$ | -19.043 | <0.001 |
| Economical use of wood | -0.921 | 0.13 | -7.064 | <0.001 |
| **SDM trait gap** | | | | |
| (Intercept) | 3.105 | 0.121 | 25.582 | <0.001 |
| Range size | $-1.89 \times 10^{-6}$ | $1.56 \times 10^{-7}$ | -12.164 | <0.001 |
| Economical use of wood | -1.105 | 0.139 | -7.975 | <0.001 |
| **SSD trait gap** | | | | |
| (Intercept) | 2.002 | 0.0856 | 23.38 | <0.001 |
| Range size | $-2.08 \times 10^{-6}$ | $1.48 \times 10^{-7}$ | -14.09 | <0.001 |
| Economical use of wood | -2.61 | 0.125 | -20.9 | < 0.001 |

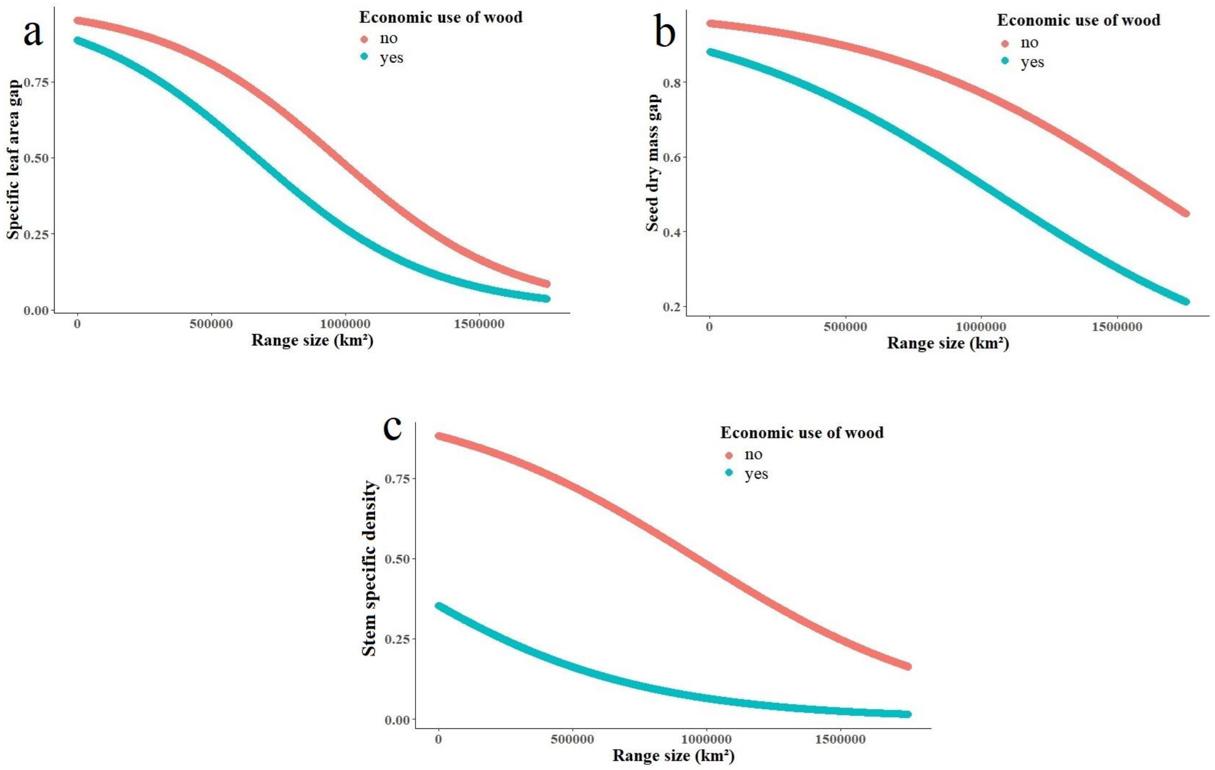


FigureS 17: Trait gaps are biased towards economic use of wood and species mean range size for a) speciefic leaf area, b) seed dry mass and c) stem specific density. Shortfalls are measured by the proportion of trait gap. Economic species (yes) are more known than species with no economic use (no). Range size is the species geographial distribution mean area.

### Geographical level

**8.1** Trait gaps at the geographical level were investigated in relation to mean range size, mean economic use of wood, distance from urban areas and distance from protected areas in different combinations. Here model selection was performed with minimum AIC values within ΔAIC≤ 2. Models were generated with general linear models (GLM) using beta distribution.

Table_S 9: Specific leaf area gap at the geographical level models with different variable combinations, Akaike's (AIC) values, degrees of freedom (df) and ΔAICi

| Variables | df | AIC | $\Delta AIC_i$ |
|---|---|---|---|
| range size + economic use of wood + distance from protected areas + distance from urban areas | 6 | -73964.8 | 0 |
| distance from urban areas + economic use of wood + range size | 5 | -73945.7 | 19.1 |
| distance from protected areas + range size + economic use of wood | 5 | -73443.9 | 520.9 |
| range size + economic use of wood | 4 | -73441.9 | 522.8 |

| | | | |
|---|---|---|---|
| distance from urban areas + distance from protected areas + range size | 5 | -68190.4 | 5774.4 |
| distance from protected areas + range size | 4 | -67966.9 | 5997.9 |
| distance from urban areas + economic use of wood | 4 | -67230.0 | 6734.8 |
| range size | 3 | -67127.8 | 6836.9 |
| distance from protected areas + distance from urban areas + economic use of wood | 5 | -52125.1 | 21839.7 |
| distance from urban areas + distance from protected areas | 4 | -52058.3 | 21906.5 |
| distance from protected areas + economic use of wood | 4 | -51907.4 | 22057.4 |
| distance from protected areas | 3 | -51845.6 | 22119.2 |
| distance from urban areas + economic use of wood | 4 | -50633.0 | 23331.8 |
| distance from urban areas | 3 | -50206.7 | 23758.1 |
| economic use of wood | 3 | -50124.1 | 23840.7 |
| only the intercept | 2 | -49652.6 | 24312.2 |

Table_S 10: Maximum height gap at the geographical level models with different variable combinations, Akaike's (AIC) values, degrees of freedom (df) and ΔAICi

| Variables | df | AIC | $\Delta AIC_i$ |
|---|---|---|---|
| range size + economic use of wood + distance from protected areas + distance from urban areas | 6 | -136510.3 | 0 |
| distance from urban areas + economic use of wood + range size | 5 | -136466.3 | 44.0 |
| distance from protected areas + range size + economic use of wood | 5 | -136443.2 | 67.2 |
| range size + economic use of wood | 4 | -136410.4 | 99.9 |
| distance from protected areas + distance from urban areas + economic use of wood | 5 | -135857.9 | 652.4 |
| distance from protected areas + economic use of wood | 4 | -135852.7 | 657.6 |
| economic use of wood | 3 | -135567.6 | 942.7 |
| distance from urban areas + economic use of wood | 4 | -135566.0 | 944.3 |

| | | | |
|---|---|---|---|
| distance from urban areas + distance from protected areas | 4 | -132402.3 | 4108.0 |
| distance from urban areas + distance from protected areas + range size | 5 | -132400.5 | 4109.8 |
| distance from protected areas | 3 | -132399.6 | 4110.7 |
| distance from protected areas + range size | 4 | -132398.5 | 4111.8 |
| range size | 3 | -131472.7 | 5037.6 |
| distance from urban areas + economic use of wood | 4 | -131471.7 | 5038.6 |
| distance from urban areas | 3 | -131433.5 | 5076.8 |
| only the intercept | 2 | -131429.9 | 5080.5 |

Table_S 11: Seed dry mass gap at the geographical level models with different variable combinations, Akaike's (AIC) values, degrees of freedom (df) and ΔAICi

| Variables | df | AIC | $\Delta AIC_i$ |
|---|---|---|---|
| range size + economic use of wood + distance from protected areas + distance from urban areas | 6 | -70928.7 | 0 |
| distance from urban areas + economic use of wood + range size | 5 | -70818.6 | 110.1 |
| distance from protected areas + range size + economic use of wood | 5 | -70732.4 | 196.2 |
| range size + economic use of wood | 4 | -70653.4 | 275.3 |
| distance from protected areas + distance from urban areas + economic use of wood | 5 | -61235.2 | 9693.4 |
| distance from protected areas + economic use of wood | 4 | -61143.3 | 9785.4 |
| distance from urban areas + economic use of wood | 4 | -59894.7 | 11033.9 |
| economic use of wood | 3 | -59600.7 | 11328.0 |
| distance from urban areas + distance from protected areas + range size | 5 | -53269.0 | 17659.7 |
| distance from protected areas + range size | 4 | -53265.0 | 17663.6 |
| distance from urban areas + distance from protected areas | 4 | -52205.7 | 18722.9 |
| distance from protected areas | 3 | -52181.3 | 18747.4 |
| distance from urban areas + economic use of wood | 4 | -51328.6 | 19600.0 |

| | | | |
|---|---|---|---|
| range size | 3 | -51310.8 | 19617.9 |
| distance from urban areas | 3 | -49515.2 | 21413.5 |
| only the intercept | 2 | -49232.9 | 21695.7 |

Table_S 12: Stem specific density gap at the geographical level models with different variable combinations, Akaike's (AIC) values, degrees of freedom (df) and ΔAICi

| Variables | df | AIC | $\Delta AIC_i$ |
|---|---|---|---|
| range size + economic use of wood + distance from protected areas + distance from urban areas | 6 | -66972.7 | 0 |
| distance from urban areas + economic use of wood + range size | 5 | -66665.0 | 307.6 |
| distance from protected areas + range size + economic use of wood | 5 | -66665.0 | 307.6 |
| range size + economic use of wood | 4 | -66657.9 | 314.8 |
| distance from protected areas + distance from urban areas + economic use of wood | 5 | -58338.0 | 8634.6 |
| distance from protected areas + economic use of wood | 4 | -58302.0 | 8670.6 |
| distance from urban areas + economic use of wood | 4 | -57311.2 | 9661.5 |
| economic use of wood | 3 | -57145.0 | 9827.7 |
| distance from urban areas + distance from protected areas + range size | 5 | -47693.3 | 19279.4 |
| distance from protected areas + range size | 4 | -47688.5 | 19284.2 |
| distance from urban areas + distance from protected areas | 4 | -47059.5 | 19913.1 |
| distance from protected areas | 3 | -47051.1 | 19921.6 |
| distance from urban areas + economic use of wood | 4 | -45870.0 | 21102.7 |
| range size | 3 | -45856.8 | 21115.9 |
| distance from urban areas | 3 | -44651.1 | 22321.6 |
| only the intercept | 2 | -44462.1 | 22510.6 |

Table_S 13: Generalized linear models (GLM) fitted values for specific leaf area (SLA), seed dry mass (SDM) and wood density (SSD) at the species level.

| | Estimate (β) | Standard Error | z value | *P* |
|---|---|---|---|---|
| **SLA trait gap** | | | | |
| (Intercept) | 2.966 | 0.112 | 26.571 | < 0.001 |
| Range size | $-3.06 \times 10^{-6}$ | $1.61 \times 10^{-7}$ | -19.043 | <0.001 |
| Economical use of wood | -0.921 | 0.13 | -7.064 | <0.001 |
| **SDM trait gap** | | | | |
| (Intercept) | 3.105 | 0.121 | 25.582 | <0.001 |
| Range size | $-1.89 \times 10^{-6}$ | $1.56 \times 10^{-7}$ | -12.164 | <0.001 |
| Economical use of wood | -1.105 | 0.139 | -7.975 | <0.001 |
| **SSD trait gap** | | | | |
| (Intercept) | 2.002 | 0.0856 | 23.38 | <0.001 |
| Range size | $-2.08 \times 10^{-6}$ | $1.48 \times 10^{-7}$ | -14.09 | <0.001 |
| Economical use of wood | -2.61 | 0.125 | -20.9 | < 0.001 |

Table_S 14: Generalized linear models (GLM) fitted values for specific leaf area (SLA), seed dry mass (SDM) and wood density (WD) at the geographical level.

| | Estimate | Standard Error | z value | *P* |
|---|---|---|---|---|
| **SLA trait gap** | | | | |
| (Intercept) | 1.947 | $4.97 \times 10^{-3}$ | 391.836 | <0.001 |
| Distance from urban areas | $5.82 \times 10^{-2}$ | $2.54 \times 10^{-3}$ | 22.922 | <0.001 |
| Economical use of wood | -0.80 | $9.34 \times 10^{-3}$ | -85.779 | <0.001 |
| Distance from protected areas | -0.0157 | $3.42 \times 10^{-3}$ | -4.596 | <0.001 |
| Range size | $-1.61 \times 10^{-6}$ | $6.86 \times 10^{-9}$ | -235.17 | <0.001 |
| **SDM trait gap** | | | | |
| (Intercept) | 2.740 | $6.85 \times 10^{-3}$ | 399.88 | <0.001 |

| | | | | |
|---|---|---|---|---|
| Distance from urban areas | 0.049 | $3.49 \times 10^{-3}$ | 14.01 | <0.001 |
| Economical use of wood | -2.46 | 0.0128 | -192.78 | <0.001 |
| Distance from protected areas | -0.049 | $4.60 \times 10^{-3}$ | -10.67 | <0.001 |
| Range size | $-1.12 \times 10^{-6}$ | $9.30 \times 10^{-9}$ | -120.50 | <0.001 |
| **WD trait gap** | | | | |
| (Intercept) | 1.256 | $5.30 \times 10^{-3}$ | 236.81 | <0.001 |
| Distance from urban areas | 0.048 | $2.73 \times 10^{-3}$ | 17.685 | <0.001 |
| Economical use of wood | -2.052 | $9.96 \times 10^{-3}$ | -206.14 | <0.001 |
| Distance from protected areas | -0.019 | $3.70 \times 10^{-3}$ | -5.108 | <0.001 |
| Range size | $-8.12 \times 10^{-7}$ | $7.39 \times 10^{-9}$ | -109.85 | <0.001 |

**8.2.** Trait dominance (CWM) of SLA ranged from 83.25 to 225.04 $cm^2.g$-1, while trait dispersion (Q) ranged from 0.17 to 0.42. CWM of HMAX ranged from 11.75 to 19.51 m, while Q ranged from 0.25 to 0.36. CWM of SDM ranged from 0.089 to 3.92 g, while Q ranged from 0 to 0.19. CWM of SSD ranged from 0.470 to 0.742 $g.cm^{-3}$, while Q ranged from 0.30 to 0.42.
